# The health-relevant architecture of the everyday light exposome

**DOI:** 10.64898/2026.06.04.730226

**Authors:** Johannes Zauner, Altug Didikoglu, Sam Aerts, Gabriel Kwaku Agbeshie, Kwadwo Owusu Akuffo, Sena Gulsum Akgun, Sema Nur Aydin, David Baeza Moyano, Daan Boesten, John F.B. Bolte, Kai Broszio, Guadalupe Cantarero García, Roberto Alonso González Lezcano, Carolina Guidolin, Sarina Hilden, Nico Hogervorst, Astrid Jansen, Zeynep Kayar, Stefan Källberg, Suyoun Lee, Sofía Melero Tur, Maria Nilsson Tengelin, María Concepción Pérez Gutiérrez, Andrea Sancho-Salas, Oliver Stefani, Ingemar Svensson, Helga von-Breymann, Manuel Spitschan

## Abstract

Light supports circadian regulation and is associated with non-communicable diseases, yet ocular exposure patterns remain poorly understood. We recruited 191 adults across nine sites in seven countries, combining diaries and contextual reports with near-eye (141 participants; 816 participant-days) and complementary chest-level measurements (154 participants; 902 participant-days). In near-eye analyses, only 24.0% of recorded daytime minutes met the recommendation of at least 250 lx melanopic equivalent daylight illuminance; 63.3% of pre-sleep and 87.7% of bedside sleep-environment minutes met their respective limits. Variation among people and days within sites exceeded that among sites (ratio 1.99; 95%-CI, 1.29–4.77). In hourly models, light source and setting contributed the largest shares of fitted variation after time of day. Site-average outdoor exposure was 9.35 times that while awake at home (95%-CI, 6.95–12.59; 714.2 versus 76.4 lx). This baseline identifies a daytime exposure gap and supports testing interventions in everyday settings to improve light exposure and health.

## Introduction

Light reaching the eye supports vision and signals circadian timing, sleep, alertness and neuroendocrine responses^1–5^. Melanopic equivalent daylight illuminance (melanopic EDI) weights illuminance for melanopsin-related sensitivity and is most relevant to these effects^6,7^. Expert consensus recommends at least 250 lx melanopic EDI at the eye during daytime, no more than 10 lx during the three hours before sleep and no more than 1 lx in the sleep environment for healthy adults with regular daytime schedules^8^. These physiology-based ranges make daily light a health-relevant exposure, but are not individual risk functions.

Population studies and reviews associate personal light patterns with sleep and major non-communicable diseases^9^. Wrist-sensor studies link light timing and day-night distribution with sleep timing and sleepiness, psychiatric disorders, incident type 2 diabetes, cardiovascular disease and mortality^10–15^. Other cohorts link evening, nocturnal or seasonal light with sleep in bipolar disorder and with obesity, diabetes, hypertension and metabolic health^16–19^. The exposome framework characterises non-genetic exposures across the life course^20,21^, yet rarely includes time-resolved ocular melanopic light, the everyday component examined here. These studies establish population relevance but mostly measured broad-spectrum wrist or bedside light, providing only proxy evidence for melanopic ocular exposure.

An everyday exposure baseline against these recommendations is missing. Personal exposure to natural and electric light follows an expected 24-hour rhythm, including in large wrist-sensor cohorts^22^, but previous studies focus on particular populations, occupations or settings^23–25^. Its amplitude and timing across sites and countries, and the contributions of time, site, individuals, day-to-day variation and immediate context, remain uncertain. Daylight availability, satellite estimates and fixed-site measurements cannot resolve these layers as people move through buildings, transport, outdoor spaces and sleep environments^26^. Sensor position also matters: near-eye sensing is closer to the incident ocular field but is not a retinal measurement; wrist and chest sensors capture related fields shaped by posture, wear and occlusion^27–33^.

Using the harmonised MeLiDos protocol, we measured personal light across nine sites in seven countries^34,35^. The near-eye sensor characterised ocular light exposure during wear, alongside diaries, light-source and immediate-setting reports, sleep information and questionnaires. The complementary chest sensor provided non-ocular environmental evidence and enabled participation among people reluctant to wear the glasses-mounted sensor. We sought a multisite recommendation-based baseline, assessing temporal, environmental and routine contributions alongside selected person-level correlates. We use architecture to describe how everyday light exposure is organised across time, sites, people, days and immediate settings, and benchmark this component of the exposome against health-based light recommendations. Although not global, the protocol enables harmonised measurement elsewhere, progressively closing gaps across locations, civil photoperiods, latitudes and climates and building towards a global account of ocular light exposure.

## Results

### A multisite baseline and recommendation adherence

The normalised roster comprised 191 participants across Borås (SE), Delft (NL), Dortmund (DE), Tübingen (DE), Munich (DE), Madrid (ES), Izmir (TR), San José (CR) and Kumasi (GH). Of these, 184 contributed 1,478 participant-days with recorded, non-all-zero light data at either position before quality screening. After screening, the near-eye dataset contained 141 participants and 816 participant-days, the complementary chest dataset 154 and 902, and the common sample 112 participants and 643 participant-days. Near-eye measurements provide ocular-exposure evidence during wear, whereas chest measurements are non-ocular; both describe the bedside sleep environment during reported sleep. Supplementary Table S1 gives the sample flow.

The pooled near-eye record followed the expected 24-hour rhythm: low overnight exposure, a morning rise, higher daytime values and an evening decline. Seventeen complementary metrics captured thematic streams rather than one score. Level metrics described daily geometric mean melanopic EDI and the brightest and darkest supported 10-hour means; duration and timing metrics quantified exposure ranges, sustained bright periods, bright or dark windows and threshold crossings.

Dose integrated intensity, regularity captured stability and fragmentation, and melanopic daylight efficacy (MDER) described spectral composition relative to visual illuminance. These features are not interchangeable health-response estimates. Supplementary Table S2. and Supplementary Figure S1 give the metric dictionary and site distributions; Supplementary Figure S2 shows a worked derivation. Figure 1 and Table 1 summarise the design and data structure.

**Figure 1:**
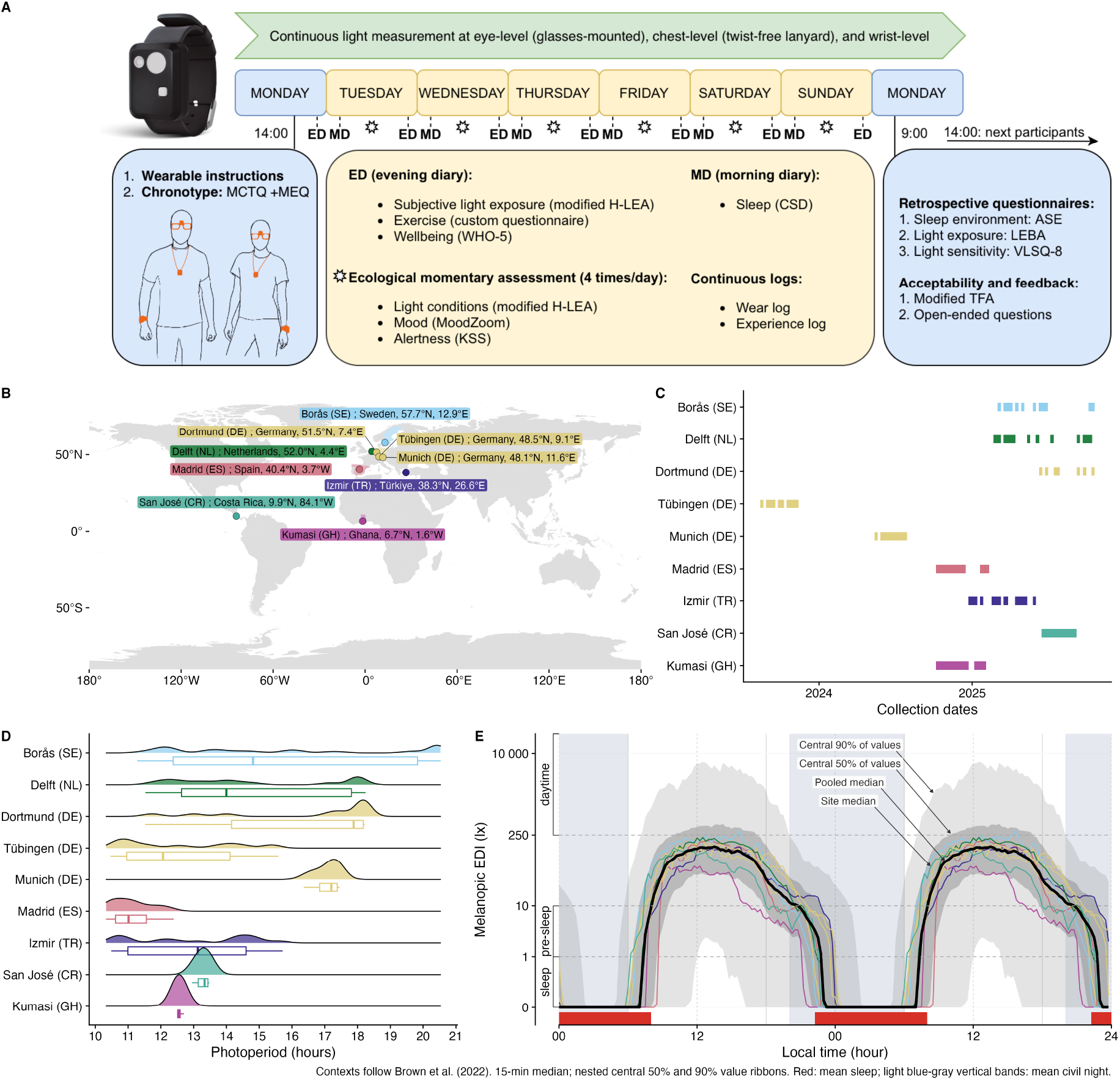
Five-panel overview of the eight-day protocol, sensor positions, nine country-coded sites, collection windows, civil photoperiod, and the pooled near-eye daily profile. **A**, Eight-day field protocol and sensor positions. The wrist logger is shown as a protocol device but is not an exposure channel analysed in this manuscript. **B**, Nine sites in seven countries. **C**, Site collection periods. **D**, Observed civil photoperiod. **E**, Site and pooled 15-minute median near-eye melanopic equivalent daylight illuminance profiles from 816 participant-days contributed by 141 participants, repeated over 48 hours for display only. Grey ribbons show the central 50% and 90% of measured values, not confidence intervals; coloured lines are site medians, the black line is the pooled median, and red and blue shading show mean reported sleep and civil night (sun elevation < −6°). During reported sleep, the measurement represents the bedside sleep environment.

**Table 1:** Participant and measurement characteristics overall and by country-coded study site. The table distinguishes the 191-person roster, participants and participant-days with recorded light data, and the screened near-eye and complementary chest samples. Model-specific samples are reported with each analysis. Of the 191 roster participants, 184 contributed the 1,478 recorded participant-days. Continuous participant characteristics are median with the stated percentile range.

|  | Overall | Borås (SE) | Delft (NL) | Dortmund (DE) | Tübingen (DE) | Munich (DE) | Madrid (ES) | Izmir (TR) | San José (CR) | Kumasi (GH) |
| --- | --- | --- | --- | --- | --- | --- | --- | --- | --- | --- |
| **Site specifics** |  |  |  |  |  |  |  |  |  |  |
| Institution | Overall | RISE | THUAS | BAuA | MPI | TUM | FUSPCEU | IZTECH | UCR | KNUST |
| Country | Not applicable | Sweden | Netherlands | Germany | Germany | Germany | Spain | Türkiye | Costa Rica | Ghana |
| City | Not applicable | Borås | Delft | Dortmund | Tübingen | Munich | Madrid | Izmir | San José | Kumasi |
| Coordinates | Not applicable | 57.7°N, 12.9°E | 52.0°N, 4.4°E | 51.5°N, 7.4°E | 48.5°N, 9.1°E | 48.1°N, 11.6°E | 40.4°N, 3.7°W | 38.3°N, 26.6°E | 9.9°N, 84.1°W | 6.7°N, 1.6°W |
| **Data collection** |  |  |  |  |  |  |  |  |  |  |
| Participants<sup>1</sup> | 191 roster<br>(g:143; c:157;<br>p:116) | 17 roster<br>(g:14; c:17;<br>p:14) | 20 roster<br>(g:13; c:15;<br>p:13) | 24 roster<br>(g:19; c:22;<br>p:19) | 26 roster<br>(g:26; c:0; p:0) | 10 roster<br>(g:10; c:10;<br>p:10) | 23 roster<br>(g:23; c:22;<br>p:22) | 17 roster<br>(g:17; c:17;<br>p:17) | 39 roster<br>(g:6; c:39; p:6) | 15 roster<br>(g:15; c:15;<br>p:15) |
| Participant-days<sup>1</sup> | 1478 roster<br>(g:1134<br>c:1246; p:902) | 137 roster<br>(g:107<br>c:137; p:107) | 125 roster<br>(g:107<br>c:124; p:106) | 176 roster<br>(g:145<br>c:163; p:132) | 208 roster<br>(g:208<br>c:0; p:0) | 80 roster<br>(g:80<br>c:80; p:80) | 182 roster<br>(g:182<br>c:174; p:174) | 138 roster<br>(g:138<br>c:138; p:138) | 312 roster<br>(g:48<br>c:312; p:48) | 120 roster<br>(g:119<br>c:118; p:117) |
| Participant time<sup>2</sup> | 116w 4d | 11w 23h | 11w 23h | 15w 2d | 21w 3d | 8w 4d | 18w 3d | 14w 3d | 4w 4d | 11w 4d |
| Declared non-wear<sup>2</sup> | **g:4w 1d**<br>**(3.66%)** | **g:2d 14h**<br>**(3.34%)** | **g:3d 1h**<br>**(3.92%)** | **g:3d 15h**<br>**(3.42%)** | **g:6d 10h**<br>**(4.30%)** | **g:2d 17h**<br>**(4.55%)** | **g:2d 20h**<br>**(2.20%)** | **g:3d 3h**<br>**(3.13%)** | **g:1d 11h**<br>**(4.66%)** | **g:3d 21h**<br>**(4.80%)** |
| Screened days<sup>1,3</sup> | **g:816**<br>**c:902** | **g:78**<br>**c:96** | **g:78**<br>**c:93** | **g:107**<br>**c:114** | **g:150**<br>**c:0** | **g:60**<br>**c:60** | **g:129**<br>**c:123** | **g:101**<br>**c:102** | **g:32**<br>**c:230** | **g:81**<br>**c:84** |
| Civil photoperiod<sup>4,2</sup> | 13.25 h<br>(10.62–18.20)<br>(n=1478 d) | 14.81 h<br>(11.80–20.49)<br>(n=137 d) | 14.00 h<br>(11.88–18.09)<br>(n=125 d) | 17.88 h<br>(11.76–18.22)<br>(n=176 d) | 12.07 h<br>(10.55–15.41)<br>(n=208 d) | 17.20 h<br>(16.45–17.45)<br>(n=80 d) | 11.01 h<br>(10.35–12.22)<br>(n=182 d) | 13.13 h<br>(10.52–15.57)<br>(n=138 d) | 13.34 h<br>(13.01–13.47)<br>(n=312 d) | 12.54 h<br>(12.49–12.68)<br>(n=120 d) |
| **Participant information** |  |  |  |  |  |  |  |  |  |  |
| Age<sup>4</sup> | 28 y (21 y–52 y)<br>(n=191 N) | 38 y (22.6 y–64.8 y)<br>(n=17 N) | 30.5 y (19.9 y–57.1 y)<br>(n=20 N) | 35.5 y (20.1 y–58.5 y)<br>(n=24 N) | 27 y (22 y–37 y)<br>(n=26 N) | 27.5 y (22.2 y–29 y)<br>(n=10 N) | 31 y (20.2 y–54.3 y)<br>(n=23 N) | 24 y (21 y–29.6 y)<br>(n=17 N) | 34 y (22 y–48.1 y)<br>(n=39 N) | 23 y (20.1 y–25 y)<br>(n=15 N) |
| Sex<sup>5</sup> | F:105 / M:86<br>(n=191) | F:6 / M:11<br>(n=17) | F:8 / M:12<br>(n=20) | F:13 / M:11<br>(n=24) | F:14 / M:12<br>(n=26) | F:6 / M:4<br>(n=10) | F:15 / M:8<br>(n=23) | F:11 / M:6<br>(n=17) | F:24 / M:15<br>(n=39) | F:8 / M:7<br>(n=15) |
| Employment status<sup>6</sup> | F:163 / P:26 / N:2<br>(n=191) | F:16 / P:0 / N:1<br>(n=17) | F:11 / P:9 / N:0<br>(n=20) | F:21 / P:3 / N:0<br>(n=24) | F:23 / P:3 / N:0<br>(n=26) | F:9 / P:1 / N:0<br>(n=10) | F:21 / P:2 / N:0<br>(n=23) | F:15 / P:2 / N:0<br>(n=17) | F:34 / P:5 / N:0<br>(n=39) | F:13 / P:1 / N:1<br>(n=15) |
| Chronotype group<sup>7</sup> | m:62 / i:94 / e:30<br>(n=186) | m:8 / i:4 / e:5<br>(n=17) | m:3 / i:9 / e:3<br>(n=15) | m:8 / i:13 / e:3<br>(n=24) | m:9 / i:9 / e:8<br>(n=26) | m:0 / i:8 / e:2<br>(n=10) | m:5 / i:17 / e:1<br>(n=23) | m:2 / i:10 / e:5<br>(n=17) | m:16 / i:20 / e:3<br>(n=39) | m:11 / i:4 / e:0<br>(n=15) |
| Sleep-corrected midsleep on free days<sup>4,8</sup> | 04:03 (02:08–06:15)<br>(n=185 N) | 02:55 (02:04–05:09)<br>(n=17 N) | 04:31 (03:05–05:37)<br>(n=15 N) | 03:47 (02:17–04:49)<br>(n=24 N) | 04:33 (02:41–06:04)<br>(n=26 N) | 05:08 (04:12–06:04)<br>(n=10 N) | 04:48 (03:10–06:52)<br>(n=22 N) | 05:29 (03:35–06:49)<br>(n=17 N) | 03:42 (02:17–05:31)<br>(n=39 N) | 02:19 (01:22–03:26)<br>(n=15 N) |

<sup>1</sup>g, near-eye glasses position; c, complementary chest position; p, paired common sample in which the same participant-day is available at both positions.
<sup>2</sup>w: weeks; d: days; h: hours; min: minutes. Participant time and declared non-wear use eligible real minutes on screened days; both are shown for near eye only.
<sup>3</sup>Screened days are the final participant-days after the 80% completeness screen and exclusion of exact-all-zero melEDI days.
<sup>4</sup>Median (5th percentile, 95th percentile).
<sup>5</sup>Sex categories: Female and Male.
<sup>6</sup>Employment categories: Full/studying, Part/marginal, and Not employed.
<sup>7</sup>Chronotype groups follow the Morningness–Eveningness Questionnaire score.
<sup>8</sup>MCTQ: Munich Chronotype Questionnaire; MEQ: Morningness–Eveningness Questionnaire. Clock summaries are circular.

Exposure was within the applicable recommendation during 24.0% of daytime minutes, 63.3% of pre-sleep minutes and 87.7% of sleep minutes. This represented 137,792 of 573,712 waking daytime minutes, 81,894 of 129,390 pre-sleep minutes and 336,052 of 383,366 minutes during reported sleep. The sleep quantity describes the bedside sleep environment because the device was not worn at the eye; the Discussion addresses low-light measurement accuracy. Supplementary Table S3 gives exact numerators, denominators and complementary ranges.

These pooled-minute percentages describe how often exposure lay within range, not participant adherence. Model-based comparisons retained each window’s adherent and valid-minute counts.

The adherence model included 2,298 periods from 140 participants and 794 cycles (1,043,192 valid minutes), linking preceding sleep, daytime and following pre-sleep to the wake-start date’s day type; windows qualified independently. Free-versus-work adherence was 5.1 percentage points lower during daytime (95% confidence interval [95% CI] 2.4 to 7.8 lower), 5.9 higher pre-sleep (1.4 to 10.5) and 6.5 lower during sleep (3.8 to 9.1). At least 80% coverage preserved daytime and sleep interval conclusions, but pre-sleep was inconclusive (4.1 points, 95% CI −0.9 to 9.1). Temporal dependence remained unresolved.

The window-by-site-by-day-type interaction was retained (FDR-adjusted *p* = 0.008), with site variation during daytime and sleep (both *p* = 0.002), not detected pre-sleep (*p* = 0.818). Relative to the site-average free-minus-work contrast, Dortmund (DE) was 14.8 points higher during daytime (95% CI 6.7 to 22.9; FDR-adjusted *p* = 0.004), Madrid (ES) 9.2 lower (−15.6 to −2.8; *p* = 0.044), and Kumasi (GH) 6.3 higher during sleep (3.5 to 9.1; *p* < 0.001). These departures met a separate 27-test FDR criterion and retained directions and interval conclusions at 80% coverage.

On the design-standardized response scale, marginal R² was 58.0% and conditional R² 60.2%; the participant increment was 2.3 percentage points and observation/distribution variation 39.8%. Shapley allocated 94.2% of marginal R² to recommendation window, 4.8% to site and 1.0% to day type. Site’s 2.8-point allocation was about 1.2 times the participant-intercept increment and about five times the 0.6-point day-type allocation. Allocations are point-only; the window share partly reflects different thresholds. Table 2 and Supplementary Figures S4 and S5 give adherence estimates.

**Table 2:**
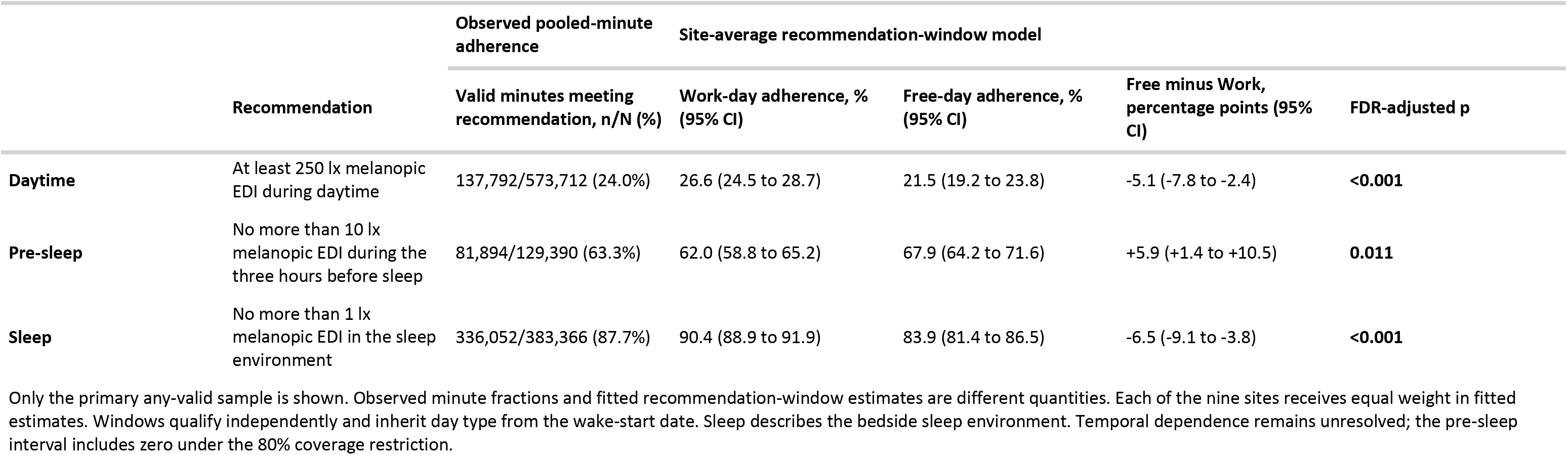
Recommendation adherence by Brown et al. recommendation window and day type. Observed valid-minute fractions are descriptive; work-day and free-day estimates and their difference are model-based, with 95% confidence intervals and false-discovery-rate-adjusted p-values. Each of nine sites receives equal weight in the model estimates. Daytime, Pre-sleep and Sleep identify the recommendation windows: daytime excludes the three hours before reported sleep, pre-sleep comprises those three hours, and sleep describes the bedside sleep environment. The wake-start date assigns the day type to the preceding sleep, daytime and following pre-sleep windows; each window qualifies independently. Only the primary any-valid sample is shown: 2,298 periods, 140 participants, 794 cycles and 1,043,192 valid minutes. The 80% coverage sensitivity remains reported in the Results and Methods; its pre-sleep interval includes zero, and temporal dependence remains unresolved. Confidence-interval exclusion and FDR decisions are separate summaries. Descriptive pooled-minute fractions are detailed in Supplementary Table S3. Adherence does not classify participants.

Exploratory cross-window within-participant contrasts per 10-percentage-point difference in daytime adherence were −0.26 points for sleep (95% CI −0.95 to 0.43) and −1.09 for pre-sleep (−2.75 to 0.56). Unresolved temporal dependence precluded a within-participant claim; intervals including zero do not establish absence. Between participants, 10 points higher average daytime adherence was associated with 2.52 points lower sleep adherence (1.48 to 3.55 lower; FDR-adjusted *p* < 0.001) and 3.59 lower pre-sleep adherence (1.25 to 5.94 lower; *p* = 0.005), consistent with higher evening and nighttime exposure during monitoring. Both met the four-effect FDR criterion; directions and interval conclusions persisted at 80% coverage (maximum change 0.60 points). Supplementary Table S4 gives estimates and Supplementary Figure S6 participant profiles.

### Personal and day-to-day differences exceed site differences

A nonlinear model decomposed personal light exposure into common local-clock, site and participant curves, and participant-day shifts. Near-eye in-sample R² was 0.775. Participant curves were 1.80 times as dispersed as site curves (95% CI 1.16 to 4.33); participant curves plus day shifts had a dispersion ratio of 1.99 (1.29 to 4.77). After accounting for time of day, fitted variation among people and their days within sites exceeded variation among sites.

Shares of full-model R² showed the same hierarchy: 78.9% for the shared local-clock pattern, 12.9% for participant patterns, 6.2% for participant-day shifts and 2.0% for site patterns. The participant-pattern share was 6.41 times the site-pattern share (3.85 to 15.14), and the combined participant-pattern and participant-day-shift share was 9.49 times the site-pattern share (5.77 to 21.81).

The shared rhythm did not imply uniform amplitude. Relative to the fitted site-average daily pattern, Borås (SE) ratios were 2.04 to 4.64 from 06:00 to 11:00 and 1.80 to 2.73 from 15:00 to 19:00. Madrid (ES) ratios were 0.15 to 0.47 from 05:30 to 10:30 and 0.40 to 0.57 from 17:30 to 20:30. Izmir (TR) shifted from 0.51 to 0.59 at 06:00 to 08:30 to 1.98 to 4.28 at 21:00 to 24:00, while Kumasi (GH) was 0.11 to 0.38 from 11:30 to 23:00. These are clock-specific departures from the average curve.

The chest model had an in-sample R² of 0.737. Of this full-model R², common time accounted for 79.1%, participant patterns 11.9%, participant-day shifts 7.0% and site patterns 2.1%. Common-sample comparisons supported the combined participant-plus-day ordering, but the participant-only contrast was less stable. Cross-position similarity is complementary context, not replication, equivalence or interchangeability. Figure 2 shows the fitted layers; Supplementary Tables S5 and S6 give both decompositions.

**Figure 2:**
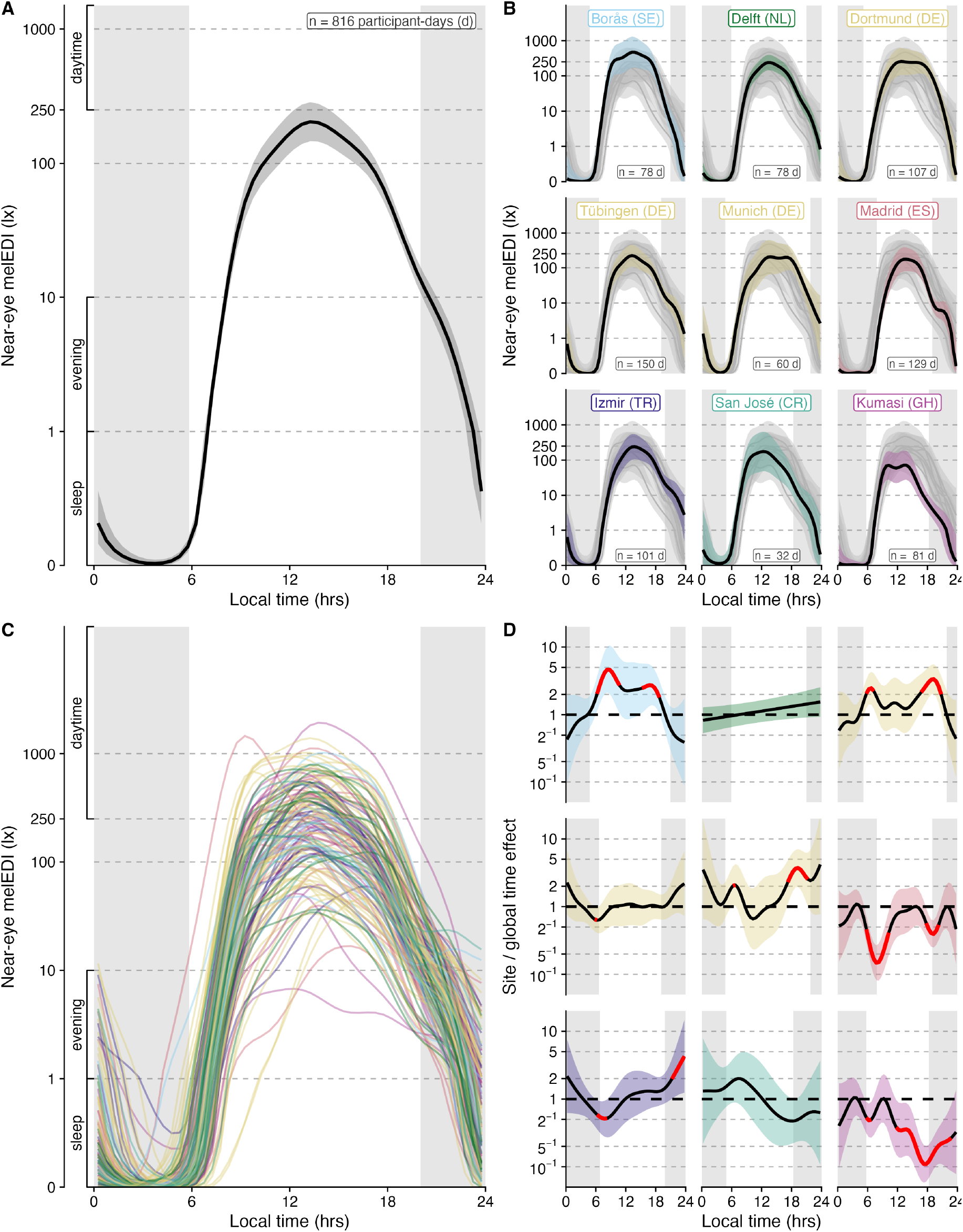
Multiscale daily pattern of near-eye melanopic equivalent daylight illuminance. **A**, Site-average daily curve. **B**, Fitted site curves. **C**, Fitted participant curves after the shared local-clock and site patterns were considered. **D**, Ratio of each fitted site curve to the site-average curve on the melanopic equivalent daylight illuminance plus 0.1 lx scale. Grey bands show mean civil night. Shaded bands are clock-specific 95% confidence intervals; red segments identify displayed half-hours whose interval did not include the site-average curve and are descriptive clock-specific contrasts. The analysis included 141 participants and 816 participant-days across nine sites. The label “global time effect” denotes the site-average local-clock curve (panel **A**).

### Geography and photoperiod provide partial context

Site associations were stream-specific. Site differences met the FDR criterion for 10 of 17 near-eye metrics: all three level metrics, time above 250 lx melEDI during wake, time below 10 lx melEDI before sleep, midpoint of the brightest 10 hours, midpoint of the darkest 10 hours, mean timing of exposure above 250 lx melEDI, last light timing above 250 lx melEDI, and MDER. They were not FDR-retained for interdaily stability, intradaily variability, time above 1,000 lx melEDI, time below 1 lx melEDI during sleep, longest continuous period above 250 lx melEDI, first light timing above 250 lx melEDI, and melEDI dose. Across metric models, conditional R² averaged 37.8%; participant-associated R² averaged 26.1% among the 15 participant-day metrics. Among the 10 metrics with site support, site-associated part R² averaged 8.7%. These non-additive components reinforce the stronger individual-level contribution in the daily-pattern analysis.

Civil-photoperiod associations were selective. Per additional hour, the fitted daily geometric mean was 1.18 times higher (95% CI 1.11 to 1.26), the brightest-10-hour mean 1.24 times higher (1.13 to 1.36), waking time above 250 lx 1.14 times higher (1.07 to 1.20) and daily dose 1.28 times higher (1.17 to 1.39). Each hour was also associated with 0.134 h less time below 10 lx before sleep (0.066 to 0.201 h less) and the last exposure above 250 lx occurring 0.323 h later (0.186 to 0.460 h). Civil photoperiod met the FDR criterion for 12 of 17 metrics but not interdaily stability, intradaily variability, time below 1 lx melEDI during sleep, midpoint of the brightest 10 hours, and first light timing above 250 lx melEDI. Among supported models, photoperiod-associated part R² averaged 5.9%; the corresponding mean for the 7 supported latitude models was 3.8%. These averages describe different supported metric sets and do not compare independent or causal contributions.

Across the recorded 10.33 to 20.52 h of civil photoperiod, six of nine level, exposure-history, and duration-based metrics changed from increasing to a sustained near-flat tail between 14.24 and 16.20 hours of photoperiod: the daily mean, both 10-hour window means, time above 1,000 lx, waking time above 250 lx and dose. Pre-sleep time below 10 lx, sleep-environment time below 1 lx and the longest period above 250 lx did not. This transition means that the effect of photoperiod becomes uncertain around 14 to 16 hours. The fitted slope became compatible with zero and remained so through the observed maximum. Some classifications were sensitive to site, sample or model form. Several limitations apply as discussed below: latitude was confounded with site, collection periods and photoperiod overlap differed, and weather, built environment, culture and routine could not be separated. The metric-specific results are summarised in Table 3, Supplementary Figures S7 and S8, and Supplementary Tables S7 and S8.

**Table 3:**
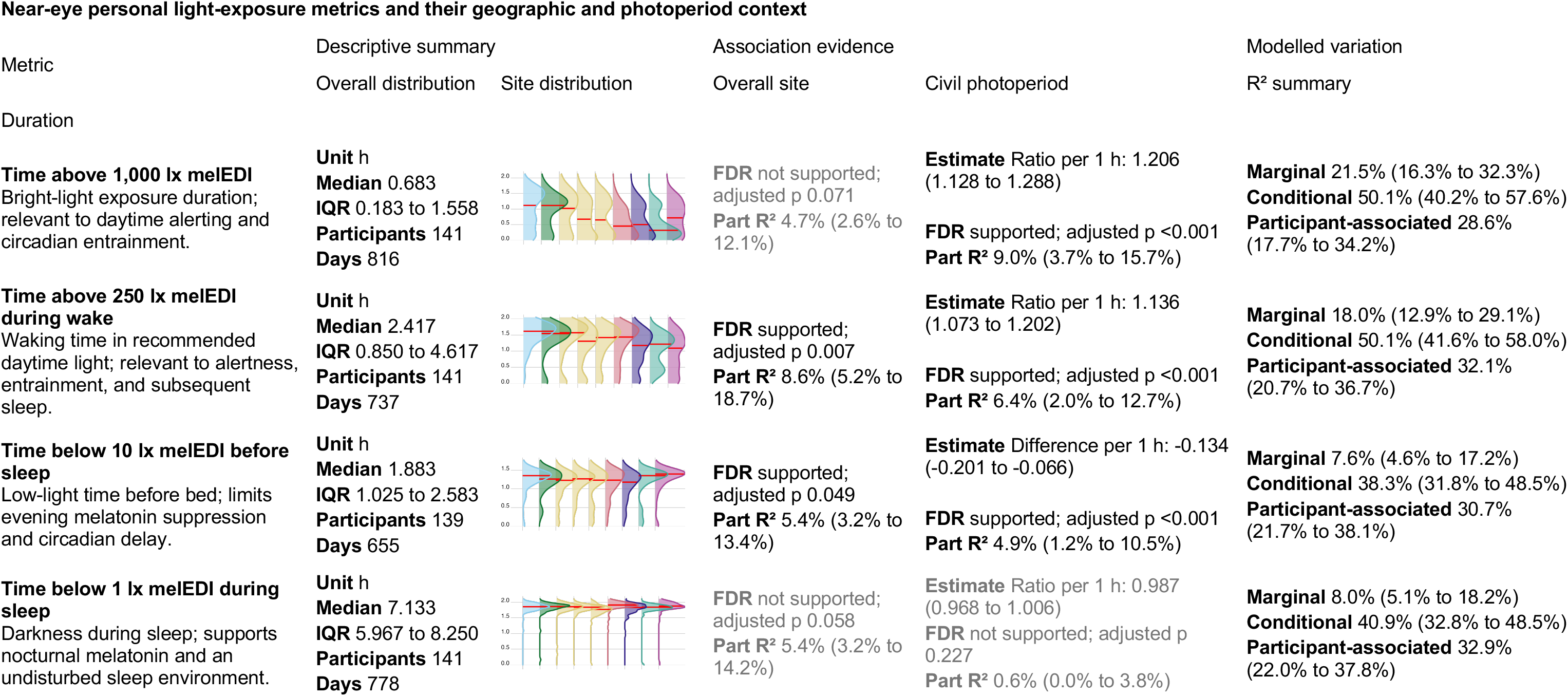
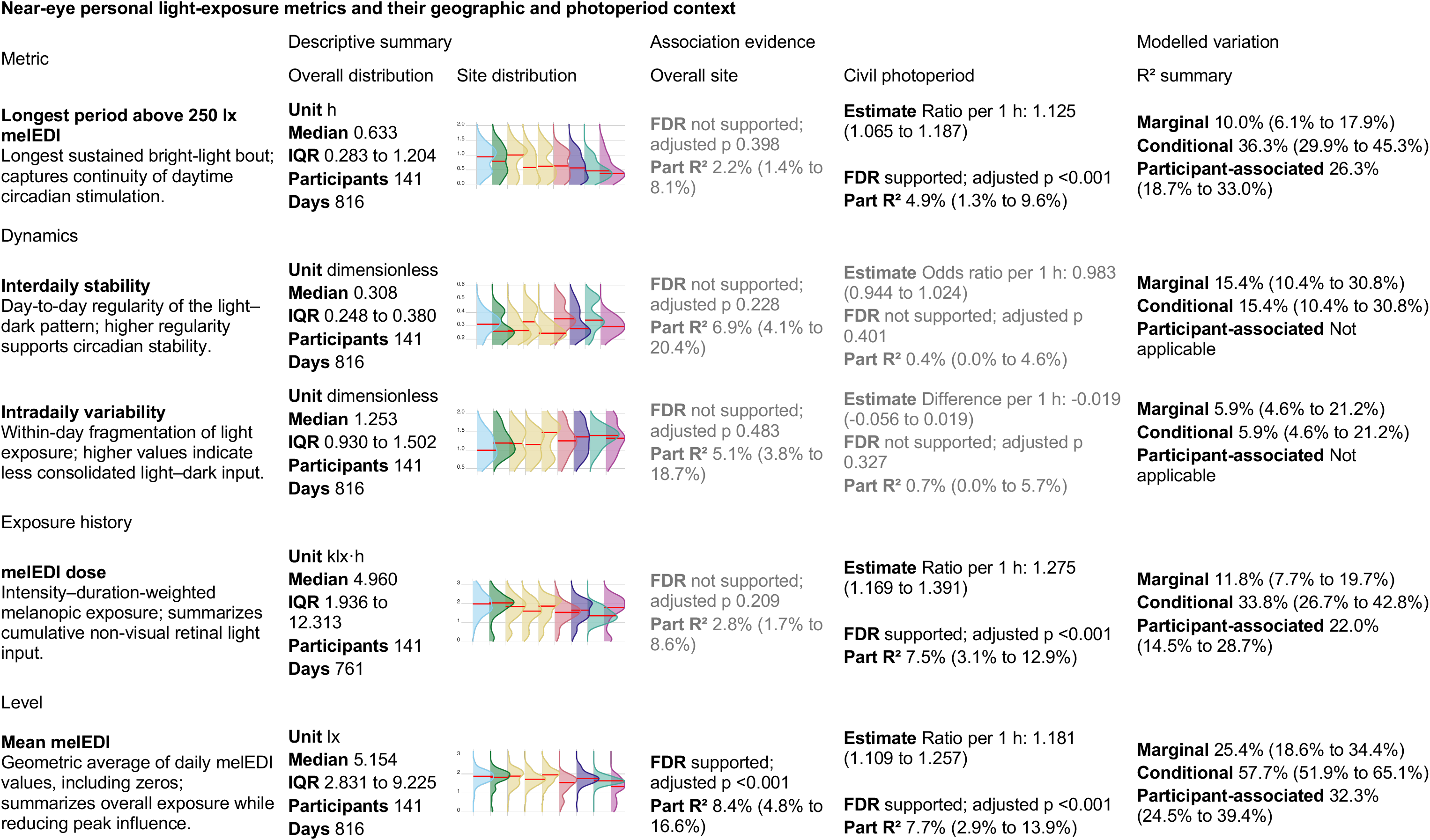
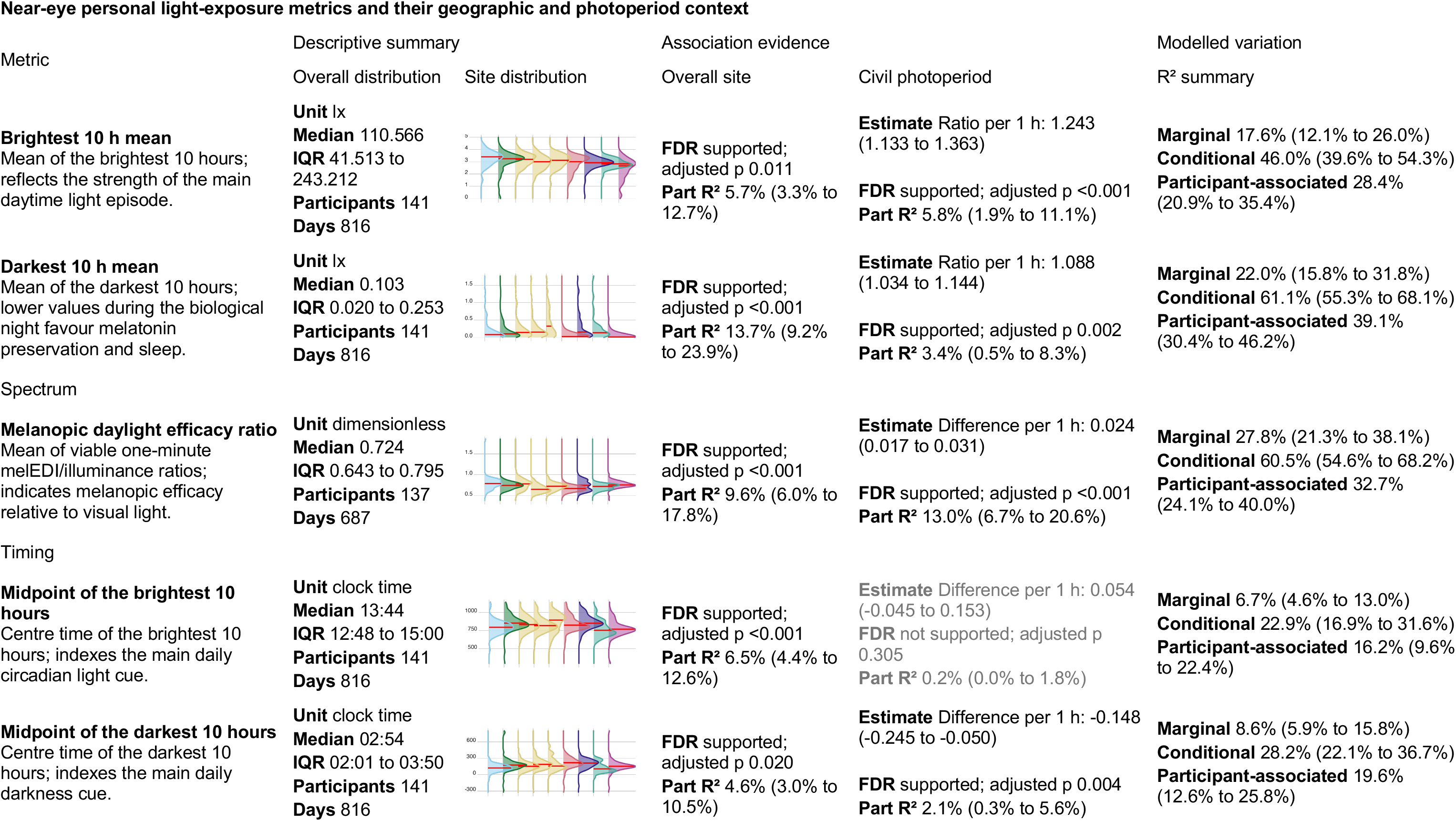
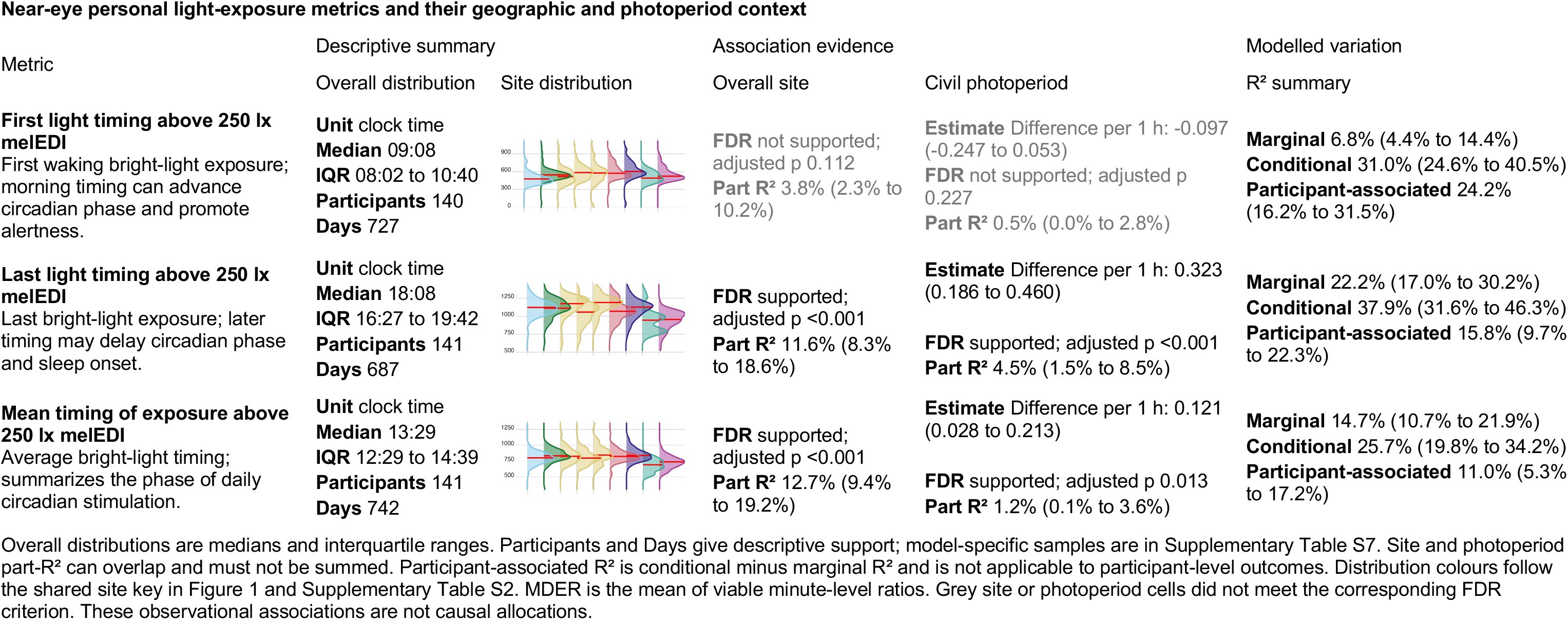
Near-eye personal light-exposure metrics and their geographic and photoperiod context. Descriptive exposure values are medians with interquartile ranges (25th to 75th percentiles). Density plots show site distributions and are descriptive. Metrics are grouped into thematic streams and describe distinct features of the daily exposure pattern rather than interchangeable health-response estimates. Model cells report overall-site evidence, civil-photoperiod associations with 95% confidence intervals, FDR decisions, participant-associated model information and model-specific R² summaries where applicable. Site and photoperiod part-R² values may contain overlapping fitted information and must not be summed. Associations are observational. Absolute latitude is fixed within and confounded with study site, so separate models containing either site or latitude are reported; complete latitude and model-adequacy results are provided in the Supplementary Information.

### Immediate environments and routines reorganise exposure

Self-reported light source was strongly associated with one-hour near-eye exposure, with an FDR-supported light-source-by-site interaction. Site-average interaction-model estimates were 959.6 lx melanopic EDI for outdoor daylight (95% CI 786.8 to 1,170.2), 198.8 lx for indoor daylight (169.3 to 233.4), 85.1 lx for indoor electric light (71.6 to 101.1) and 23.8 lx for a display screen (17.4 to 32.5). Outdoor daylight was 11.27 times the indoor-electric estimate (8.71 to 14.60) and exceeded the 250-lx daytime recommendation, whereas the indoor-daylight and indoor-electric intervals were below it. During reported sleep, darkness and external-light estimates were 2.24 lx (1.54 to 3.26) and 11.18 lx (8.06 to 15.51), both above the 1-lx bedside upper limit. Outdoor electric light remained imprecise relative to indoor electric light near eye (ratio 1.019, 0.336 to 3.090; FDR-adjusted *p* = 0.973). These contextual averages do not classify people or hours as adherent.

External light during sleep in Munich (DE) was estimated at 130.2 lx, 11.65 times the corresponding site average; sleep darkness in Kumasi (GH) was 0.371 lx, 0.166 times its site average. Sparse site-category cells limited some estimates.

The primary analysis used a population-mean quasi-Tweedie model with participant-cluster-robust covariance for uncertainty; lag-one residual correlation remained 0.288. Fitted versus observed exact-zero proportions were 39.8% and 27.8%, respectively. An exploratory mixed-effects model quantified variance explained after adding a participant intercept to site-by-light-source fixed terms. Marginal R² was 0.796 and conditional R² 0.876, an 8.0-percentage-point increase in explained variance from the participant intercept. Of the marginal R², light source accounted for 89.3%, site 7.1% and their interaction 3.7%. Separately, an exploratory time-of-day model attributed 60.4% of variation in fitted hourly patterns to global time of day and 24.2% to light-source deviations. Among these two components, light source accounted for 28.6%. The remaining shares were 5.2% for site, 7.6% for participant patterns and 2.5% for day-to-day shifts.

A sensitivity adjusted the population-average light-source comparisons for each participant’s overall category distribution, distinguishing hour-to-hour differences within participants from differences in category composition between participants. The overall category pattern was retained. For external light during sleep, the near-eye ratio relative to indoor electric light changed from 0.449 in the primary additive model to 0.333 (95% CI 0.190 to 0.585), a 25.8% decrease. Thus, between-participant category composition influenced the magnitude, but not the direction, of this contrast. Supplementary Figure S9 shows the temporal, support and site-specific results.

Immediate setting combined behaviour and environment, with an FDR-supported activity-by-site interaction. Site-average near-eye estimates were 714.2 lx outdoors (95% CI 561.7 to 908.2), 331.2 lx on the road or in a vehicle (274.4 to 399.8), 198.9 lx during office or home work (173.9 to 227.5), 76.4 lx while awake at home (62.0 to 94.1) and 4.28 lx during sleep (3.06 to 5.99). The sleep category included many low or zero observations and occasional high stray-light readings, so its fitted mean need not represent a typical observation. Relative to awake time at home, the ratios were 9.35 outdoors, 4.34 for travel, 2.60 for work and 0.056 during sleep. The outdoor estimate exceeded the 250-lx daytime recommendation, the work estimate remained below it, and the sleep estimate exceeded the 1-lx bedside-environment upper limit. Borås (SE) had a fitted outdoor estimate of 3,054 lx, 4.28 times the site average, whereas Tübingen (DE) had 127 lx, 0.18 times the site average. In an exploratory mixed model of five setting categories, marginal R² was 0.766 near eye and 0.768 at the chest. Adding the participant intercept raised conditional R² to 0.858 and 0.859, an explained-variance increment of 9.2 percentage points at both positions. Of the marginal R², immediate setting accounted for 80.7% near eye and 86.1% at the chest, site 12.5% and 5.9%, and their interaction 6.9% and 8.0%, respectively. Figure 3 shows the immediate-setting patterns and site-specific estimates. Separately, an exploratory near-eye time-of-day model attributed 52.5% of variation in fitted hourly patterns to time of day and 30.7% to immediate setting. Among these two components, immediate setting accounted for 36.9%. The remaining shares were 5.6% for site, 8.4% for participant patterns and 2.8% for day-to-day shifts.

**Figure 3:**
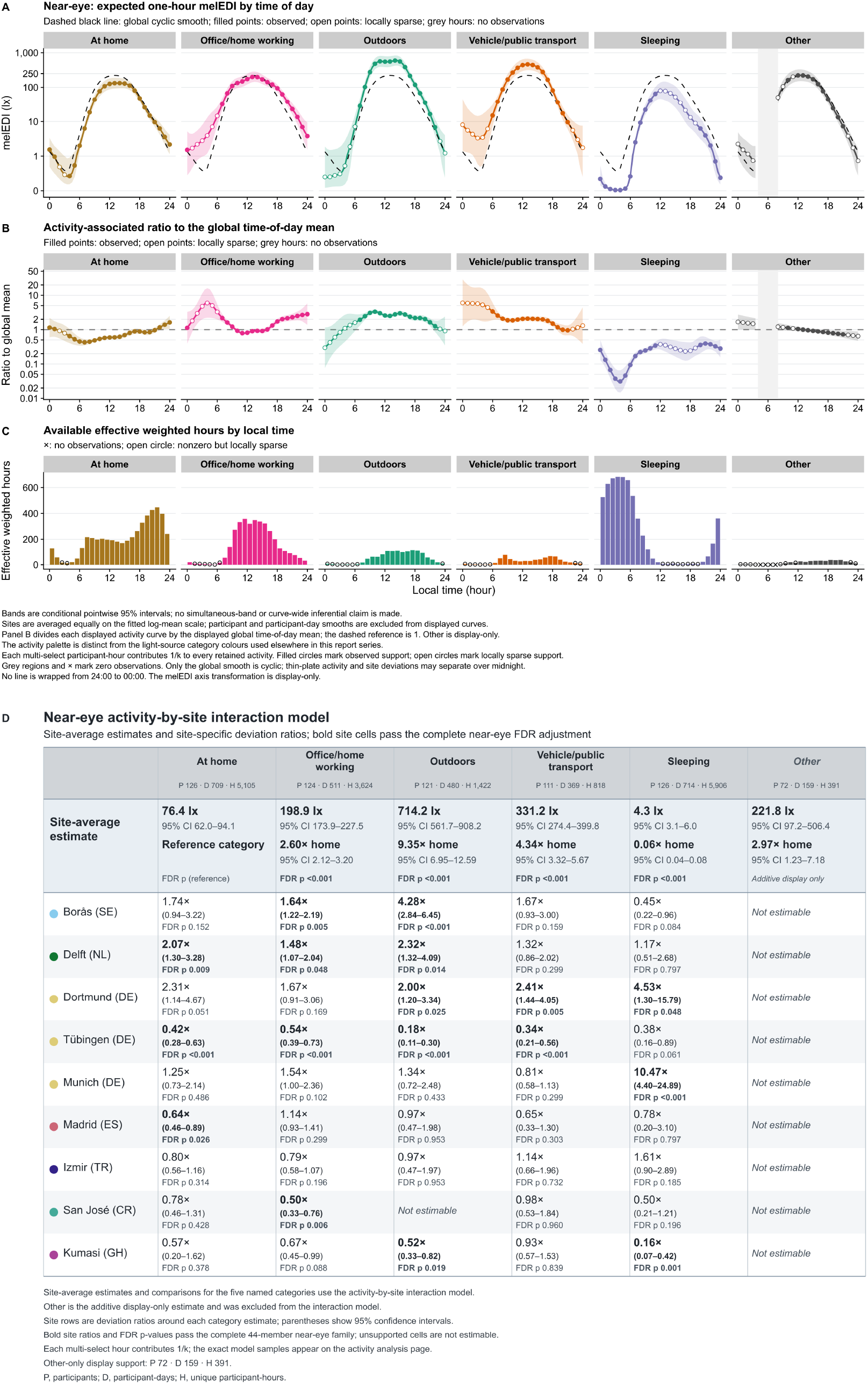
Personal light exposure by immediate setting across time of day and study sites. The diary variable labelled activity combines behaviour and environmental setting. **A–C**, Exploratory time-of-day analysis. Panel **A** shows site-average expected one-hour melanopic EDI with clock-specific 95% confidence intervals; the dashed black curve is the site-average local-clock smooth. Panel **B** shows each setting curve relative to that smooth, with 1 as the reference. Panel **C** shows effective weighted hours; open circles indicate locally sparse support and grey regions indicate times without observations. **D**, Site-average category estimates and site-specific deviation ratios from the activity-by-site interaction model, with participant-cluster-robust 95% confidence intervals and FDR-adjusted p values. Each multi-select participant-hour contributes weight 1/k to every retained setting category. The Other category is sparse and display-only. The fitted frame contains 126 participants, 724 participant-days, 16,135 unique participant-hours, 16,135 effective weighted hours and 4,746 exact-zero participant-hours. Estimates are observational.

Across models of free versus work day, active versus sedentary daily activity status and previous-night sleep duration, the site-average reference was a work, sedentary day following an average 7.8 h of sleep; expected near-eye exposure was 89 lx melanopic EDI (95% CI 67 to 119). Active versus sedentary status was associated with 2.06 times the expected exposure (1.54 to 2.76; FDR-adjusted *p* < 0.001). A model constraining the free-versus-work association across sites estimated 1.45 times higher exposure on free days (1.14 to 1.85; FDR-adjusted *p* = 0.004). Allowing the association to differ by site gave an inconclusive site-average ratio of 1.15 (0.93 to 1.44; FDR-adjusted *p* = 0.298), while some sites differed (interaction *p* = 0.017).

Exploratory ratios were FDR-retained in Borås (SE; 1.95, 1.26 to 3.01) and Dortmund (DE; 2.44, 1.42 to 4.22), with FDR-adjusted *p* = 0.013 for each. The free-versus-work result was also inconclusive under the alternative working variance and sensitive to omitting either site. Each additional hour of previous-night sleep duration, centred at 8 h, had an inconclusive ratio of 0.98 (0.85 to 1.12; FDR-adjusted *p* = 0.745). Excluding six participants outside the employment requirement left 15,871 supported hours from 684 participant-days and 131 participants across all nine sites. The free-versus-work, active-versus-sedentary and previous-sleep associations remained stable within model uncertainty; the predictor-by-site interaction for day type also remained FDR-supported. Supplementary Table S9 and Supplementary Figures S10, S11, S12 and S13 report common, temporal, position-specific and site-specific results.

### Person-level correlates are selective

Self-report analyses did not support general exposure profiling. None of 68 prespecified near-eye associations between four light-exposure-related behaviour (LEBA) factors and 17 metrics retained FDR support. Unsatisfactory model checks made the sleep-environment metric unfit for inference, so its estimates were suppressed. Neither average associations nor site interactions across nine visual-light-sensitivity (VLSQ-8) outcomes retained FDR support at either position. Supplementary Tables S10, S11 and S12, and Supplementary Figures S14 and S15, give the full results.

Each hour later in corrected midsleep on free days (MCTQ questionnaire) was associated with first exposure above 250 lx occurring 0.381 h later (95% CI 0.146 to 0.616; FDR-adjusted *p* = 0.003). Each 10-point shift towards greater morning preference (MEQ questionnaire) was separately associated with first exposure above 250 lx occurring 0.444 h earlier (0.189 to 0.699 h earlier; FDR-adjusted *p* = 0.001). The brightest- and darkest-10-hour midpoints followed the same directions; last exposure and the longest-bright-period midpoint did not retain support. The distinct questionnaires were not combined. Supplementary Table S13 and Supplementary Figure S16 show adjusted estimates; Supplementary Figure S17 shows observed timing distributions.

Each additional decade of age was associated with higher near-eye brightest-10-hour mean, time above 1,000 lx and daily dose, with ratios from 1.27 to 1.31. These cross-sectional associations may reflect cohort, occupation, behaviour or other confounding, not ageing itself. Neither near-eye metric associations with biological sex nor near-eye age-by-site or sex-by-site interactions met the FDR criterion. In a separate daily-pattern analysis, near-eye and chest profiles differed by biological sex (FDR-adjusted *p* = 0.028 and 0.029), but the difference did not meet the FDR criterion after restriction to records with complete immediate-setting data and adjustment. This attenuation is compatible with immediate-setting patterning, but restriction and adjustment cannot be disentangled as mechanisms. Gender was recorded separately but not analysed; its category matched biological sex in all but one case. Supplementary Tables S10, S14 and S15, and Supplementary Figures S18 and S19, give the complete results.

### Measurement position and robustness constrain interpretation

The paired sample of 112 participants and 643 participant-days allowed separate near-eye and chest models to be compared within the same participants and participant-days. The full chest dataset of 154 participants and 902 participant-days broadened coverage beyond the 141 participants and 816 participant-days near eye. At several sites, especially San José (CR), offering a chest sensor enabled recruitment among people reluctant to wear the glasses-mounted sensor. This improved feasibility, not the ocular-exposure sample.

A parallel analysis of overlapping MeLiDos data reported structured position differences across scale, metric, context, site, day and participant^33^. Our separate and common-sample analyses likewise show directional agreement but differences in magnitude or support. Eight-site chest adherence retained daytime and pre-sleep directions and interval conclusions, with calibration and temporal qualifications, but provided no sleep ocular claim or paired test. We neither pooled positions nor inferred equivalence or a universal correction.

Non-estimable planned quantities were four sleep-environment variants (main plus three sensitivity variants) at each position in the light-behaviour analysis (LEBA), and one longest-period sensitivity comparison incompatible with the metric representation. We retained these outcomes rather than replacing them with zero or null findings. Qualifications for zeros, residuals, dependence, nonlinear-term overlap, sparse support and influence remain with the affected analyses and in Methods.

## Discussion

The expected 24-hour rhythm was evident across nine sites in seven countries. The advance was a multiscale account of this everyday exposome component: amplitude across sites, people and days, and its reorganisation across immediate settings. The shared local-clock pattern accounted for the largest share of full-model R² in the daily-pattern analysis. Separate hourly decompositions showed large light-source and immediate-setting contrasts, with the next-largest shares of variation in fitted hourly patterns after time of day. Participant patterns and participant-day shifts together exceeded site patterns in the daily-pattern model, while site, photoperiod and measured person-level associations were metric-specific. Different outcomes and decompositions make this a qualitative synthesis rather than a single quantitative ranking.

The consensus recommendations provide the clearest health-relevant benchmark. Only 24.0% of waking minutes reached the daytime range, while 63.3% of pre-sleep and 87.7% of bedside sleep-environment minutes met their upper limits. Modelled free-day adherence was lower during daytime and preceding sleep, varying by site. Pre-sleep adherence was higher but inconclusive under stricter coverage; temporal dependence remained unresolved. Outdoor exposure exceeded the daytime recommendation, but all other conditions remained below it on average. The fitted sleep-category mean exceeded the bedside limit, while 87.7% of pooled bedside sleep-environment minutes met it. These quantities describe different aspects of the exposure distribution and should be interpreted together. These pooled fractions, recommendation-window proportions and hourly means are distinct estimands. Together, they identify intervention opportunities and shortfalls. The recommendations principally address healthy adults aged 18 to 55 with regular daytime schedules^8^.

This baseline connects mechanistic recommendations with population evidence. Large wrist-sensor cohorts have associated day and night light patterns with psychiatric disorders, incident type 2 diabetes, cardiovascular disease and mortality, and light timing with sleep timing and sleepiness^10–15^. Other cohorts have linked evening, nocturnal or strongly seasonal light with sleep in bipolar disorder, obesity, diabetes, hypertension and metabolic health^16–19^. These outcomes span sleep and major non-communicable diseases. The exposome frames environmental contributions across the life course^20,21^ but rarely includes time-resolved ocular light. This evidence supports personal-light measurement, not clinical risk classification of our exposure categories. Most cohorts used wrist or bedside illuminance without resolving melanopic exposure at the eye, immediate light source or its multiscale architecture.

Exploratory inverse associations of average daytime with sleep and pre-sleep adherence show why the three recommendation windows should not be collapsed into one score. They may reflect schedules, environments or other between-participant differences, not a within-person trade-off or stable phenotype. In exploratory nonlinear models, light source and immediate setting accounted for substantially more variation in fitted hourly patterns than participant-specific patterns: 24.2% versus 7.6%, and 30.7% versus 8.4%, respectively. This finding is consistent with schedules and immediate environments being prominent correlates of hourly exposure, but it is not a causal explanation for the cross-window adherence association. Longitudinal models of dependence and changing routines are needed to test whether improving one part of the pattern displaces or supports another.

Location mattered, but not as one north-to-south ranking. Site differences included daily level, waking bright-light duration, pre-sleep duration, spectral composition and several timing metrics, while dose and regularity did not show the same site pattern. Civil photoperiod was associated with a broader set of metrics, with larger day-length associations for bright-light level and dose than for the darkest daily window. Between 14.2 and 16.2 h of civil photoperiod, six of nine metrics showed signs of uncertainty about a photoperiod effect beyond that point. Sites were sampled in distinct calendar periods and covered unequal photoperiod ranges, leaving latitude, season, weather, temperature, culture, mobility and the built environment entangled. Repeated seasonal sampling across more sites is needed to separate these influences.

Immediate setting patterned exposure, consistent with environmental opportunity, built settings and behaviour^36^. Outdoor daylight and being outdoors exceeded indoor electric light and awake time at home, respectively; active days exceeded sedentary days. Site interactions placed outdoors-category exposure higher in Borås (SE) than Tübingen (DE), while the free-versus-work association was most evident in Borås (SE) and Dortmund (DE). Participant intercepts captured remaining between-participant variation, not individual responses or predictions. Future monitoring should preserve local clock time, location, immediate setting, daylight access and site; self-selection, occupation, mobility and weather remain potential confounders.

Light-behaviour and visual-sensitivity scores did not meet their FDR criteria, whereas chronotype aligned with selected timing outcomes and age with selected bright-light and dose metrics. This accords with evidence linking circadian preference or timing to daylight exposure, sleep and routine, without identifying causal direction^37–40^. Metric-level biological-sex comparisons were inconclusive near the eye. The complete available-data curve differed by biological sex, but immediate-setting-complete analyses did not retain this separation. Person-level patterns may concern specific exposure dimensions rather than a general high-versus-low phenotype.

Near-eye sensing approximates the incident ocular field during wake but is not a retinal measurement; chest sensing samples a related environmental field. Studies show structured differences by body position, morphology, posture and surrounding light^29–31^. The matched sample and parallel placement study show metric- and context-specific MeLiDos differences without a universal correction^33^. The chest option broadened recruitment among people unwilling to wear glasses-mounted sensors, especially in San José (CR). Study design must balance biological proximity, burden and recruitment while keeping estimands position-specific.

The harmonised protocol combined repeated near-eye sensing with local-clock and contextual information across nine sites and seven countries. A generalised additive model separated the shared 24-hour pattern from site, participant and participant-day layers obscured by daily summaries. Collecting these data demanded substantial effort from participants and study teams. Glasses-mounted loggers required fitting, explanation and daily handling; removal, occlusion, charging, discomfort, weather and social acceptability all affected measurement quality^41–43^. Despite this burden, the protocol measured closer to the incident ocular field at high temporal and contextual resolution.

Several limitations qualify these findings. Modest site samples limit site estimates, interactions and subgroup evaluation. Healthy working-age adults may not represent children, adolescents, older or clinical populations, shift workers, people outside employment, or those with restricted mobility or atypical living conditions. Sparse latitude-by-photoperiod coverage leaves latitude, seasonality, culture, climate and built environment entangled and requires broader seasonal sampling. Retrospective hourly setting and light-source reports enabled harmonisation but could not distinguish fine-grained settings or behaviours within home, work or transport. Very low readings require caution. The manufacturer specifies an operating range starting at 1 lx, and independent laboratory testing began at 2 lx^44^. A scene-based field validation found high overall agreement with laboratory-grade spectral references, but underestimation increased at lower intensities and varied with lighting, time, scene complexity and site^45^. Single-digit values therefore approximate low-light exposure rather than provide precise measurements. Finally, no health outcome was measured.

This nine-site reference locates everyday ocular light exposure within the exposome, benchmarks it against physiology-based recommendations, highlights within-site variation among people and days, and identifies temporal, contextual and positional information for future cohorts. Everyday ocular light exposure is structured, measurable and strongly shaped by behaviour and micro-environment, but cannot be reduced to site, latitude or photoperiod. Next steps are broader sampling across photoperiods, latitudes and locations and prospective links to sleep, mental, metabolic, cardiovascular and other outcomes. Causal health benefits or policy effects require interventions and representative longitudinal designs. Potential interventions through clinical advice, occupational health, building design, urban planning or public-health guidance must address the contexts in which people receive light. Time-resolved near-eye measurement offers a practical route from circadian biology to precision prevention and context-sensitive environmental health.

## Methods

### Study design and sites

MeLiDos is a prospective multicentre observational field study of personal light exposure conducted with a harmonised protocol^35^. Participants completed an eight-day ambulatory protocol comprising continuous light logging, repeated smartphone assessments and daily diaries under free-living conditions. At an in-person registration visit, participants provided consent, completed baseline questionnaires, received study instructions and were fitted with the loggers. Seven consecutive recording days separated registration and return visits; most sites recorded from Monday to Monday, while Dortmund (DE) recorded from Tuesday to Tuesday. If a return visit was delayed, participants continued until formal study termination and device retrieval. Data were collected between August 2023 and October 2025 at Borås (SE), Delft (NL), Dortmund (DE), Tübingen (DE), Munich (DE), Madrid (ES), Izmir (TR), San José (CR) and Kumasi (GH). The Kumasi (GH) and Izmir (TR) collections are described in site-specific data notes^46,47^. Sites were an opportunity sample of consortium partners and were not selected to represent national populations. Questionnaire materials were administered in the local language except in Tübingen (DE) and Kumasi (GH), where English was used, and translated materials followed a harmonised translation, review, adjudication, pretesting and documentation process^48^.

### Participants, recruitment and ethics

The registered target population comprised adults aged 18 to 65 years in full-time employment or part-time employment above 80%. Exclusion criteria included diagnosed psychiatric or sleep disorders, shift work in the preceding two months, regular tobacco or recreational drug use, photosensitising medication, visual impairment incompatible with the devices and residence outside the local study area during recording. The study population included nine participants outside the registered age or employment criteria: one participant older than 65 years, two recorded as not employed and seven recorded as marginally employed; one participant met both criteria for exclusion. For the age and biological-sex analysis, a joint age-and-employment exclusion sensitivity removed six participants from the near-eye models and nine from the chest models. All 11 FDR-supported main findings retained their direction and support, and none of the 68 main-effect FDR decisions changed. For the day-type, exercise and sleep analysis, a separate near-eye employment sensitivity reduced the sample from 137 to 131 participants; the directions and FDR conclusions for the main associations, and the site-heterogeneity conclusions, were unchanged. Students and trainees were retained in both checks.

The target was at least 15 participants per site, based on prior power calculations and allowing for up to one-third dropout or incomplete recording (15->10)^49^. The study roster was below 15 participants only in Munich (DE; n = 10), and no site had fewer than 10 participants. The final near-eye sample was below 15 in Borås (SE; n = 13), Delft (NL; n = 13), Munich (DE; n = 10) and San José (CR; n = 6); San José was the only site below the dropout-adjusted threshold of 10 (Table 1). Participants were recruited through local advertisements and institutional mailing lists. At some sites, especially San José (CR), reluctance to wear the glasses-mounted near-eye sensor limited recruitment, and the chest option broadened recruitment possibilities.

The small near-eye sample in San José (CR; n = 6) limits the precision of site-specific estimates and the ability to detect associations within that site. The reported leave-one-site-out sensitivity analyses thus assessed the influence of individual sites on pooled and site-average estimates. The larger chest-level sample in San José (n = 39) provides more extensive complementary data, although findings from this measurement position remain distinct from near-eye exposure in all reported analyses.

All participants provided written informed consent. Approvals were obtained from the Ethics Committee of the Technical University of Munich for Tübingen (DE; 2023-115-S-KK) and for Munich (DE) and Dortmund (DE; 2024-118-S-SB); the Dortmund (DE) site at the Federal Institute for Occupational Safety and Health (BAuA) operated under this TUM multicentre approval. The other approvals were from the Swedish Ethical Review Authority for Borås (SE; 2024-06792-01), the THUAS Ethics Committee for Delft (NL; positive review dated 29 May 2024), the Research Ethics Committee of Universidad San Pablo-CEU for Madrid (ES; 834/24/106), the Scientific Research and Publication Ethics Committee of the Faculty of Science and Engineering at İzmir Institute of Technology for Izmir (TR; decision 5/1), the Comité Ético Científico of Universidad de Costa Rica for San José (CR; CEC-279-2025), and the Committee on Human Research, Publications and Ethics of Kwame Nkrumah University of Science and Technology for Kumasi (GH; CHRPE/AP/644/24). The protocol was first trialled with participants at the Tübingen (DE) site, whose feedback informed procedural refinements before implementation at the remaining sites; further details are reported in the published protocol^35^.

### Personal light measurements

Personal light was recorded every 10 s with spectrally sensitive ActLumus loggers (Condor Instruments, São Paulo, Brazil). Eight visible-band channels were processed by the device to generate photopic and alpha-opic quantities, including melanopic EDI^6^. The manufacturer specified a factory-calibrated operating range of 1 to 100,000 lx and 10% precision at 1,000 lx. Independent testing against a criterion spectrometer across indoor LED and outdoor daylight conditions covered 2 to 100,000 lx and reported mean photopic error of +3.3% (SD 13.4%) for ActLumus^44^.

A subsequent scene-based field validation found high overall agreement with laboratory-grade spectral reference instruments, together with systematic underestimation that varied with intensity and setting^45^. Participants wore the near-eye device centrally on non-prescription spectacle frames and, where available, a complementary pendant-mounted device at the chest. The near-eye position measures light closer to the eye but is not a direct retinal measurement. The chest position provided a separate measurement of the local environmental light field.

Paired common-sample analyses evaluated how closely it tracked the near-eye measurement as a potential ocular proxy. During reported sleep, devices were placed facing upward on a bedside surface near head level, so both positions characterise the bedside sleep environment. Participants recorded removals and placed devices in an opaque bag during waking non-wear^43,50^.

The placement analysis from the same project used overlapping MeLiDos data and is cited as related methodological evidence, not independent replication^33^. No analysis pooled sensor positions or estimated a universal conversion. Common-sample analyses retained the same participants and participant-days or participant-hours at both positions and fitted each position separately.

### Contextual and person-level measures

Participants completed the core Consensus Sleep Diary after awakening^51^, together with evening reports and paper diaries recording light source and an activity-environment category for each hour. Smartphone assessments of sleepiness and mood were scheduled at 11:00, 14:00, 17:00 and 20:00, and evening questionnaires assessed wellbeing and physical activity^52–54^. Smartphone measures and baseline or end-of-study questionnaires used MyCap integrated with REDCap^55,56^. Participants photographed and uploaded the paper hourly diaries each day.

The modified Harvard Light Exposure Assessment categories were indoor electric light, outdoor electric light, indoor daylight, outdoor daylight including shade, an emissive display, darkness during sleep and external light during sleep^57^. The primary light-source analysis used one retained category per participant-hour. The diary variable labelled activity combined behaviour and environmental setting. Its categories distinguished sleep, awake time at home, motorised travel, active travel, office or home work, outdoor work, outdoor leisure and other activities. We therefore refer to this variable as immediate setting in the narrative. For analysis, outdoor travel, outdoor work and outdoor leisure were combined as Outdoors. Reports could contain concurrent categories; duplicates were removed and an hour with *k* retained labels contributed weight 1/*k* to each, preserving total weight one. Day type was coded as work or free day, daily activity status as sedentary or active, and previous sleep duration was linked from the preceding sleep period.

Baseline measures recorded age, biological sex, gender, employment and chronotype; end-of-study measures covered visual light sensitivity, the sleep environment and habitual light-exposure behaviour. Chronotype was represented separately by corrected midsleep on free days from the Munich Chronotype Questionnaire and the Morningness-Eveningness Questionnaire, whose higher scores indicate greater morning preference^58,59^. Visual sensitivity used the VLSQ-8^60^, sleep environment used the Assessment of Sleep Environment^61^, and four Light Exposure Behaviour Assessment (LEBA) factors represented outdoor time, device use in bed, ambient light before bed and morning or daytime light use^62^. Biological sex and gender were recorded as separate variables. Models analysed biological sex, coded Female or Male, but did not analyse gender; its recorded category matched biological sex in all but one case.

### Time, sleep/wake status and coverage preparation

Processing preserved both actual time in Coordinated Universal Time and local wall-clock time. Actual time ordered observations and measured elapsed gaps; local time aligned daily patterns across sites. Repeated local minutes during autumn clock changes remained distinct in actual time, and structurally absent spring minutes remained missing. Melanopic EDI values at or above the validated 100,000-lx operating boundary were unavailable analytically but retained in the raw channel with a flag.

Diary-defined sleep began at reported sleep preparation and ended at reported wake. Sleep values were retained as the intended bedside record. Declared waking non-wear made the analytical light value unavailable; missing wear labels were not treated as non-wear. Complete 1,440-minute local participant-day grids preserved unsupported periods as missing rather than deleting rows or replacing them with zero. A day was eligible when at least 80% of this combined waking and bedside-sleep record had valid melanopic EDI. Hourly summaries required at least 50% valid minutes in that hour, but an unsupported hour did not erase valid minutes from the daily decision or minute-based metrics. Two otherwise eligible near-eye days and three chest days with finite but entirely zero melanopic EDI were excluded by the defined exact-all-zero screen.

### Exposure metrics

Metrics were derived in R^63^ with LightLogR^64,65^. The 17 outcomes represented complementary properties of the daily pattern. Three level metrics described the daily geometric mean melanopic EDI and the mean melanopic EDI in the brightest and darkest 10-hour windows. Five duration metrics quantified time above 1,000 lx, waking time above 250 lx, pre-sleep time below 10 lx, sleep-environment time below 1 lx and the longest continuous period above 250 lx. Five timing metrics located the brightest and darkest windows and the first, last and mean timing above 250 lx. Melanopic EDI dose integrated exposure over time, melanopic daylight efficacy ratio characterised spectral composition relative to visual illuminance, and interdaily stability and intradaily variability characterised regularity and fragmentation. Geometric means were calculated from log10(melanopic EDI + 0.1 lx), followed by inverse transformation and removal of the same offset^66^. All-observed-zero windows were retained as exact zero within verified numerical tolerance. Metric-specific support rules made only the affected metric unavailable, not the participant-day. A longest-period duration remained a lower bound when missing data could conceal a longer period; exactly identified periods were examined separately.

The melanopic daylight efficacy ratio was the arithmetic mean of viable one-minute melanopic EDI to photopic-illuminance ratios on a complete 1,440-position local-clock grid. Both channels had to be finite and strictly positive, and at least 720 positions were required. Repeated autumn minutes were averaged within each channel before division, and absent spring minutes remained missing.

### Recommendation-window and adherence analyses

We compared valid one-minute values with the Brown et al. recommendation ranges^8^: at least 250 lx melanopic EDI during daytime, no more than 10 lx during pre-sleep and no more than 1 lx during sleep. Daytime comprised diary-defined wake excluding the three hours before reported sleep, pre-sleep comprised those three hours and sleep comprised the reported sleep interval. Each descriptive numerator was divided by all valid near-eye-record minutes in the corresponding window and pooled across the analysed sample. Daytime and pre-sleep used the worn near-eye record; sleep used the bedside sleep-environment record.

The main adherence analysis retained one row per window period and defined adherence as adherent valid minutes divided by all valid minutes in that period. Each cycle linked the sleep interval ending at wake, the ensuing daytime interval excluding pre-sleep, and the following three hours before the next reported sleep. All three inherited the work or free label of the wake-start calendar date, even when a window crossed midnight. Actual diary dates and UTC boundaries established chronology; gaps between diary dates were not bridged. Windows qualified independently, so a cycle need not contain three eligible windows. The any-valid primary sample retained at least one valid comparison; the coverage sensitivity required valid minutes for at least 80% of each expected window interval. Expected duration defined coverage, not the likelihood denominator. Missing minutes were not imputed. Exact 0% and 100% periods remained in an endpoint-inflated beta-binomial model. Its mean included the full window-by-site-by-day-type interaction and a participant intercept; extra wholly non-adherent and wholly adherent periods and window-specific dispersion were modelled separately. Population-average adherence integrated the participant distribution and gave each of nine sites equal weight. Separate FDR families covered three window-specific free-minus-work contrasts, three window-specific site interactions, the global interaction, 27 site-specific contrasts against zero, 54 site-minus-average levels and 27 site-effect-minus-average contrasts. The last family’s coverage repetition assessed stability without another FDR adjustment. Variance decomposition used balanced 54-cell global and 18-cell within-window references; Shapley allocations and R² were point-only. Chest models used a separate two-window, eight-site sample and did not test paired placement differences.

The exploratory cross-window extension used daytime adherence as the predictor and the preceding sleep or following pre-sleep adherence as outcomes under the same wake-start-date alignment. It retained pairwise-complete observations, not necessarily complete three-window cycles: 1,376 outcome rows comprised 758 daytime-sleep and 618 daytime-pre-sleep pairs from 140 participants and 761 cycles. Its predefined eligibility rules did not include every potential main-model pair. The model separated each participant’s cycle-specific daytime deviation from their observed average, adjusting for window, site, day type, their corresponding interactions and the participant’s day-type composition. Response-scale contrasts used equal-site and 50:50 work/free weighting. The four within- and between-participant associations formed a separate FDR family. Positive serial dependence remained unresolved and some sensitivity fits were non-estimable, so no within-participant day-level claim was stated. Descriptive profiles used unweighted equal-cycle participant means, with daytime deduplicated within participant and cycle, for 139 complete profiles and 417 participant-window points. These are anonymous monitoring-period summaries, not ranks or stable participant types.

The main model passed all five applicable endpoint-calibration checks in both samples, but temporal dependence still qualified inference. The 80% coverage restriction retained the daytime and sleep directions and interval conclusions; pre-sleep retained direction but not exclusion of zero. Diary-indexed grouping was a separate sensitivity. Pre-sleep interval conclusions also changed under 70% and 90% coverage and restriction to participants observed on both day types. Actual-date temporal endpoint models failed structural checks. Their beta-binomial fallbacks changed both the response family and random-effects structure, retained boundary cautions and failed all five endpoint-calibration checks in both samples. They did not replace the primary model or resolve its temporal limitation. All nine site-deletion checks and four participant-deletion checks were estimable; one participant deletion and the ordinary-binomial diagnostic failed and supplied no derived inference. The separate chest model retained calibration and temporal qualifications. In the exploratory extension, the 80% restriction preserved all four directions and interval conclusions, but incomplete influence checks and unresolved dependence limited robustness claims.

### Daily temporal architecture

The daily-pattern analysis used supported 30-minute arithmetic means, retaining a bin when at least 15 one-minute values were valid. The outcome was log10(melanopic EDI + 0.1 lx). A generalised additive model fitted with `mgcv::bam()` and fREML combined a cyclic common local-clock smooth, sum-to-zero site curves, participant curves and participant-day random intercepts^67–70^. A boundary-aware first-order autoregressive working structure prevented residual sequences from crossing participant-day, clock-repeat or gap boundaries. Fitted-curve dispersion was compared on a common 48-bin grid. A Shapley decomposition averaged each component’s addition to R² across all component orders, thereby distributing the model’s explained variance^71,72^. The decomposition partitioned the full model’s row-weighted in-sample R² for log10(melanopic EDI + 0.1 lx), defined as the reduction in squared prediction error relative to predicting the fitted-sample mean. The percentages therefore partition full-model R² on that transformed response scale. They are not percentages of variance explained in raw melanopic EDI and do not indicate causal or out-of-sample predictive importance. Confidence intervals for dispersion and R²-share ratios used 2,000 hierarchical cluster resamples while holding the fitted model structure fixed.

### Geographic and photoperiod analyses

We analysed 17 participant-level or participant-day metrics with metric-specific Gaussian or Tweedie exponential-dispersion models^73,74^. We treated overall site differences, civil photoperiod (day length including civil twilight), absolute latitude and the adequacy of a linear latitude term relative to site as four separate model questions, each adjusted as a complete 17-test FDR family. Participant-day models included participant random intercepts. Mixed-effects models were fitted with `lme4` or `glmmTMB`, and component R² values were obtained with `performance` where reported^75–77^. Site and latitude were not entered as independent fixed predictors in one model because each site has one latitude; site and latitude models instead used the same observations and photoperiod adjustment. Site follow-up contrasts were calculated with `emmeans` only after a supported overall site test and compared each site with a site-average estimate, obtained by giving each included site equal weight^78^.

Nonlinear photoperiod analyses fitted each of nine planned metrics separately with a generalised additive model estimated by restricted maximum likelihood, a thin-plate photoperiod smooth with basis dimension six and participant random effects nested within site. The derivative, meaning the slope of the fitted curve, was evaluated with `gratia` at 100 equally spaced photoperiod values with 95% intervals^79^. A qualifying descriptive transition required the preceding slope interval to be wholly above zero, the next interval to include zero and every later interval through the recorded maximum to include zero as well. This procedure located a change in the fitted slope but did not test equivalence, a mechanism or a physiological ceiling. Expanded-basis, fixed-site, common-sample and leave-one-site-out analyses assessed stability. The gap-timing-unaware sensitivity dataset still passed the 50%-per-hour and 80%-per-day rules, but did not use the timing of remaining missing observations for an additional metric-specific adjustment.

### Immediate-context and routine analyses

Light-source and activity-environment analyses modelled the zero-aware one-hour geometric mean melanopic EDI with fixed-power quasi-Tweedie log-mean models and participant-cluster-robust inference^73,74^. Additive fixed-site models supplied primary category tests. Category-by-site interaction models supplied site-average category estimates, ratios and site-specific context when the interaction was supported. The activity-environment analysis retained fractional 1/*k* weights for concurrent labels. FDR adjustments were applied to the declared category and site-contrast families. Exact-zero calibration, residual dependence, sparse cells and deletion refits were retained as model checks. A secondary population-mean sensitivity added six participant-level category-proportion terms to the light-source model while retaining participant-clustered covariance. This separated hour-to-hour category differences within participants from between-participant differences in category composition; it did not estimate participant random slopes. Exploratory `glmmTMB` Tweedie log-link models fitted by maximum likelihood then added one common participant intercept to site-by-category fixed effects at the near-eye position for light source and separately at both positions for the activity-environment variable, with the power fixed at 1.539919^76,77^. The activity-environment models retained the five named categories, excluded Other-only hours and used exact 1/*k* weights. Their fixed-predictor variance used the same fractional weights so every participant-hour contributed one unit. Marginal and conditional R² used a lognormal distribution-specific variance approximation, and hierarchy-respecting Shapley allocations distributed marginal R² across site, category and their interaction. Percentages from the exploratory time-of-day models instead partition variance among fitted linear-predictor values. Because they do not compare this with observed outcome variance, they are not R². Results describe them as variation in fitted hourly patterns, not variance explained in raw exposure. These were model-based point descriptions without bootstrap intervals. No model included random category slopes.

The participant-hour analysis modelled each supported participant-hour with a zero-inclusive quasi-Poisson log-link model, fixed site associations and participant-cluster HC3 uncertainty. Free versus work day, active versus sedentary daily activity status and previous-night sleep duration centred at 8 h were entered together. Three-test FDR families were applied separately to pooled associations, predictor-by-site interactions and practical contrasts.

### Person-level analyses

Light-related behaviour, visual sensitivity, chronotype, age and measured biological sex were evaluated with site-adjusted metric-specific models. Participant-day outcomes generally used Gaussian or Tweedie mixed models with participant random intercepts; participant-level dynamics outcomes used fixed-effect models. Each analysis retained its complete prespecified false-discovery-rate families, including non-estimable outcomes. Chronotype instruments were modelled separately across five timing outcomes. Age associations were scaled per decade, and female-minus-male contrasts used male as the reference. Average associations and predictor-by-site interactions were distinct questions.

The complete biological-sex curve analysis used supported 30-minute arithmetic means transformed as log10(melanopic EDI + 0.1 lx). The generalised additive model included cyclic common and female-specific curves, site curves, participant curves, participant-day random effects and boundary-aware first-order autoregression. The primary test jointly assessed the time-constant and time-varying female-minus-male contributions across the complete 24-hour curve. Clock-specific uncertainty was participant-cluster robust. Level-versus-shape attribution and adjustment for the activity-environment variable were secondary or exploratory analyses.

### Multiplicity, uncertainty, model checks and sensitivities

Tests were two-sided. We controlled the false discovery rate with the Benjamini-Hochberg procedure separately within each declared complete family^80^. We report 95% confidence intervals and FDR-adjusted *p* values for confirmatory families. A result that did not meet an FDR-adjusted criterion was treated as inconclusive rather than as evidence of absence or equivalence. Unless stated otherwise, site-average estimates gave each included site equal weight on the fitted scale before back-transformation. Exploratory analyses retained separate multiplicity families and are labelled as such.

Model checks addressed convergence, rank, singularity, residual distribution, zero mass, fitted support, heteroscedasticity, serial dependence, nonlinear-term overlap and participant or site influence. Qualifications were attached to the affected results.Prespecified or structured sensitivities included the separately prepared gap-timing-unaware dataset, common sensor-position samples, alternative support or metric definitions, model-form changes, participant deletion and leave-one-site-out checks. Near-eye and chest models remained separate in every comparison. Analyses were conducted in R, with `tidyverse` packages supporting data handling and reporting^63,81^. Exact package versions, formulas, inputs and output files are documented in the analysis pages and project repository.

### Preregistration and deviations

The study was preregistered with AsPredicted (#273407)^82^. The final analyses preserve the registered scientific questions but include documented changes to placement priority, state and coverage rules, metric definitions, outcome construction, model families, site structure, multiplicity and sensitivity handling. The main manuscript is organised around the resulting scientific estimands. Supplementary Methods: Preregistration deviations (supplied as a separate Word document) describes the scientific changes and qualifications.

## Supporting information

Supplementary Methods: Preregistration deviations

## Data availability

All data used in this study are available through the MeLiDos Project GitHub organisationand are archived in the MeLiDos Zenodo community. Public site datasets have persistent records for Kumasi (GH), Madrid (ES), Dortmund (DE), Izmir (TR), Tübingen (DE), San José (CR), Munich (DE), Borås (SE) and Delft (NL)^83–91^. These records and other project resources are indexed through the MeLiDos Project Data Hub^92^ and accessed reproducibly through the documented `melidosData` interface^93^.

## Code availability

The project repository contains the analysis code, local input data, and instructions for reproducing the manuscript, supplementary information, and analysis webpages with Quarto.

## Acknowledgements

We thank everyone who volunteered for the study, including those who did not complete the protocol, and the MeLiDos partners Iberoptics and LNE for lending wearable devices used in this research.

## Author contributions

- **Conceptualization:** J.Z., A.D., S.A., J.B., O.S. and M.S.
- **Data curation:** J.Z., A.D., G.K.A., S.A., S.G.A., S.N.A., Z.K., D.B.M., M.N.T., S.M.T., N.H., S.H., A.J., D.B., A.S.S. and M.S.
- **Formal analysis:** J.Z.
- **Funding acquisition:** A.D., K.O.A., K.B., D.B.M., M.N.T., J.B., S.K., A.S.S. and M.S.
- **Investigation:** A.D., G.K.A., S.A., S.G.A., S.N.A., Z.K., M.C.P.G., R.A.G.L., S.L., D.B.M., M.N.T., S.M.T., N.H., S.H., A.J., C.G., D.B., I.S., A.S.S., H.v.B., K.B., S.K. and M.S.
- **Methodology:** J.Z., A.D., K.O.A., K.B., C.G., D.B.M., M.C.P.G., R.A.G.L., M.N.T., S.M.T., O.S., G.C.G. and M.S.
- **Project administration:** J.Z., A.D., G.K.A., S.A., S.G.A., S.N.A., K.O.A., D.B.M., J.B., A.S.S., K.B., A.J. and M.S.
- **Resources:** J.Z., A.D., G.K.A., K.O.A., J.B., K.B., A.S.S. and M.S.
- **Software:** J.Z.
- **Supervision:** J.Z., A.D., S.A., J.B. and M.S.
- **Validation:** J.Z., A.D. and M.S.
- **Visualization:** J.Z.
- **Writing – original draft:** J.Z.
- **Writing – review & editing:** all authors.

## Supplementary Information

### Supplementary materials

**Supplementary Methods: Preregistration deviations** is supplied as a separate Word document (`preregistration-deviations.docx`). It describes all departures from the registered plan and the associated qualifications.

Near-eye and complementary chest measurements remain separate sensor-position estimands. Common-sample analyses use the same participants and participant-days at both positions and fit the positions separately. They do not pool positions, test equivalence, or establish a universal correction. Detailed model checks, sensitivities, exact formulas, and package versions are documented in the analysis pages. The project repository provides the data, code and reproduction instructions.

### Supplementary figures

**Supplementary Figure S1.**
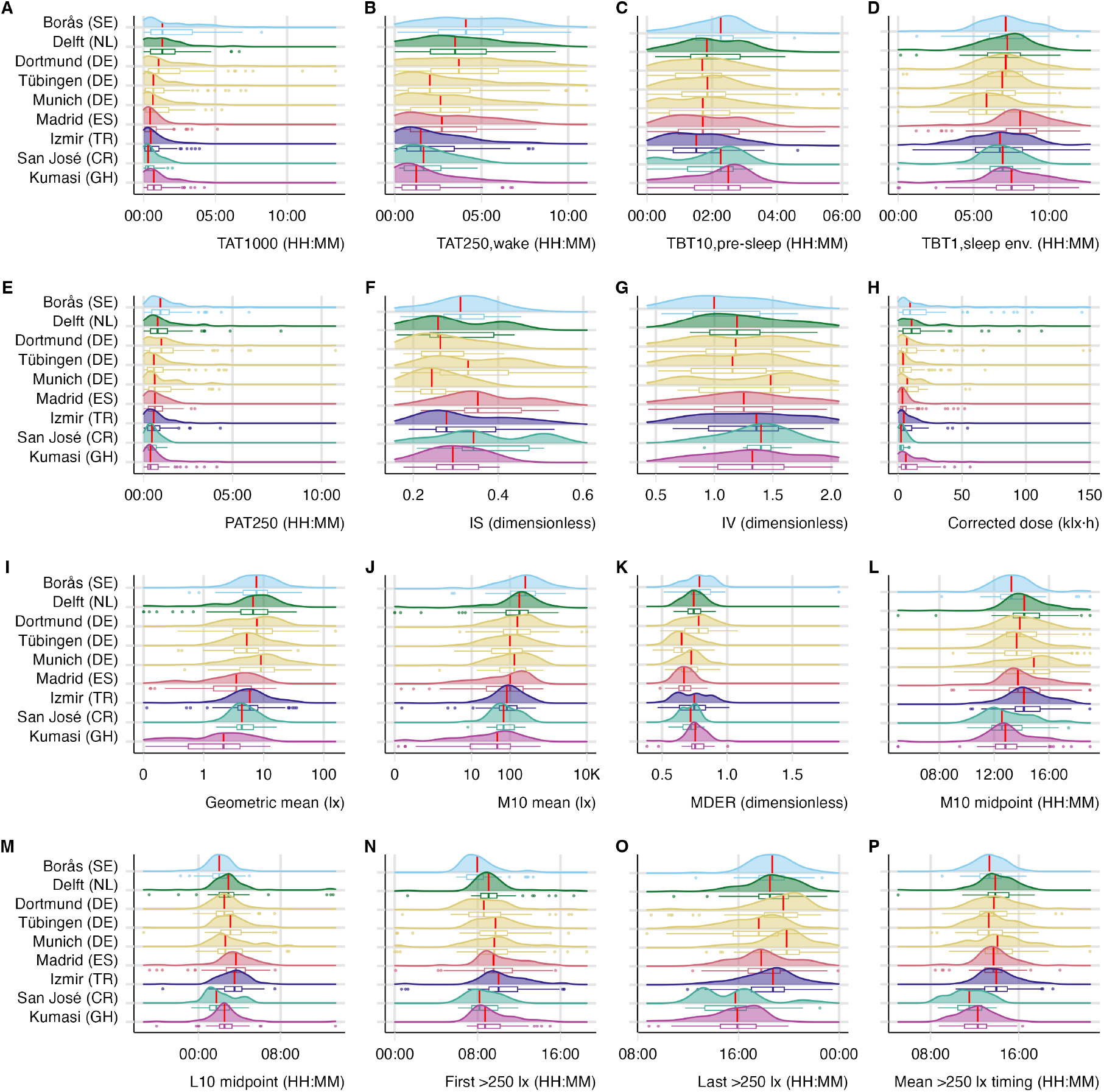
Detailed near-eye metric distributions by country-coded study site.

**Supplementary Figure S2.**
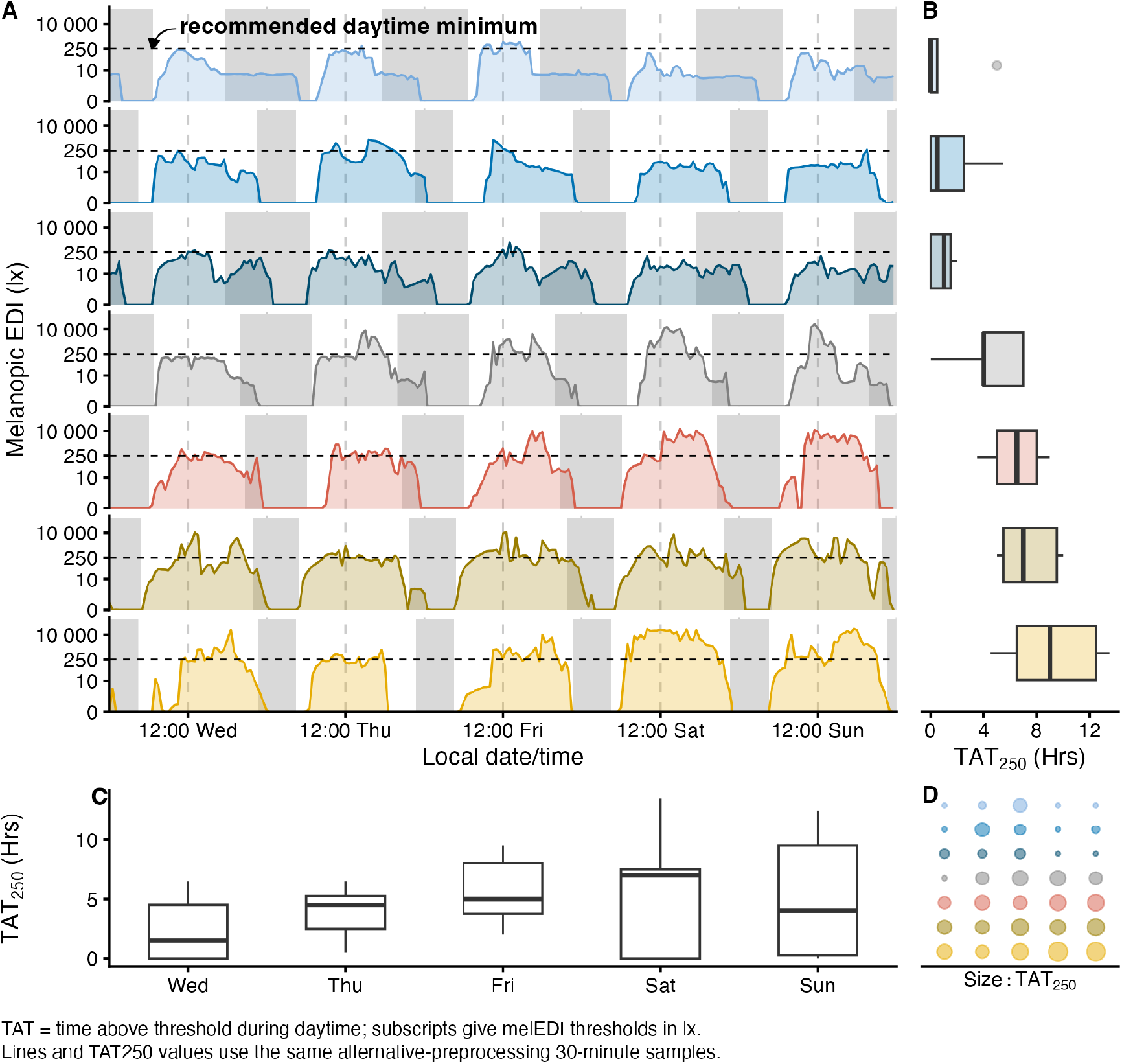
Examples linking 30-minute time series to the daily time-above-250-lx metric.

**Supplementary Figure S3.**
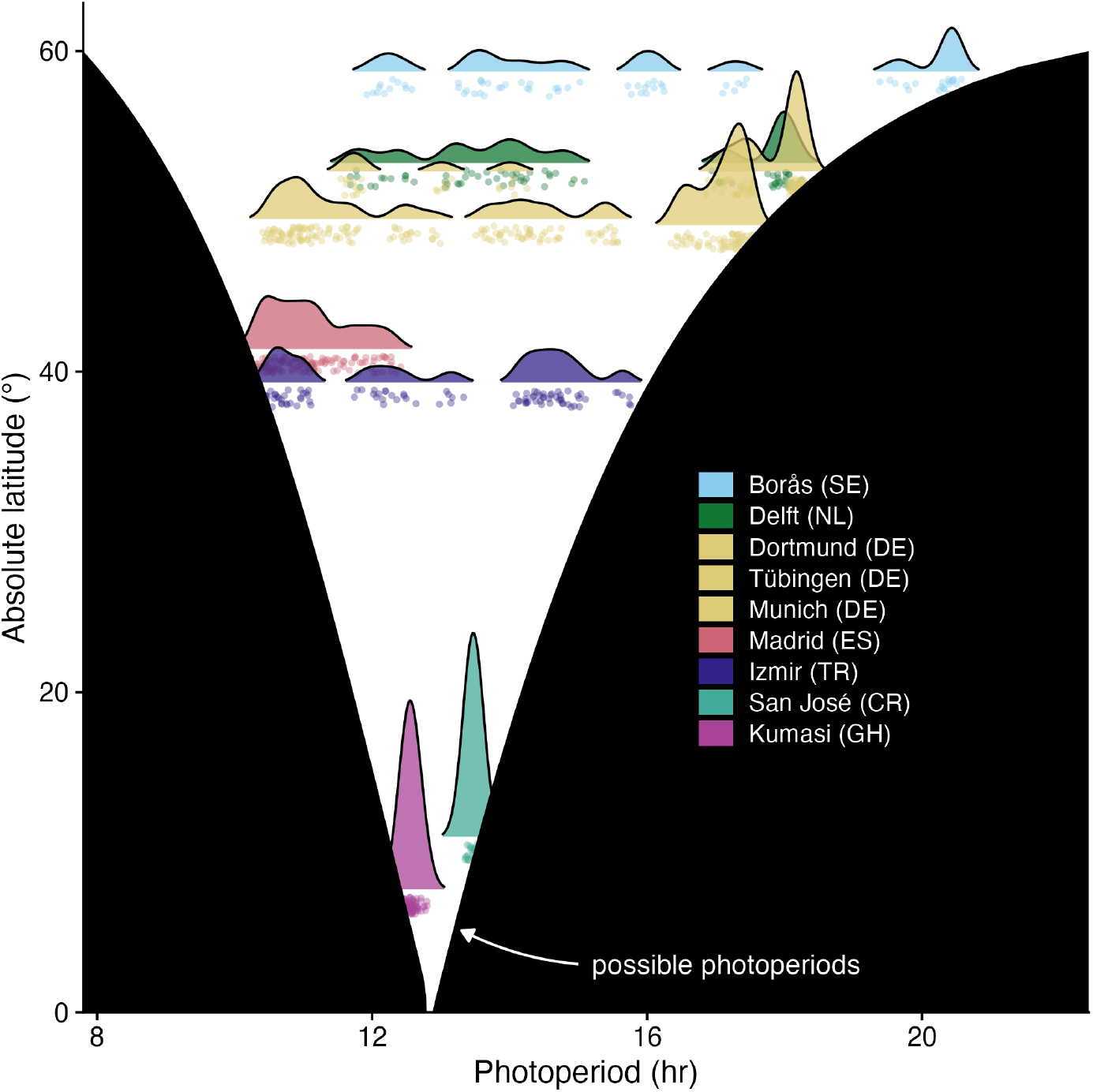
Observed civil photoperiod and theoretical bounds by absolute latitude.

**Supplementary Figure S4.**
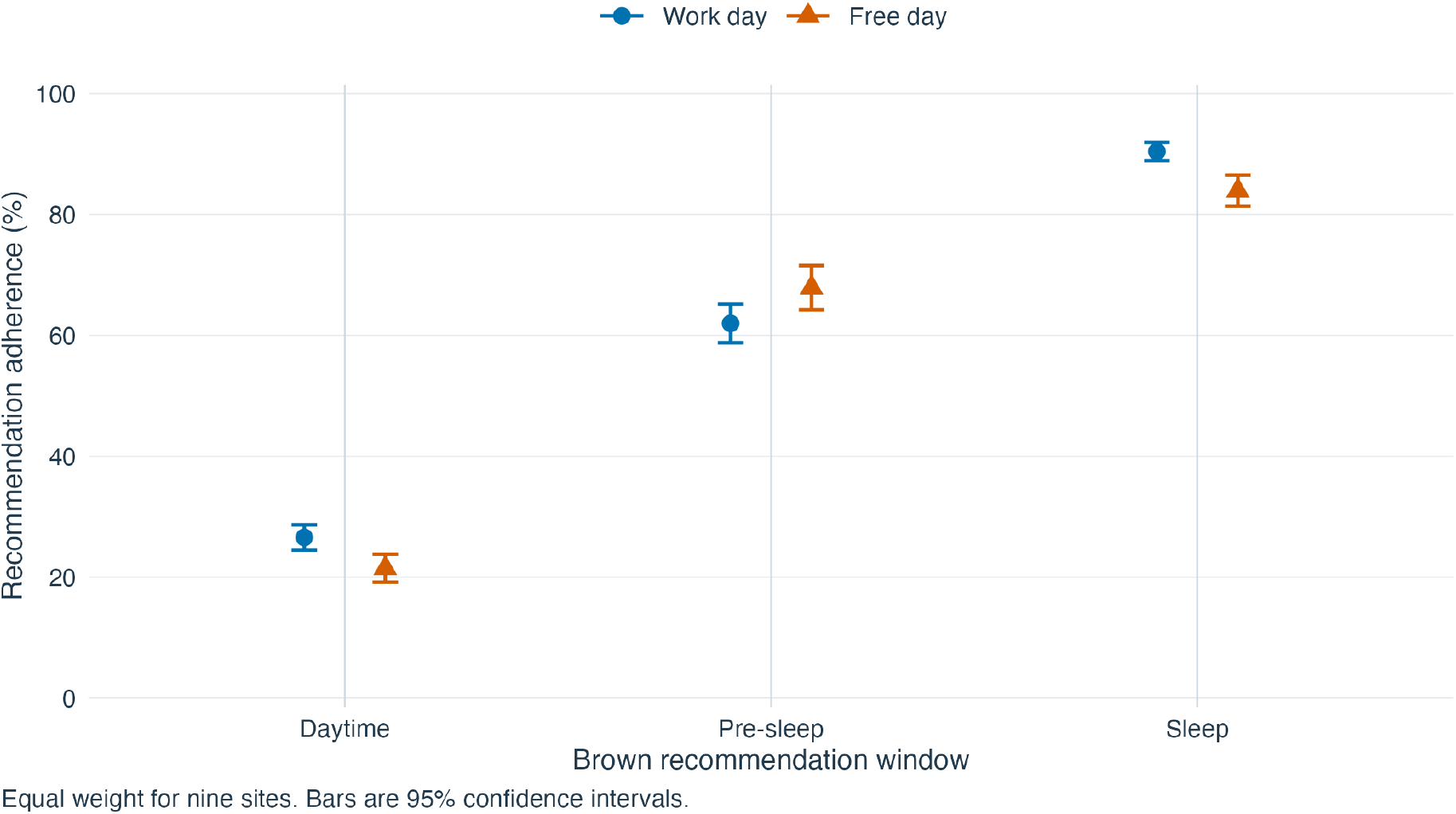
Site-average recommendation adherence on work and free days. Daytime, Pre-sleep and Sleep identify the Brown et al. recommendation windows. Each point gives equal weight to nine sites; bars are 95% confidence intervals. The wake-start date labels the preceding sleep, daytime and following pre-sleep windows. Sleep describes the device-recorded bedside environment, not ocular exposure or a whole-bedroom measurement. The pre-sleep contrast remains positive but its interval includes zero under the 80% coverage restriction; temporal dependence remains unresolved. The coverage sensitivity is reported in the Results and Methods.

**Supplementary Figure S5.**
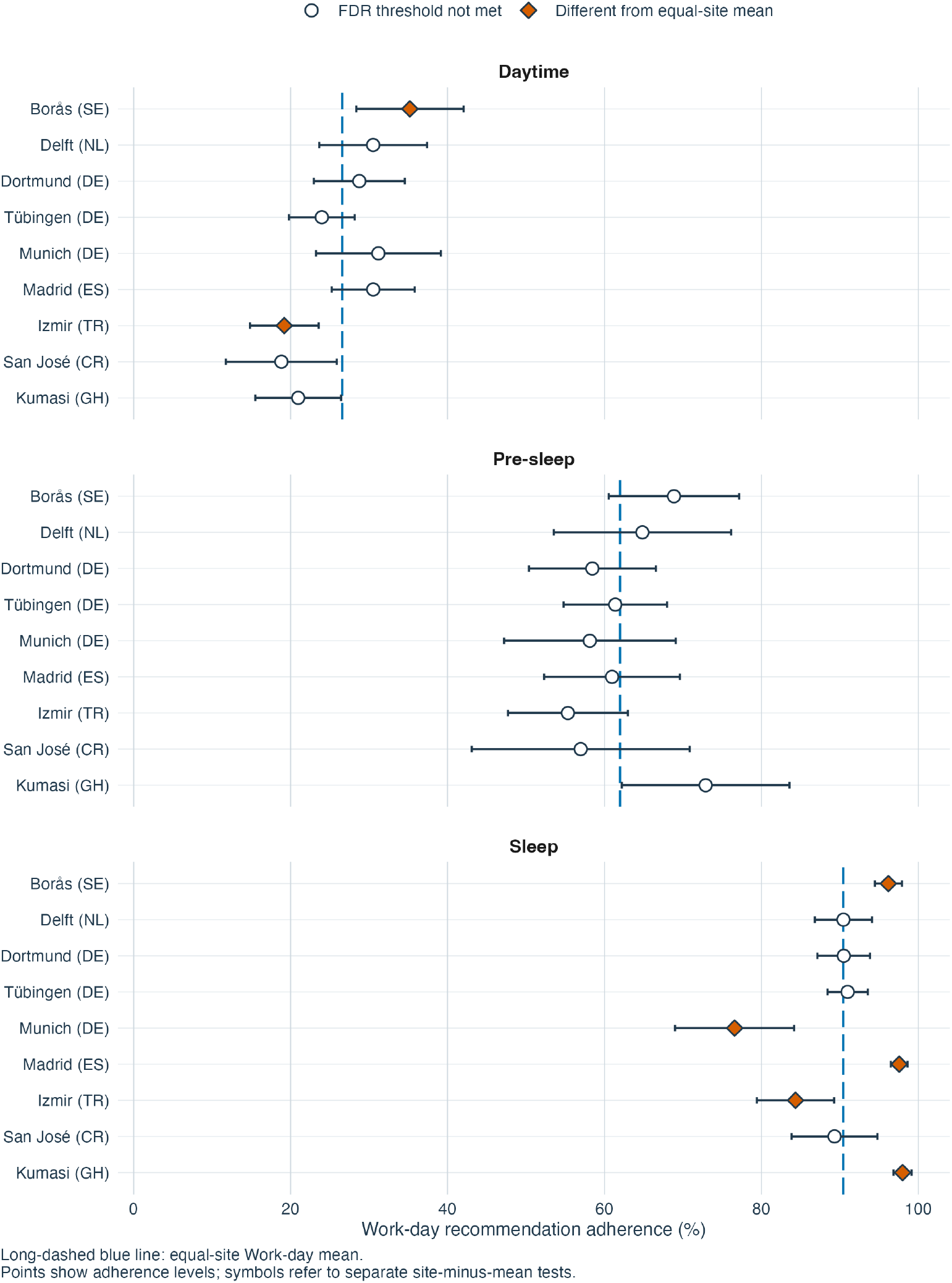
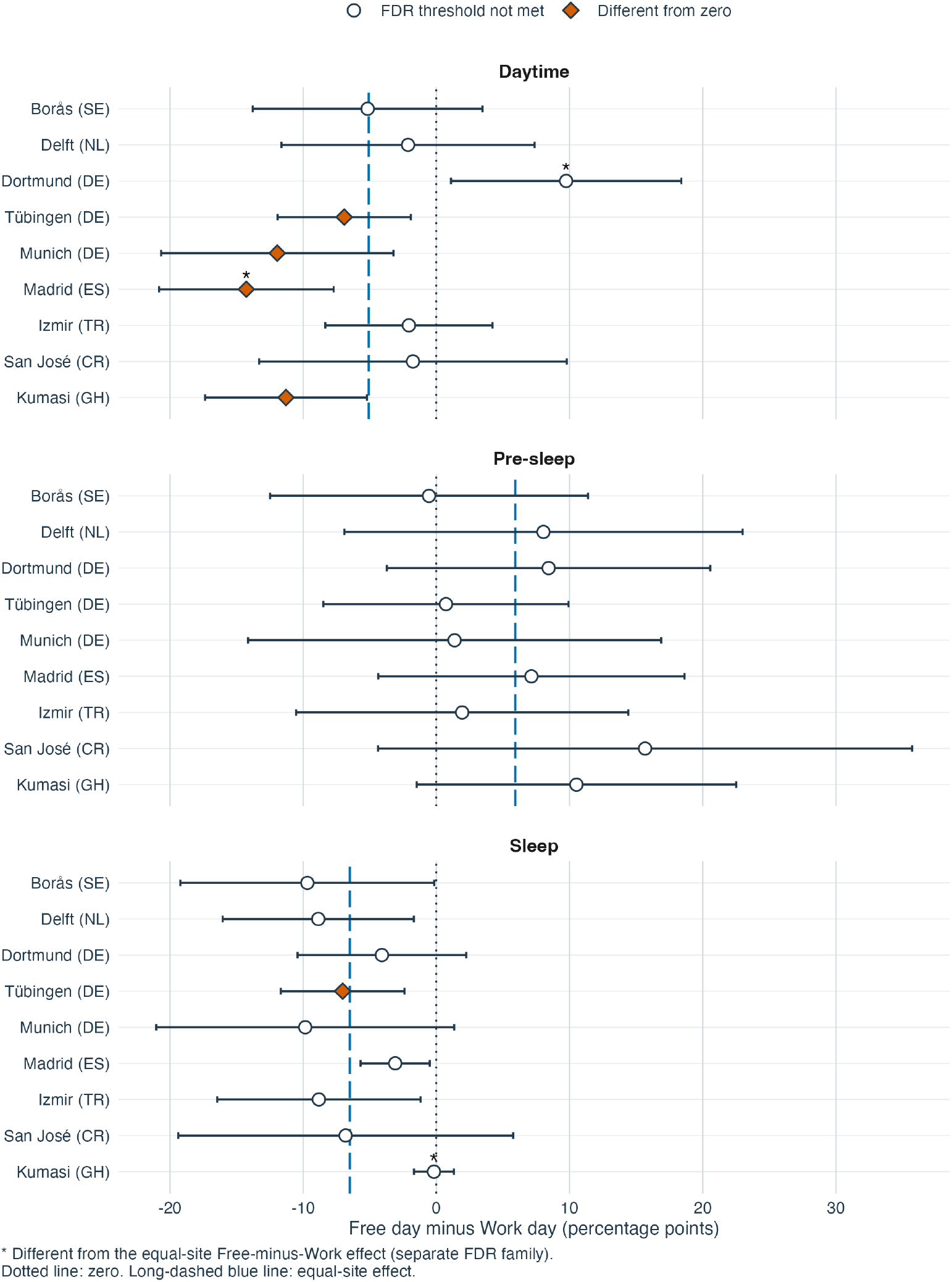
A. Site-specific work-day recommendation adherence. Filled diamonds identify site-minus-average comparisons retained after FDR adjustment; bars are 95% confidence intervals. The full explanation accompanies part B. Site-specific recommendation adherence, shown as two separate images. A, Work-day adherence, with filled diamonds identifying site-minus-average differences that met their FDR criterion. B, Free-minus-work differences, with filled diamonds identifying differences from zero and asterisks identifying departures from the window-specific equal-site difference in a separate 27-comparison FDR family. Daytime, Pre-sleep and Sleep identify the Brown et al. recommendation windows. Points are estimates and bars are 95% confidence intervals. Each equal-site reference includes the indexed site. Sites are components of one pooled observational model, not independent replications or causal effects of location. Day type follows the wake-start date; sleep is the preceding bedside sleep-environment window. Temporal dependence remains unresolved, and the pre-sleep coverage qualification applies.

**Supplementary Figure S6.**
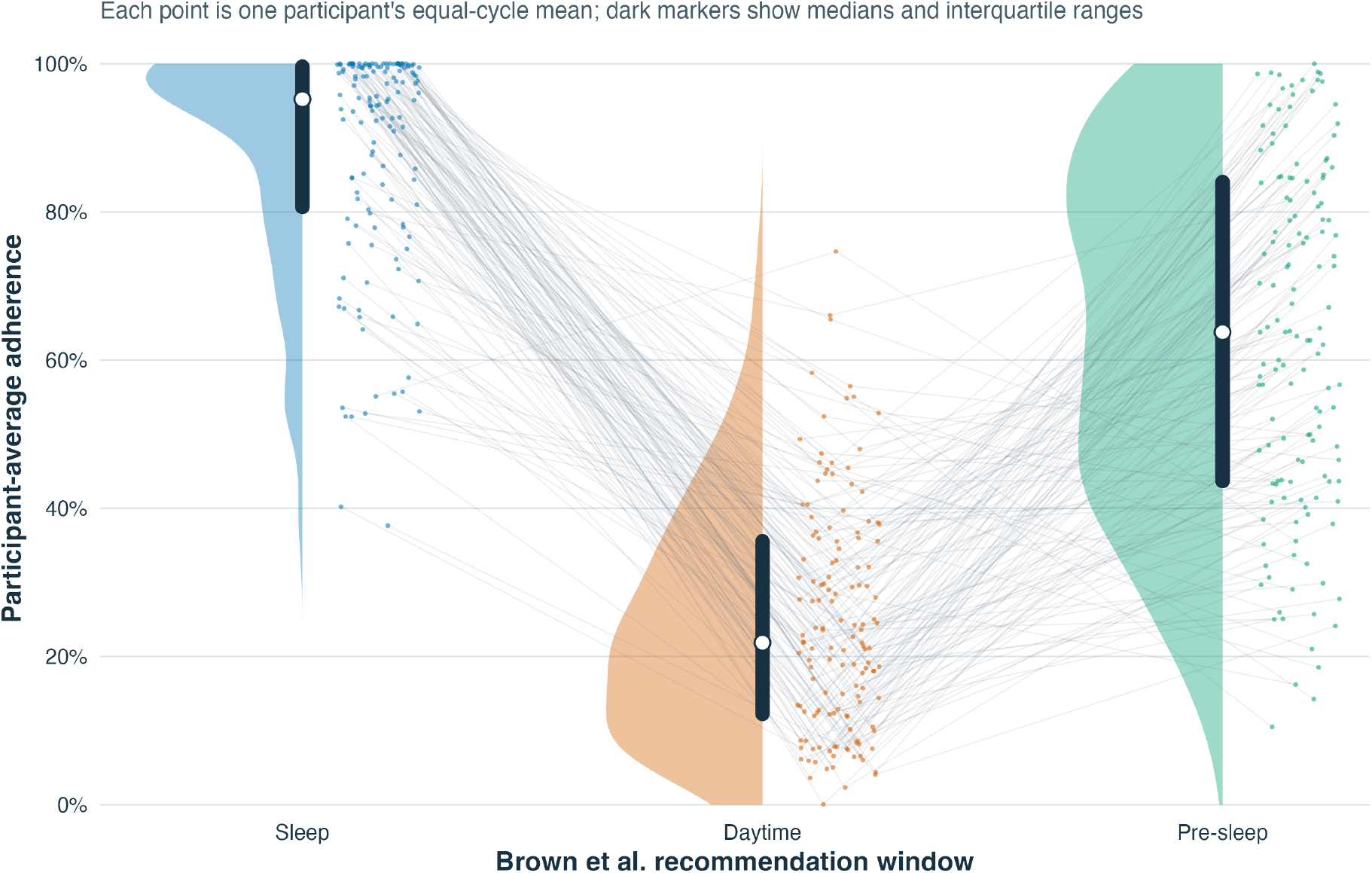
Participant-average recommendation adherence across the Brown et al. recommendation windows. Daytime, Pre-sleep, and Sleep are the window labels. Each colored point is one anonymous participant’s equal-cycle mean within that window. Faint lines connect the same anonymous participant across the three window summaries; they are not time trajectories or ranks. Half-violin shapes show the window distributions, and dark bars with white points show interquartile ranges and medians. Because each window has its own Brown et al. threshold and direction, absolute percentages should be interpreted within window.

**Supplementary Figure S7.**
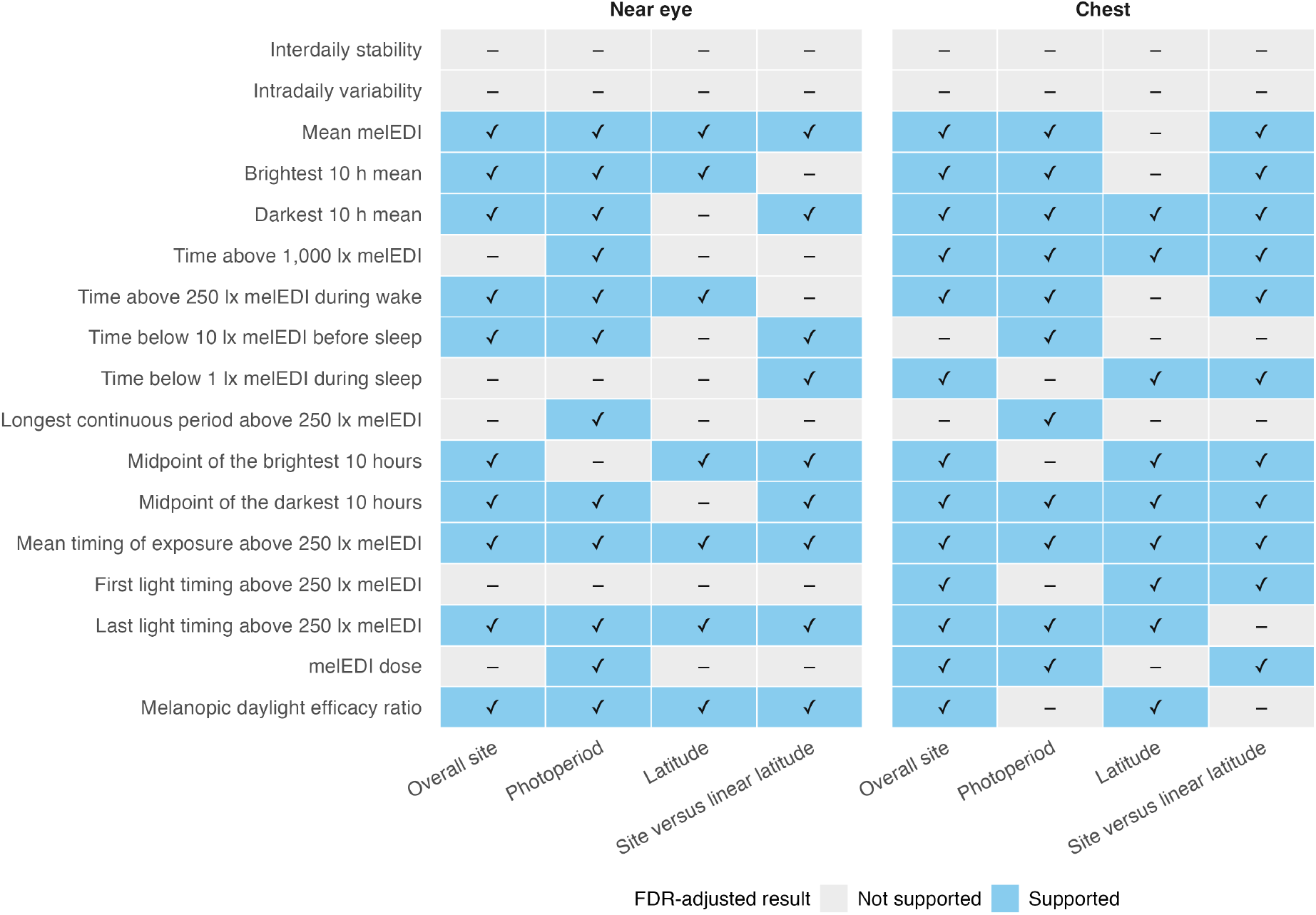
Geographic and civil-photoperiod associations across near-eye personal light-exposure metrics. Evidence retained after FDR adjustment for overall site, civil photoperiod, latitude, and site-versus-linear-latitude adequacy. These descriptive associations do not identify a mechanism.

**Supplementary Figure S8.**
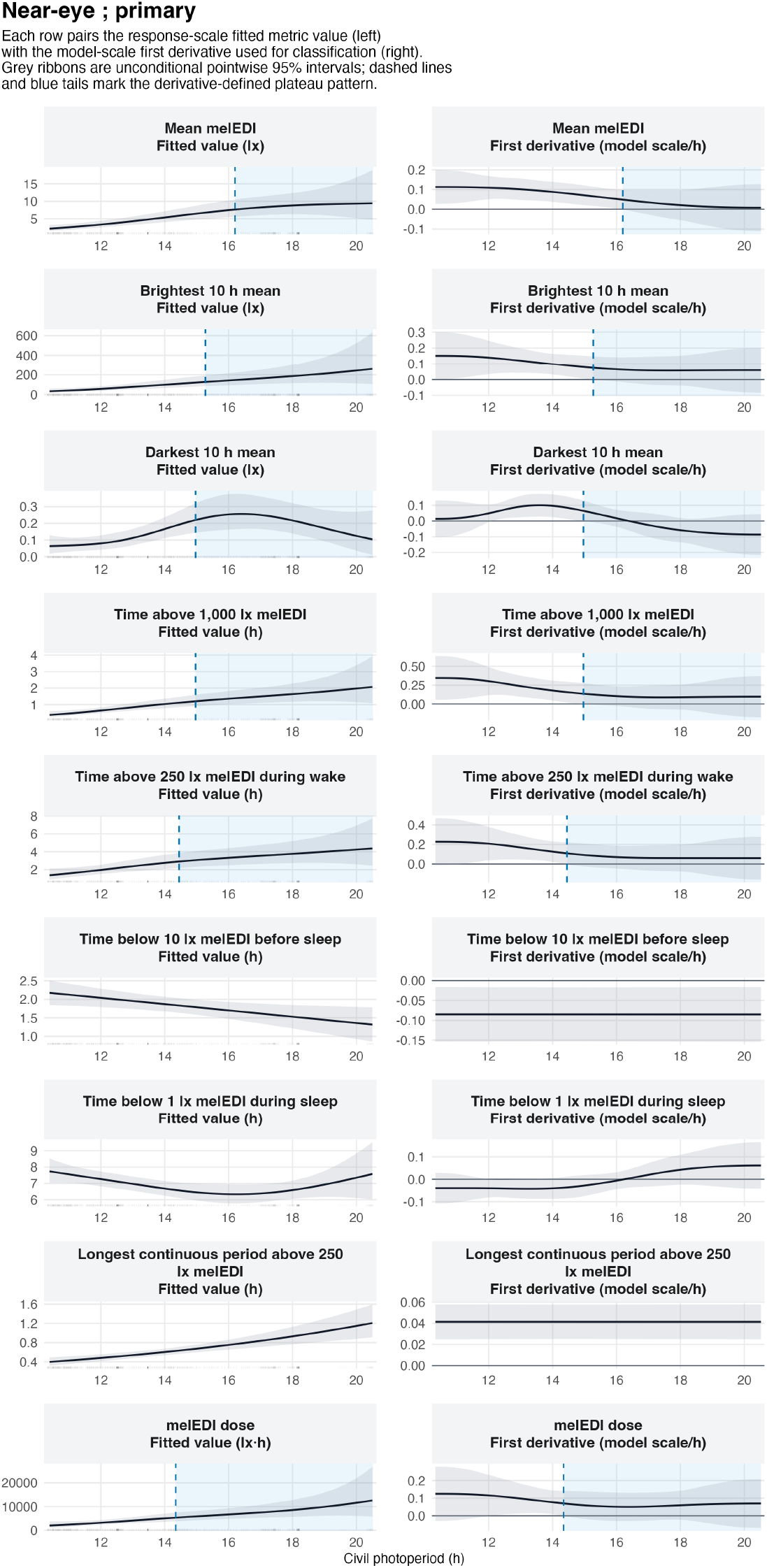
Nonlinear civil-photoperiod associations and their fitted slopes. A qualifying transition denotes an increase followed by a sustained near-flat tail under the specified slope rule. These descriptive associations do not identify a mechanism, and no physiological or environmental ceiling was identified.

**Supplementary Figure S9.**
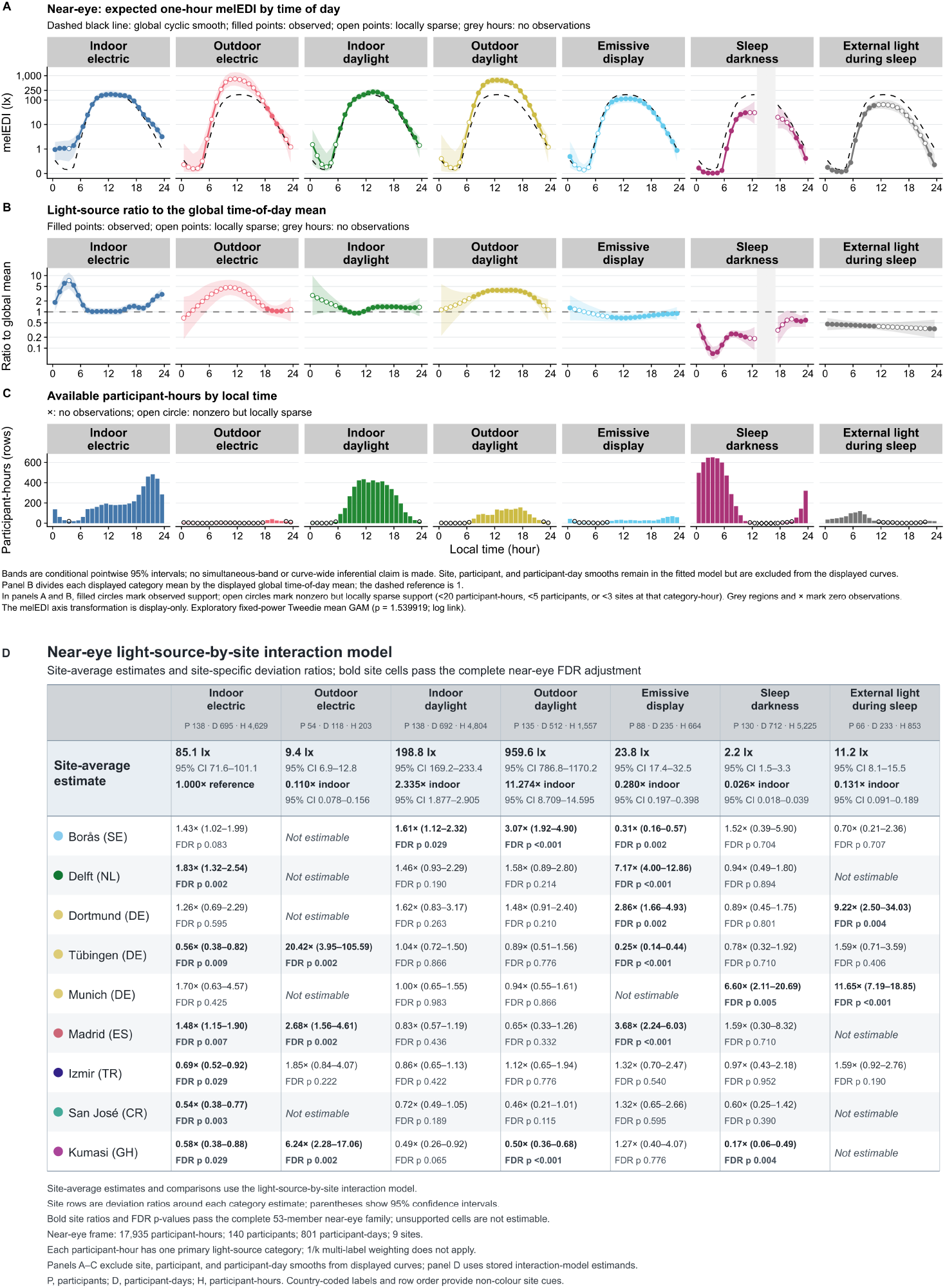
Near-eye light exposure by reported light source across time of day and study site. A–C, Exploratory time-of-day analysis. Panel A shows expected one-hour melanopic EDI with clock-specific 95% confidence intervals; the dashed line is the site-average local-clock smooth. Panel B shows each light-source curve relative to that smooth, with 1 as the reference. Panel C shows available participant-hours; open circles indicate locally sparse support and grey gaps indicate no observations. D, Site-average category estimates and site-specific deviation ratios from the light-source-by-site interaction model, with participant-cluster-robust 95% confidence intervals, FDR-adjusted p values, and exact support. The analysis includes 17,935 participant-hours from 140 participants, covering 801 participant-days and nine sites. Panels A–C and panel D answer different questions and are not numerically interchangeable. Estimates are observational. During reported sleep, measurements describe the bedside sleep environment.

**Supplementary Figure S10.**
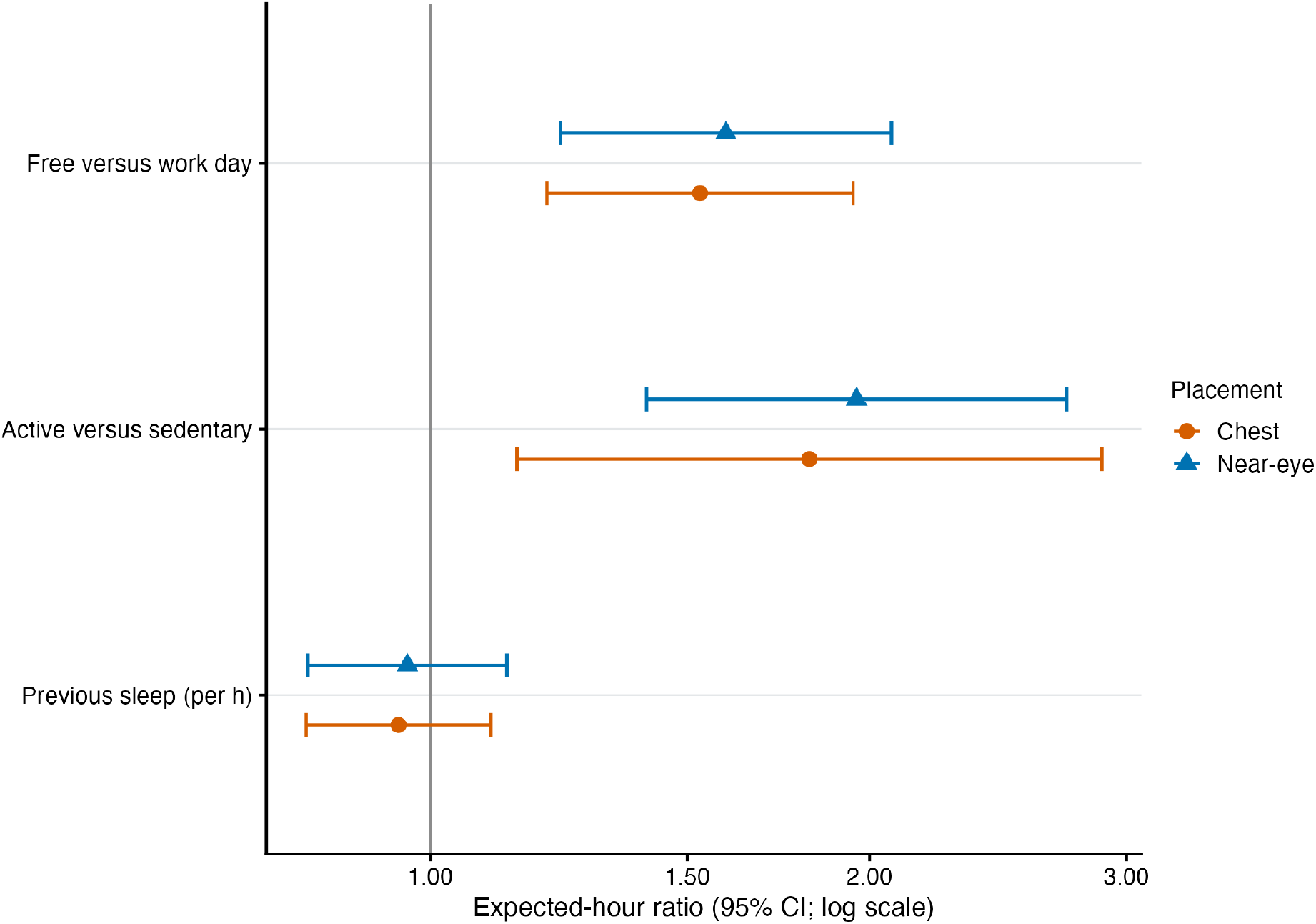
Near-eye and complementary chest associations estimated separately for the same participants and participant-days. The sets of supported participant-hours differ between sensor positions, so the display is not an observation-level identity comparison and does not test equivalence.

**Supplementary Figure S11.**
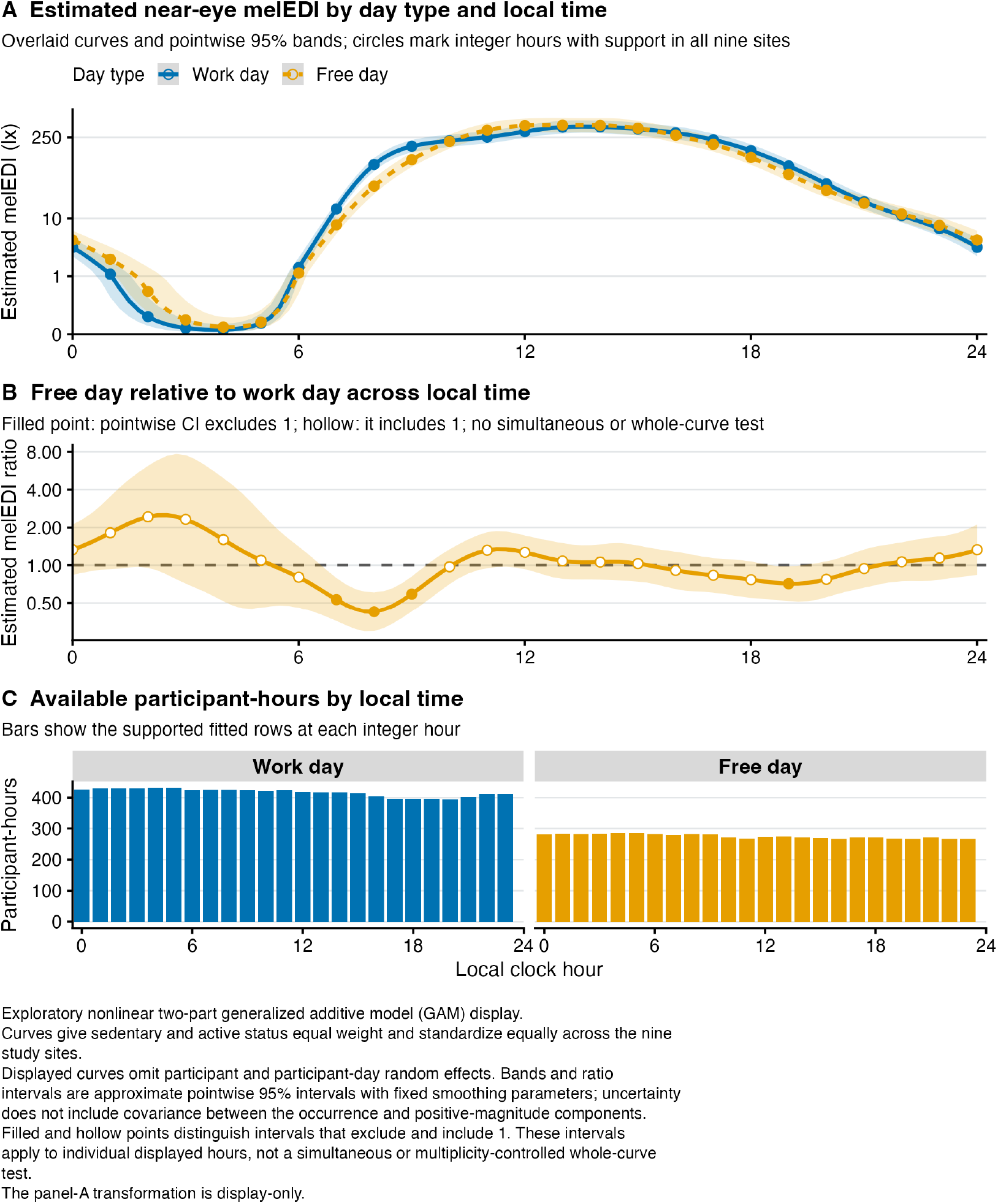
Exploratory Free-versus-Work hourly fitted exposure and response-scale ratio curves with 95% confidence intervals and participant-hour support.

**Supplementary Figure S12.**
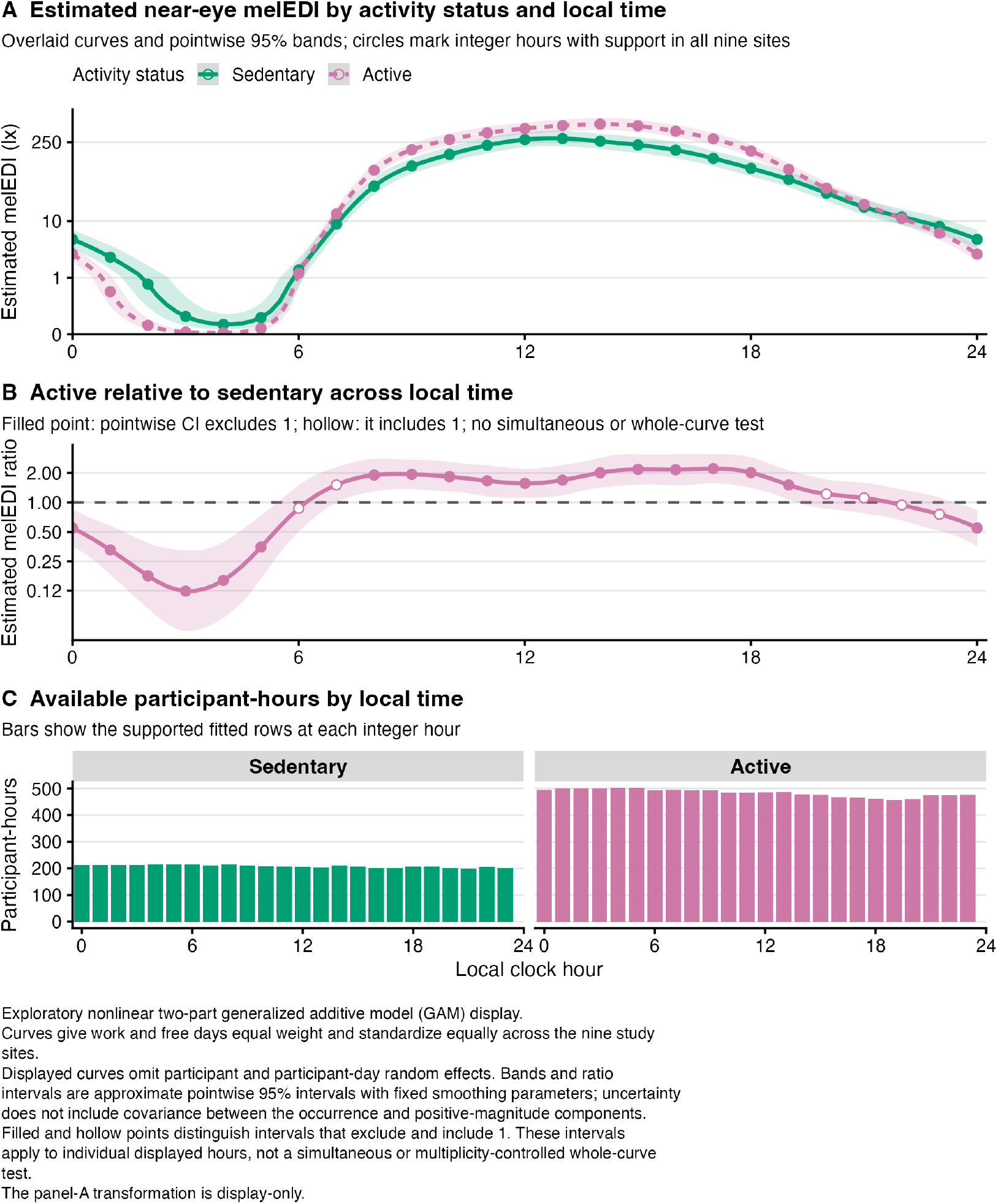
Exploratory Active-versus-Sedentary hourly fitted exposure and response-scale ratio curves with 95% confidence intervals and participant-hour support.

**Supplementary Figure S13.**
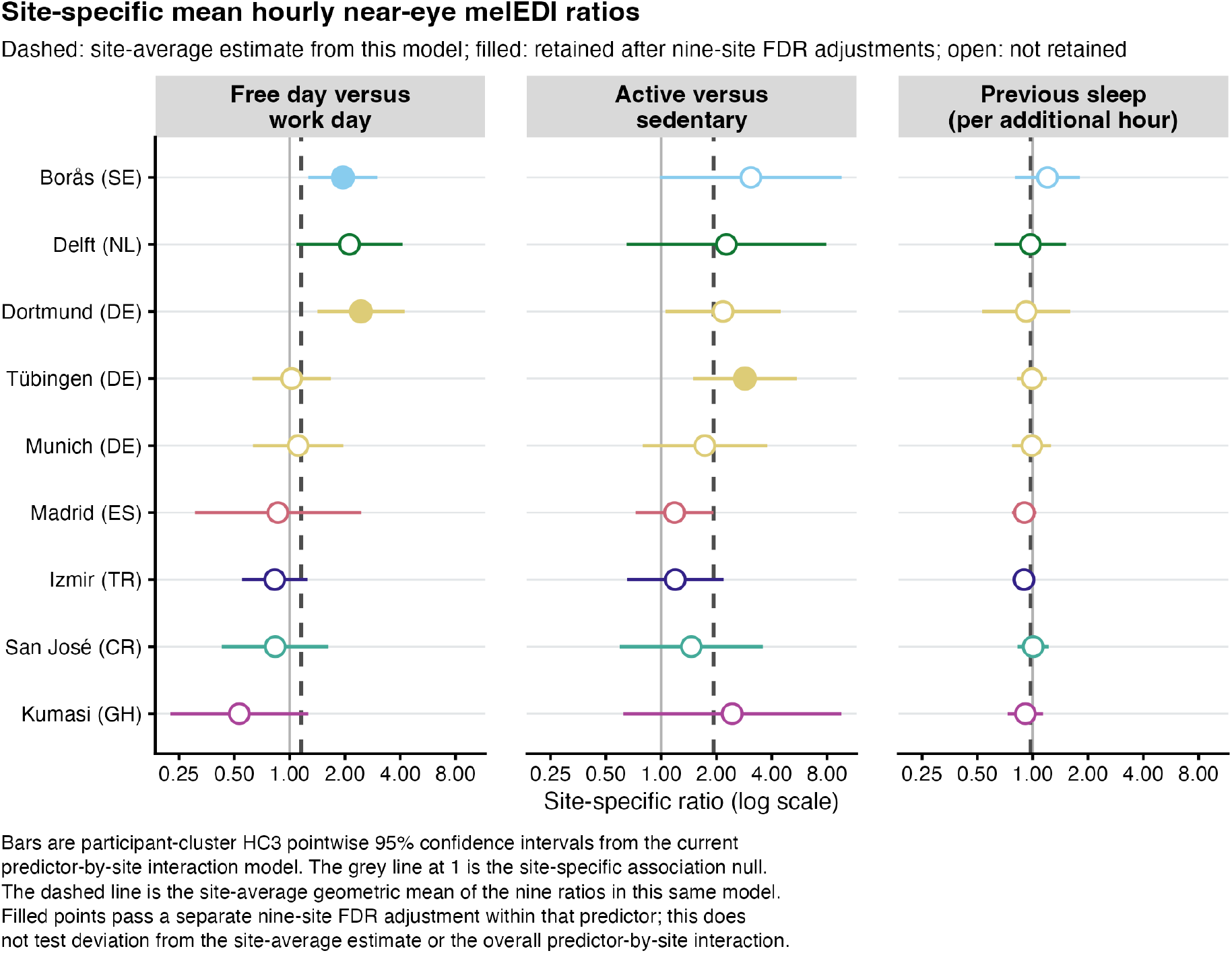
Site-specific estimates from the predictor-by-site interaction model. Filled and hollow points distinguish associations that did and did not meet the 0.050 criterion after a separate nine-site FDR adjustment within each predictor. The 95% confidence intervals are unadjusted; dashed lines show site-average estimates calculated with equal weight for each of the nine sites. Sites are components of one pooled model, not independent replications or causal effects of location.

**Supplementary Figure S14.**
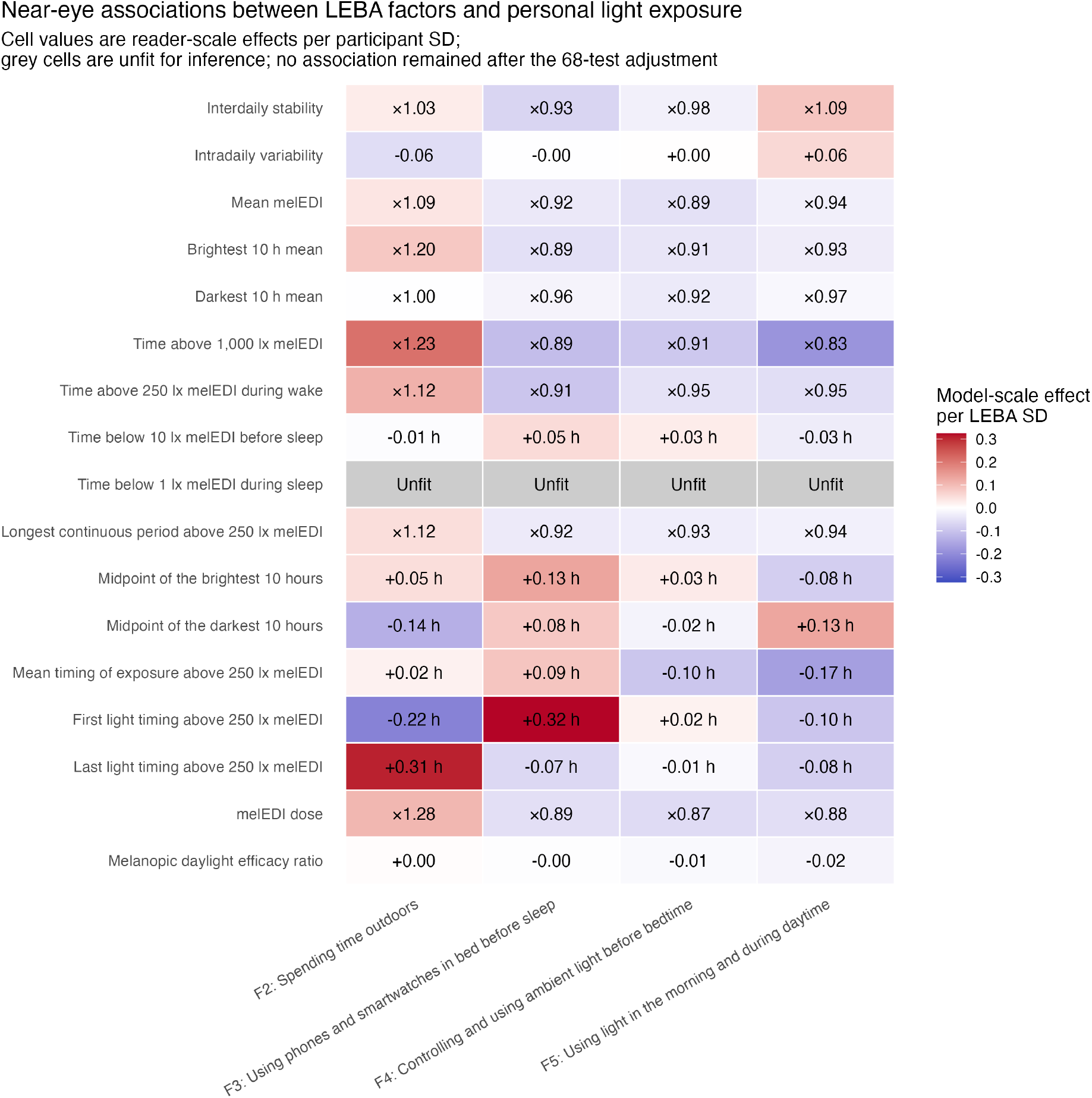
Near-eye associations per one participant-level standard deviation of each Light Exposure Behaviour Assessment factor. None of the 68 tests retained FDR-adjusted support. Four sleep-environment cells were unfit for inference and are labelled Unfit with estimates suppressed. Lack of adjusted support is inconclusive rather than proof of no association.

**Supplementary Figure S15.**
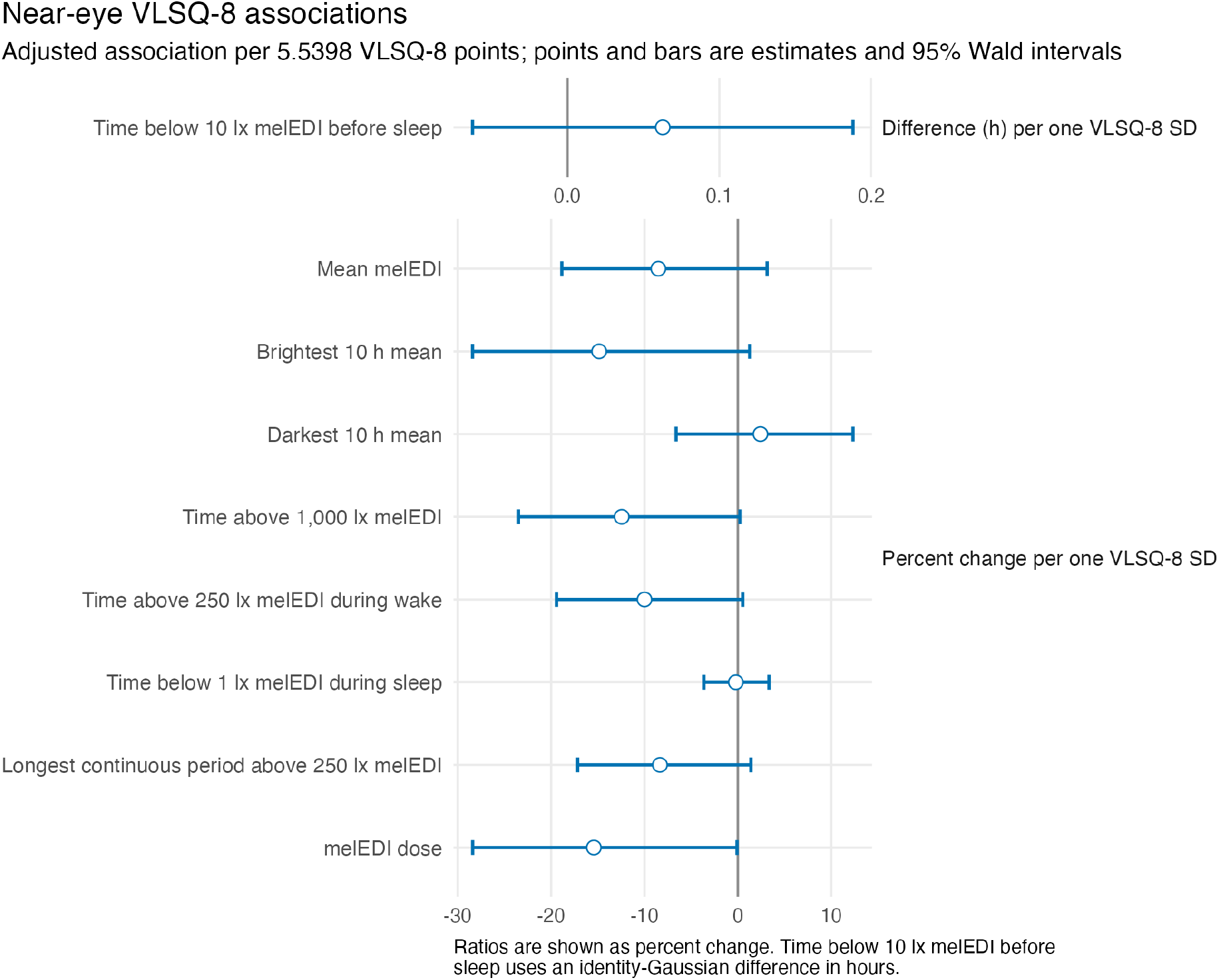
Near-eye site-average associations between the Visual Light Sensitivity Questionnaire-8 and nine personal light-exposure metrics. Associations compare scores separated by one participant-level standard deviation. Horizontal bars are 95% Wald confidence intervals. None of the nine associations retained FDR-adjusted support.

**Supplementary Figure S16.**
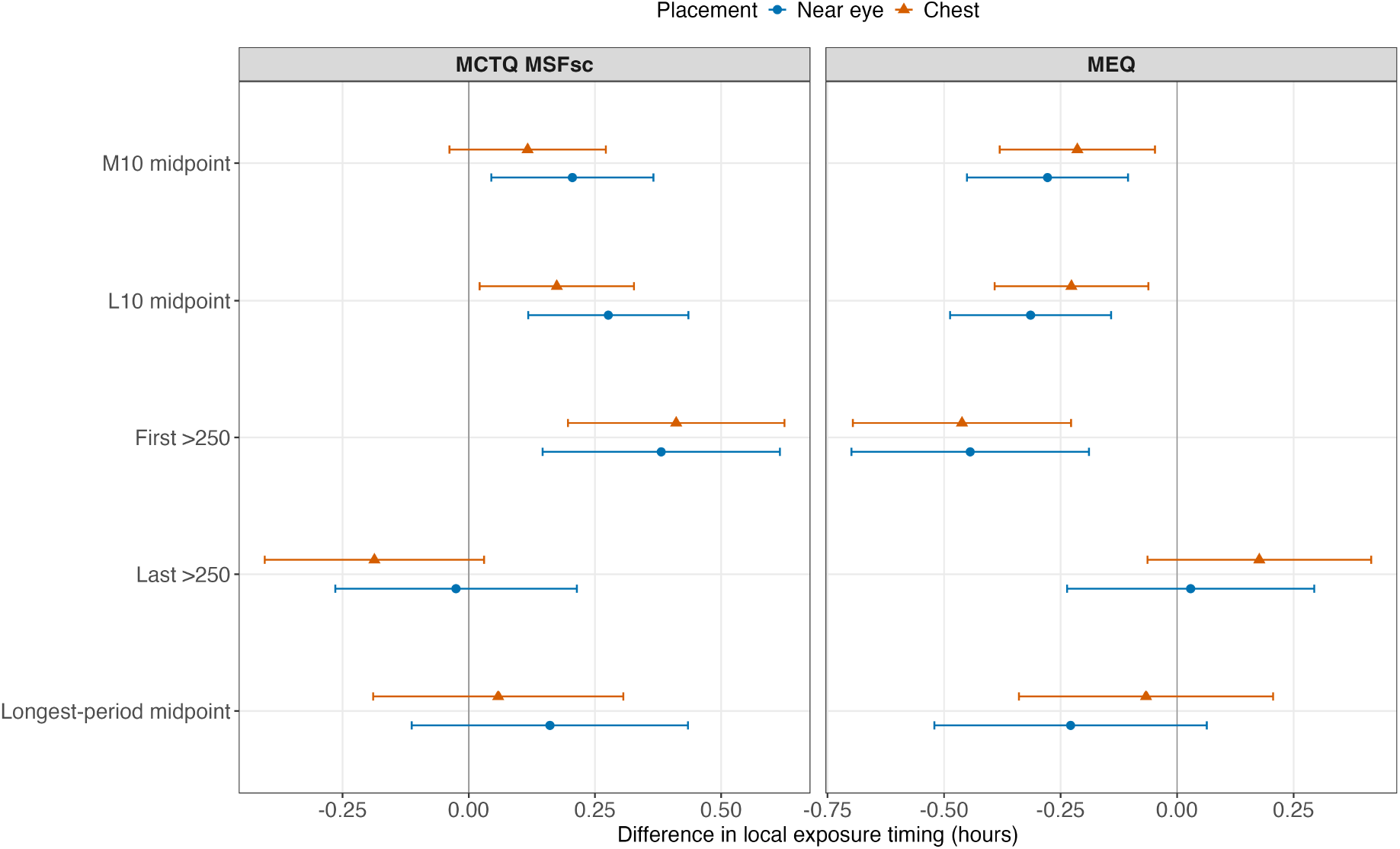
Chronotype and timing of personal light exposure. Study-site-adjusted associations of sleep-timing-based corrected midsleep on Free days and questionnaire-based morningness-eveningness with the analysed timing metrics. Points are estimates and bars are 95% confidence intervals; the two chronotype instruments remain separate. Near-eye estimates provide ocular-exposure evidence; chest estimates, where shown, are complementary non-ocular evidence.

**Supplementary Figure S17.**
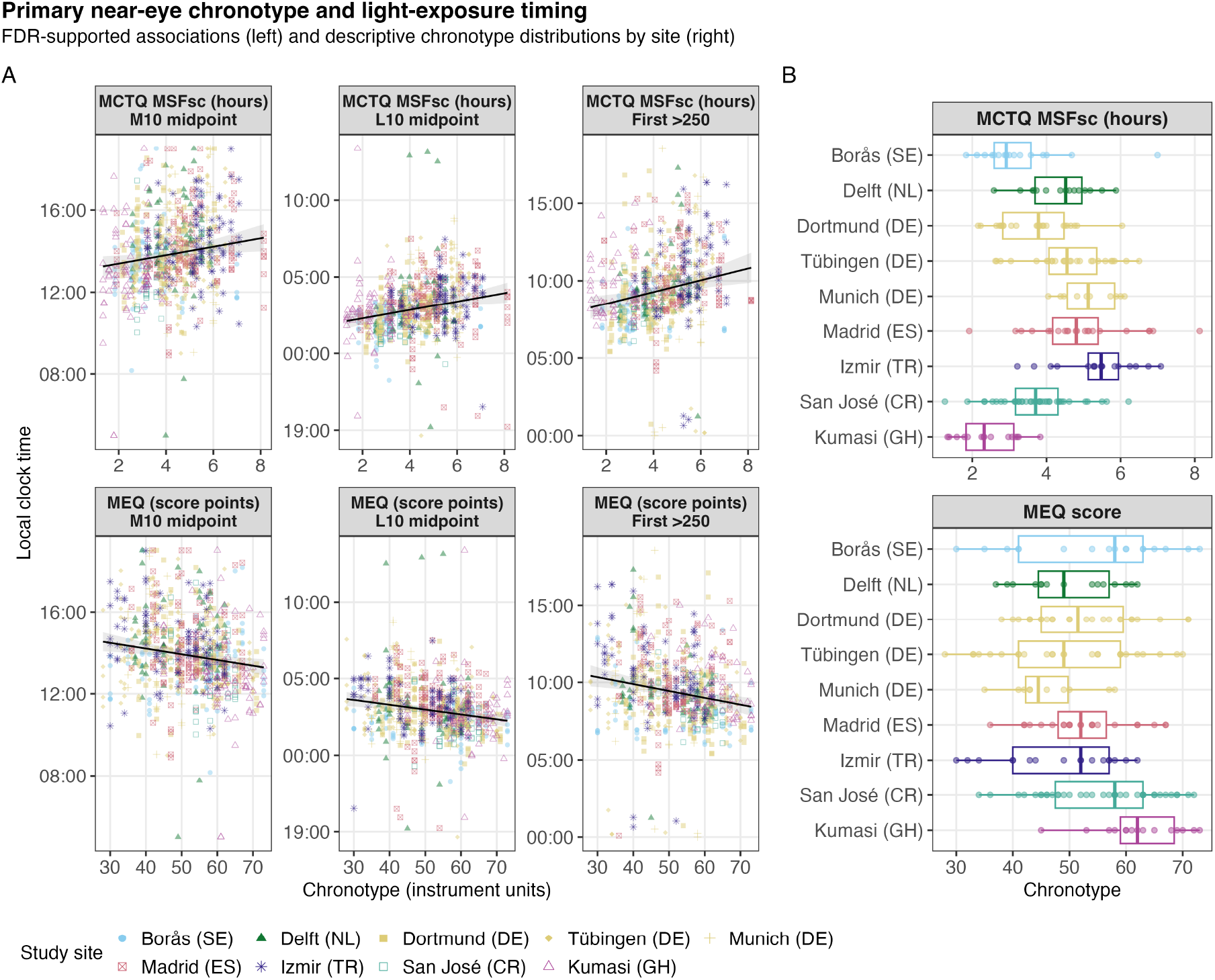
Observed timing across chronotype and study sites. Observed participant-day timing values across chronotype and country-coded study sites for the samples used in the corresponding models. Each point represents one participant-day in a fitted sample. This figure is descriptive and does not replace the adjusted models or estimate independent site effects. Near-eye estimates provide ocular-exposure evidence; chest estimates, where shown, are complementary non-ocular evidence.

**Supplementary Figure S18.**
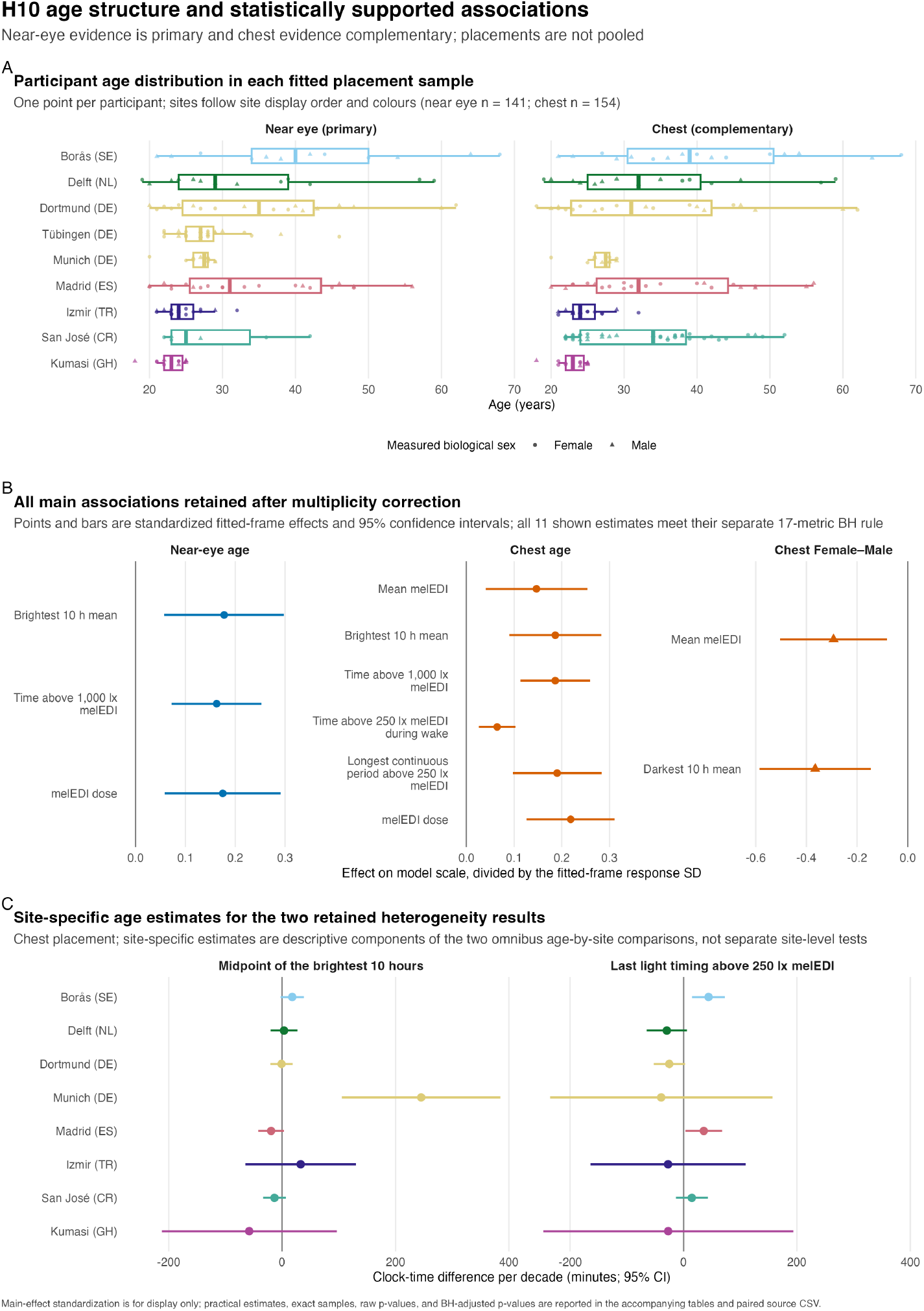
Participant age distributions, the 11 main associations retained after FDR adjustment, and descriptive site-specific estimates for two retained age-by-site interactions in the complementary chest analysis. The site-specific interaction estimates are descriptive model contrasts, not independent site tests. Biological sex, coded Female or Male, was analysed; gender was recorded separately and was not analysed.

**Supplementary Figure S19.**
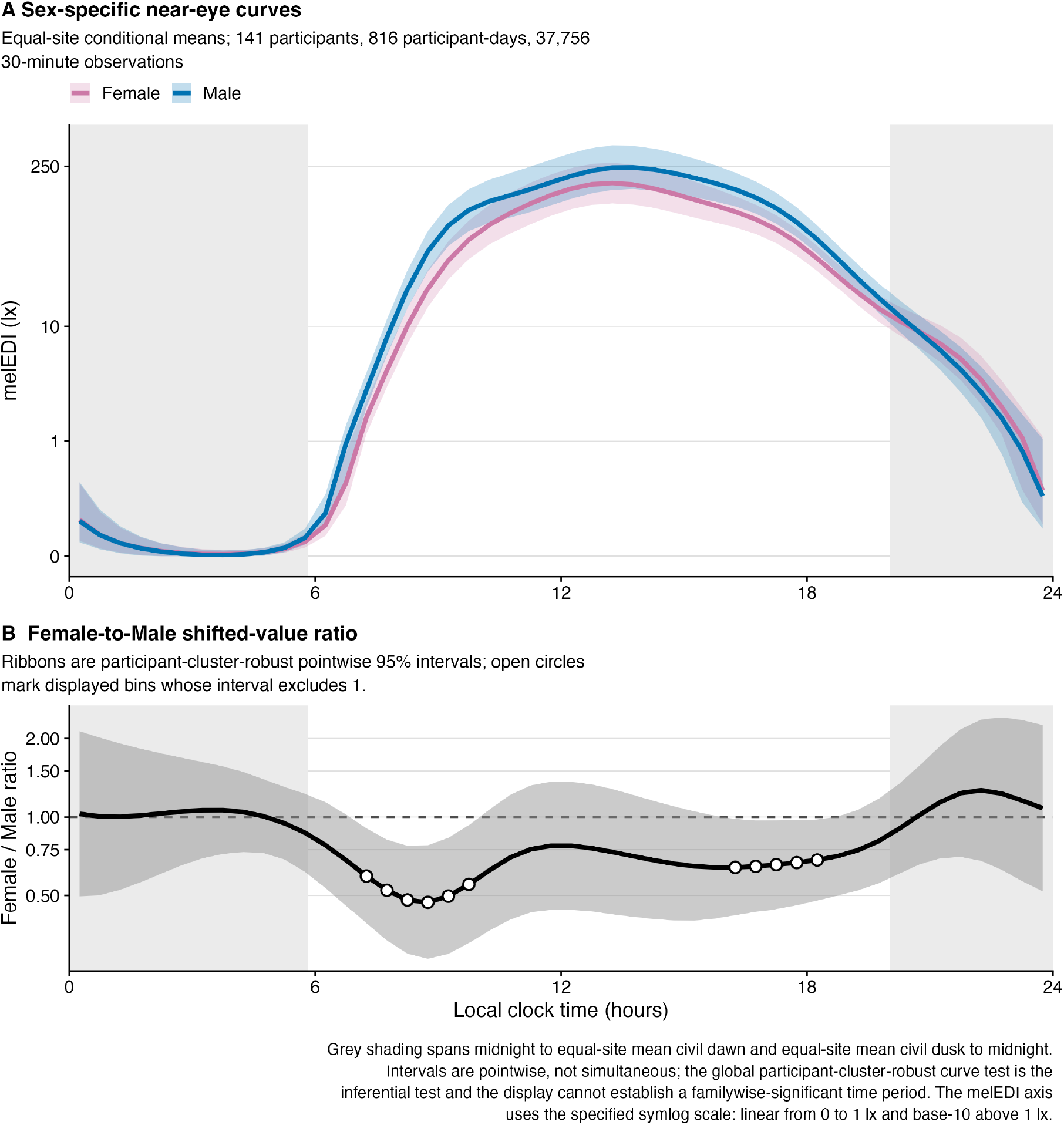
Near-eye biological-sex-specific fitted melanopic equivalent daylight illuminance curves and Female-to-Male shifted-value ratio. Ribbons and contrast intervals are participant-cluster-robust 95% confidence intervals at the displayed clock times. Grey shading gives site-average civil-night context with each site weighted equally and is not a model covariate. Biological sex, coded Female or Male, was analysed; gender was recorded separately and was not analysed.

## Supplementary tables

**Supplementary Table S1.** Participant, participant-day, and one-minute observation counts across the roster, sensor-specific completeness and all-zero screens, final descriptive datasets, and paired main subset. Each row names its denominator explicitly. An excluded participant-day is not necessarily an excluded participant.

|  | Participants | Participant-days | One-minute real observations |
| --- | --- | --- | --- |
| participant roster |  |  |  |
| Available normalized participant metadata | 191 | Not applicable | Not applicable |
| near_eye |  |  |  |
| At least 80% complete before all-zero screen | Not applicable | 818 | Not applicable |
| Exact all-zero days excluded | Not applicable | 2 | Not applicable |
| Main dataset after all-zero screen | 141 | 816 | 1,175,160 |
| chest |  |  |  |
| At least 80% complete before all-zero screen | Not applicable | 905 | Not applicable |
| Exact all-zero days excluded | Not applicable | 3 | Not applicable |
| Main dataset after all-zero screen | 154 | 902 | 1,298,880 |
| paired |  |  |  |
| Paired main subset | 112 | 643 | Not applicable |

**Supplementary Table S2.**
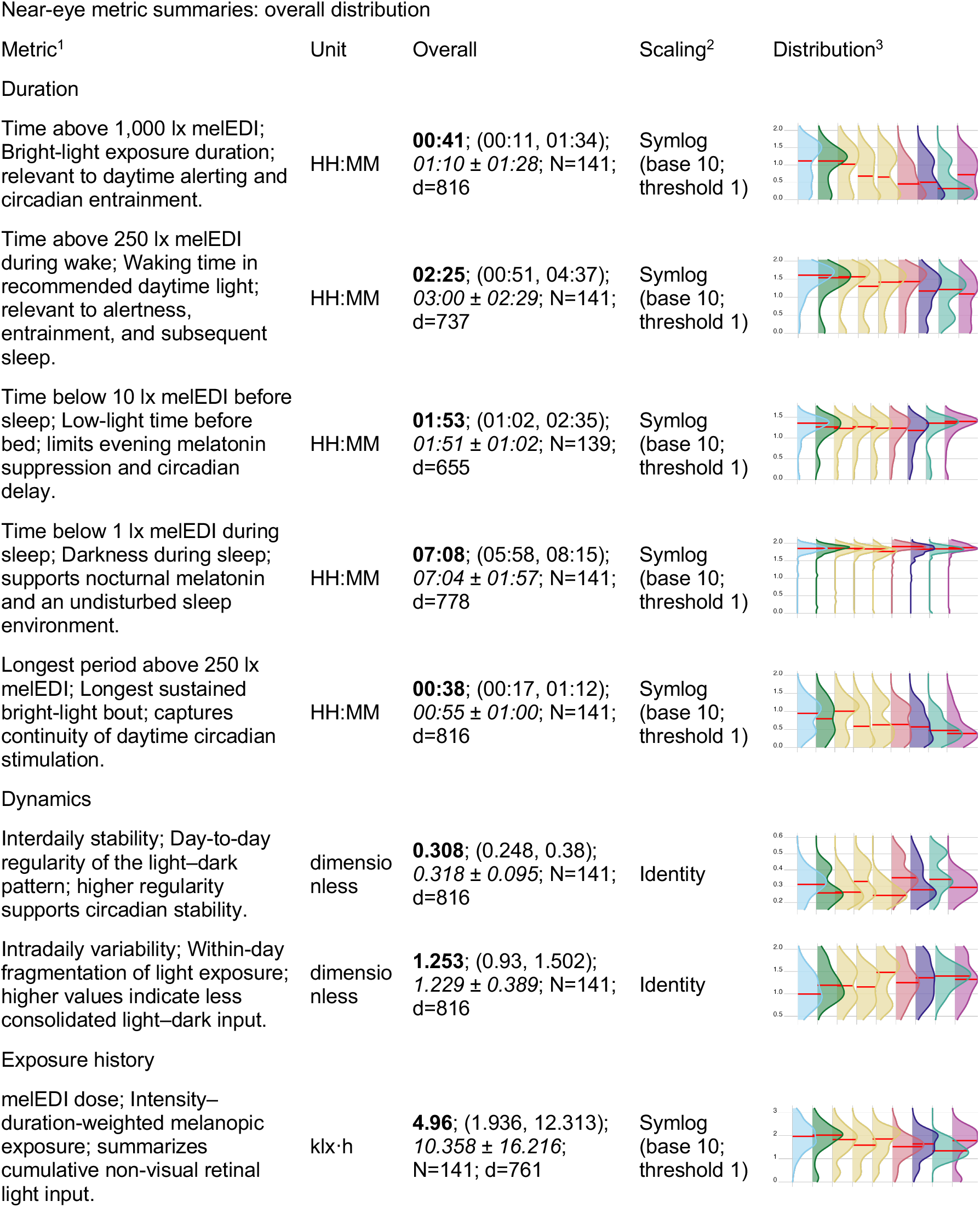
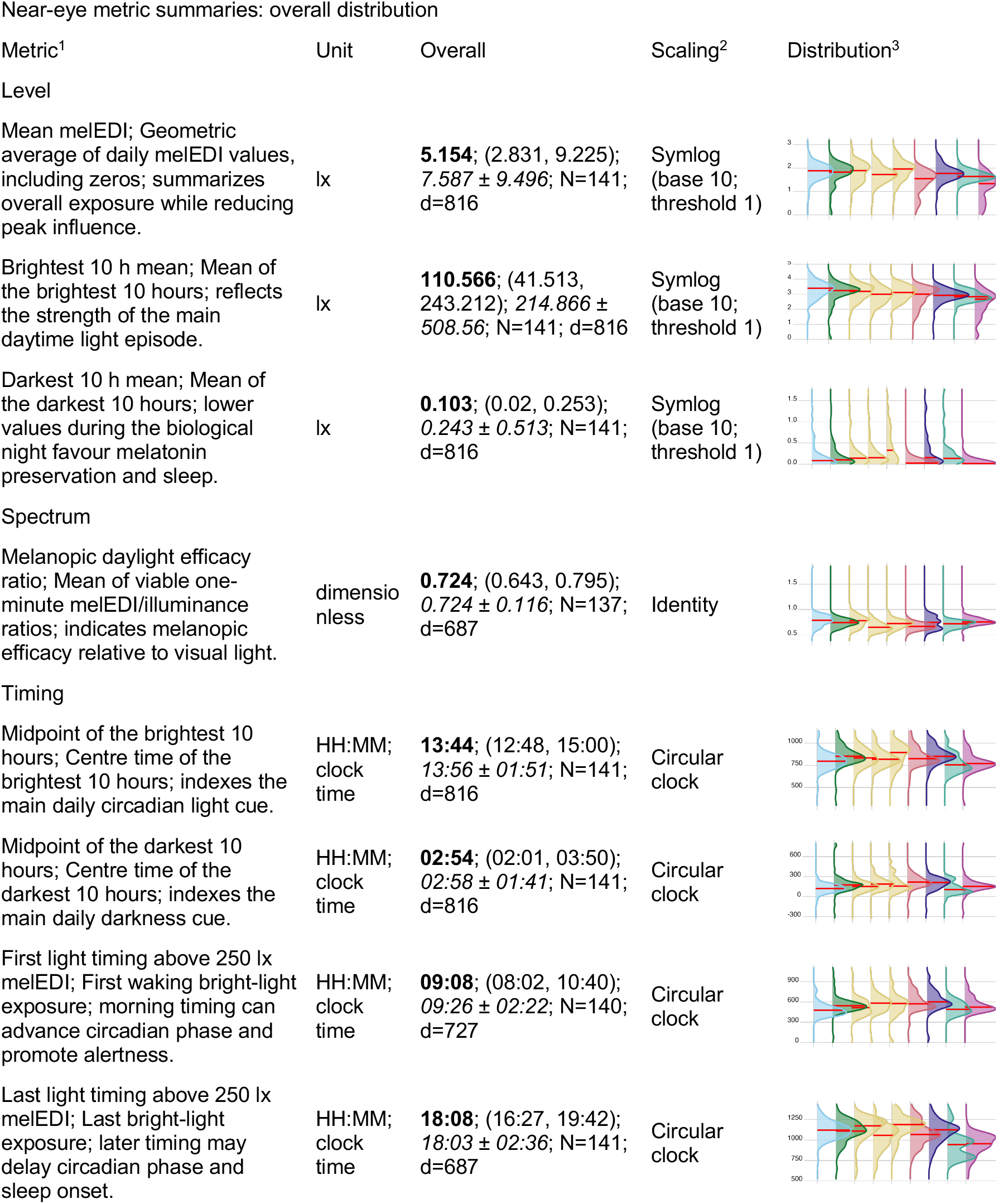
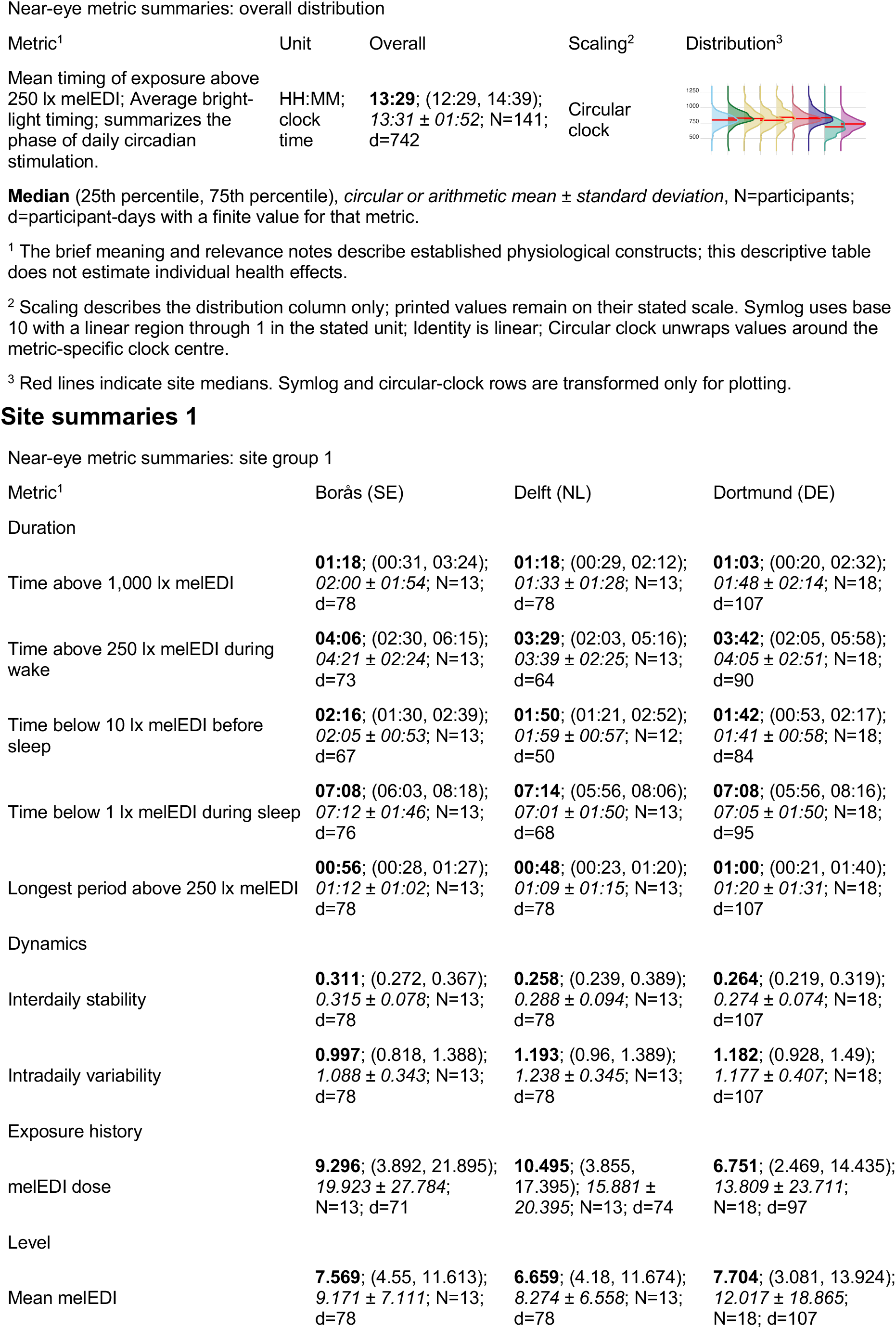
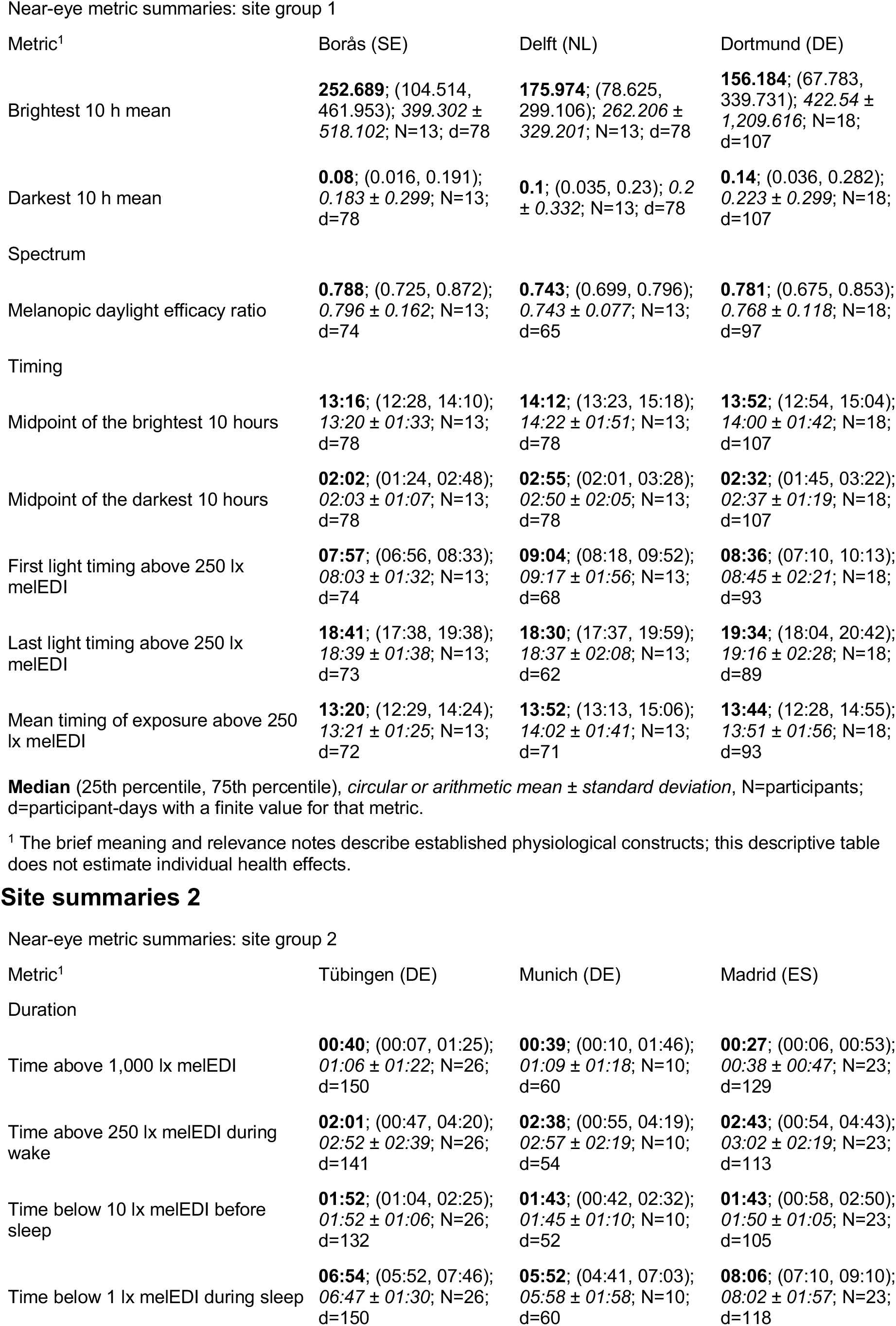
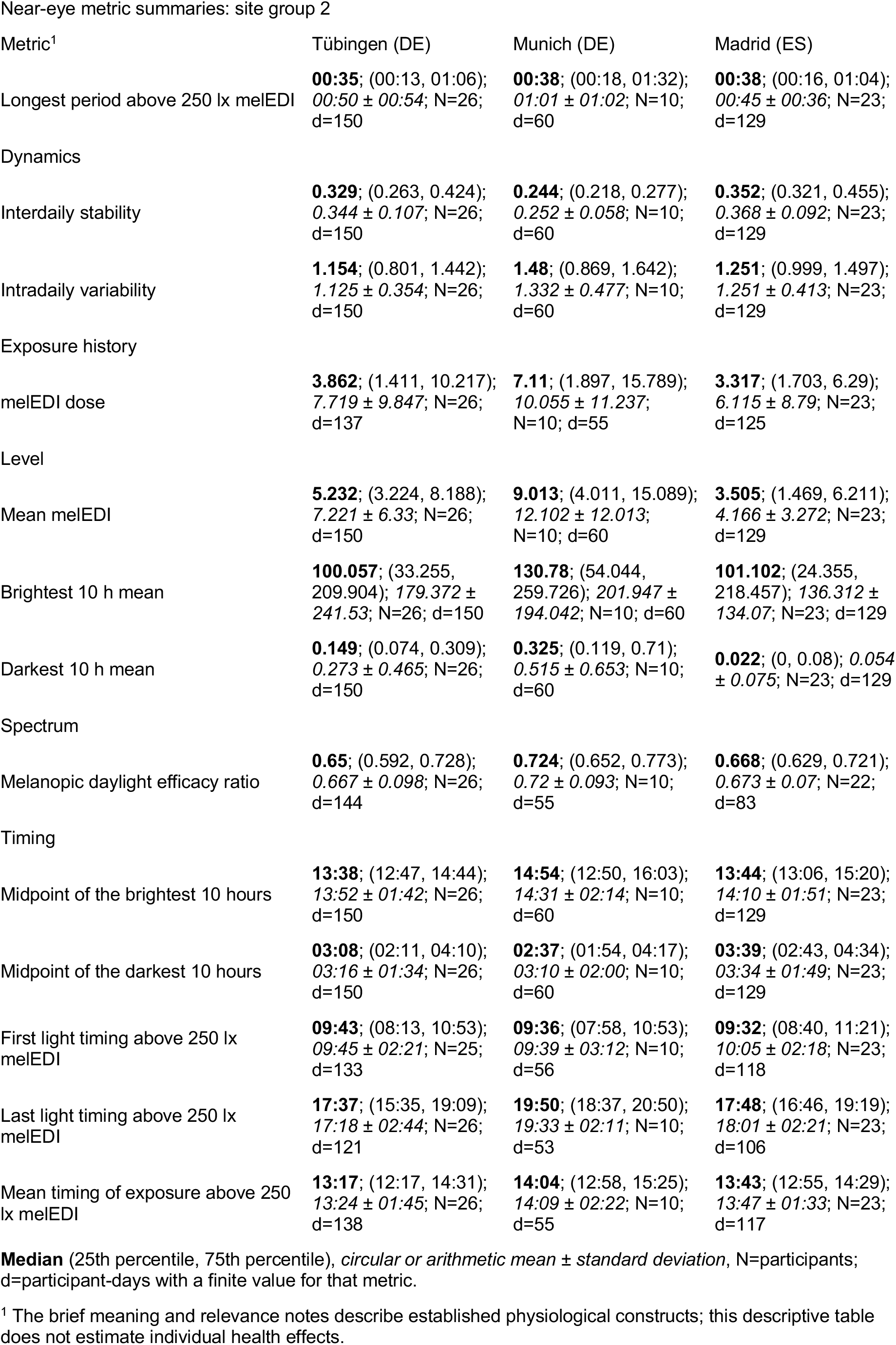
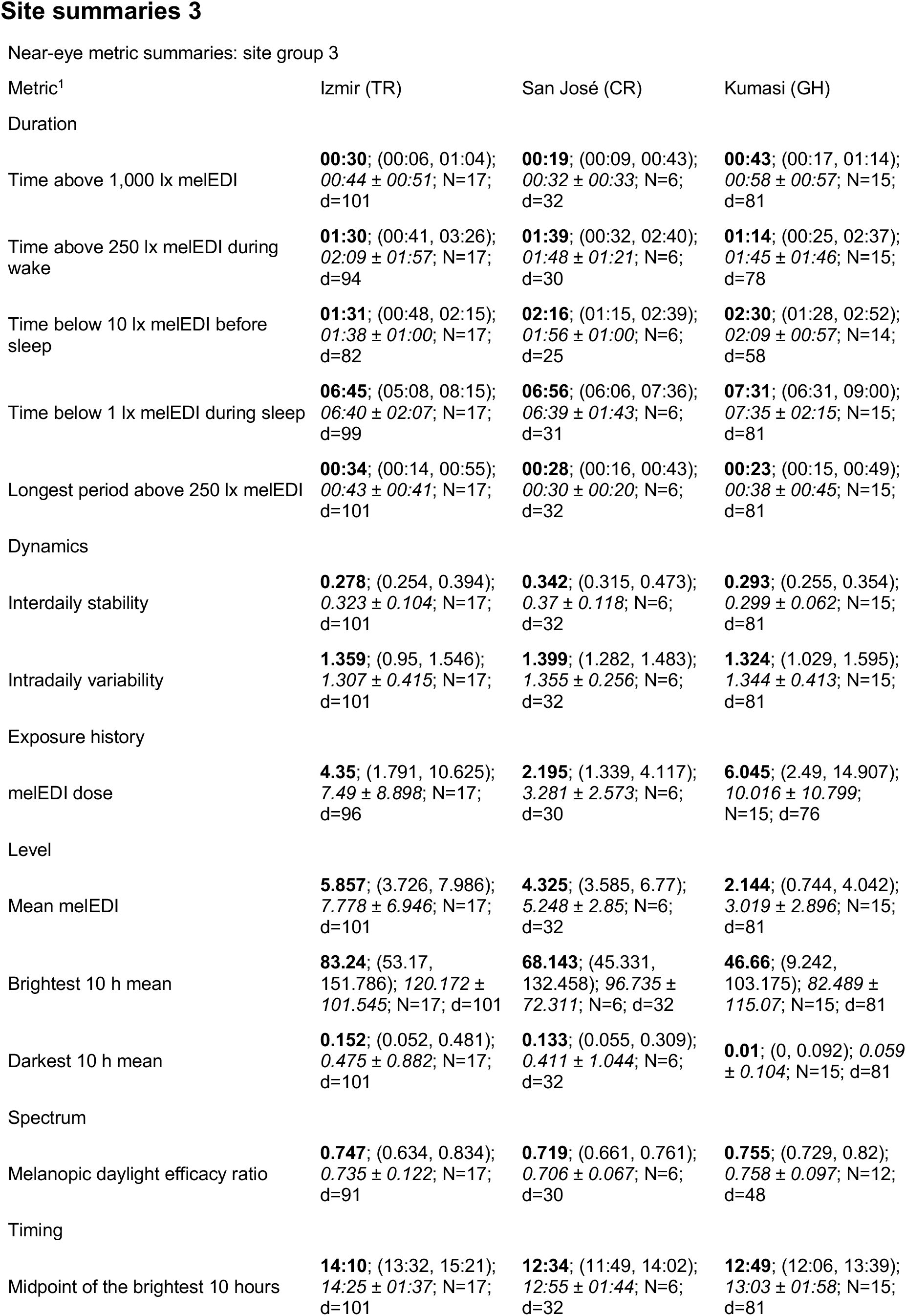
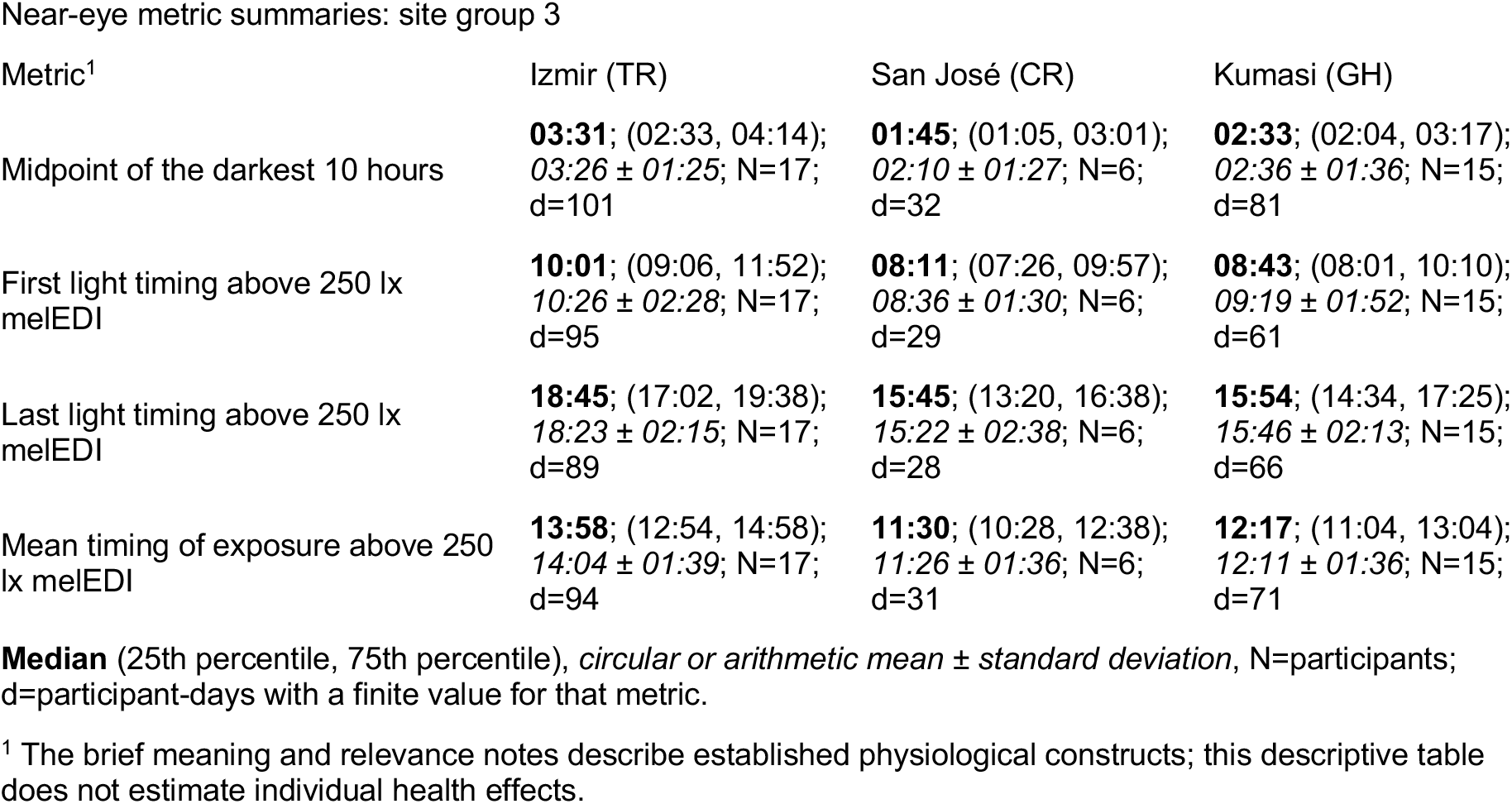
Definitions, exact support, median (interquartile range) summaries, and density displays for the 17 near-eye personal light-exposure metrics overall and by site. Values use three decimal places where applicable. Metrics are distinct summaries and should not be interpreted as interchangeable outcomes. Overall distribution

| Metric<sup>1</sup> | Unit | Overall | Scaling<sup>2</sup> | Distribution<sup>3</sup> |
| --- | --- | --- | --- | --- |
| Duration |  |  |  |  |
| Time above 1,000 lx melEDI; Bright-light exposure duration; relevant to daytime alerting and circadian entrainment. | HH:MM | **00:41**; (00:11, 01:34);<br>01:10 ± 01:28; N=141;<br>d=816 | Symlog<br>(base 10;<br>threshold 1) |  |
| Time above 250 lx melEDI during wake; Waking time in recommended daytime light; relevant to alertness, entrainment, and subsequent sleep. | HH:MM | **02:25**; (00:51, 04:37);<br>03:00 ± 02:29; N=141;<br>d=737 | Symlog<br>(base 10;<br>threshold 1) |  |
| Time below 10 lx melEDI before sleep; Low-light time before bed; limits evening melatonin suppression and circadian delay. | HH:MM | **01:53**; (01:02, 02:35);<br>01:51 ± 01:02; N=139;<br>d=655 | Symlog<br>(base 10;<br>threshold 1) |  |
| Time below 1 lx melEDI during sleep; Darkness during sleep; supports nocturnal melatonin and an undisturbed sleep environment. | HH:MM | **07:08**; (05:58, 08:15);<br>07:04 ± 01:57; N=141;<br>d=778 | Symlog<br>(base 10;<br>threshold 1) |  |
| Longest period above 250 lx melEDI; Longest sustained bright-light bout; captures continuity of daytime circadian stimulation. | HH:MM | **00:38**; (00:17, 01:12);<br>00:55 ± 01:00; N=141;<br>d=816 | Symlog<br>(base 10;<br>threshold 1) |  |
| Dynamics |  |  |  |  |
| Interdaily stability; Day-to-day regularity of the light–dark pattern; higher regularity supports circadian stability. | dimensionless | **0.308**; (0.248, 0.38);<br>0.318 ± 0.095; N=141;<br>d=816 | Identity |  |
| Intradaily variability; Within-day fragmentation of light exposure; higher values indicate less consolidated light–dark input. | dimensionless | **1.253**; (0.93, 1.502);<br>1.229 ± 0.389; N=141;<br>d=816 | Identity |  |
| Exposure history |  |  |  |  |
| melEDI dose; Intensity–duration-weighted melanopic exposure; summarizes cumulative non-visual retinal light input. | klx·h | **4.96**; (1.936, 12.313);<br>10.358 ± 16.216;<br>N=141; d=761 | Symlog<br>(base 10;<br>threshold 1) |  |

Near-eye metric summaries: overall distribution
| Metric<sup>1</sup> | Unit | Overall | Scaling<sup>2</sup> | Distribution<sup>3</sup> |
| --- | --- | --- | --- | --- |
| Level |  |  |  |  |
| Mean melEDI; Geometric average of daily melEDI values, including zeros; summarizes overall exposure while reducing peak influence. | lx | **5.154**; (2.831, 9.225); $7.587 \pm 9.496$; N=141; d=816 | Symlog (base 10; threshold 1) | |
| Brightest 10 h mean; Mean of the brightest 10 hours; reflects the strength of the main daytime light episode. | lx | **110.566**; (41.513, 243.212); $214.866 \pm 508.56$; N=141; d=816 | Symlog (base 10; threshold 1) | |
| Darkest 10 h mean; Mean of the darkest 10 hours; lower values during the biological night favour melatonin preservation and sleep. | lx | **0.103**; (0.02, 0.253); $0.243 \pm 0.513$; N=141; d=816 | Symlog (base 10; threshold 1) | |
| Spectrum |  |  |  |  |
| Melanopic daylight efficacy ratio; Mean of viable one-minute melEDI/illuminance ratios; indicates melanopic efficacy relative to visual light. | dimensionless | **0.724**; (0.643, 0.795); $0.724 \pm 0.116$; N=137; d=687 | Identity | |
| Timing |  |  |  |  |
| Midpoint of the brightest 10 hours; Centre time of the brightest 10 hours; indexes the main daily circadian light cue. | HH:MM; clock time | **13:44**; (12:48, 15:00); $13:56 \pm 01:51$; N=141; d=816 | Circular clock | |
| Midpoint of the darkest 10 hours; Centre time of the darkest 10 hours; indexes the main daily darkness cue. | HH:MM; clock time | **02:54**; (02:01, 03:50); $02:58 \pm 01:41$; N=141; d=816 | Circular clock | |
| First light timing above 250 lx melEDI; First waking bright-light exposure; morning timing can advance circadian phase and promote alertness. | HH:MM; clock time | **09:08**; (08:02, 10:40); $09:26 \pm 02:22$; N=140; d=727 | Circular clock | |
| Last light timing above 250 lx melEDI; Last bright-light exposure; later timing may delay circadian phase and sleep onset. | HH:MM; clock time | **18:08**; (16:27, 19:42); $18:03 \pm 02:36$; N=141; d=687 | Circular clock | |

Near-eye metric summaries: overall distribution
| Metric<sup>1</sup> | Unit | Overall | Scaling<sup>2</sup> | Distribution<sup>3</sup> |
| --- | --- | --- | --- | --- |
| Mean timing of exposure above 250 lx melEDI; Average bright-light timing; summarizes the phase of daily circadian stimulation. | HH:MM; clock time | **13:29**; (12:29, 14:39); **13:31 ± 01:52**; N=141; d=742 | Circular clock |  |

Near-eye metric summaries: site group 1
| Metric<sup>1</sup> | Borås (SE) | Delft (NL) | Dortmund (DE) |
| --- | --- | --- | --- |
| **Duration** |  |  |  |
| Time above 1,000 lx melEDI | **01:18**; (00:31, 03:24); **02:00 ± 01:54**; N=13; d=78 | **01:18**; (00:29, 02:12); **01:33 ± 01:28**; N=13; d=78 | **01:03**; (00:20, 02:32); **01:48 ± 02:14**; N=18; d=107 |
| Time above 250 lx melEDI during wake | **04:06**; (02:30, 06:15); **04:21 ± 02:24**; N=13; d=73 | **03:29**; (02:03, 05:16); **03:39 ± 02:25**; N=13; d=64 | **03:42**; (02:05, 05:58); **04:05 ± 02:51**; N=18; d=90 |
| Time below 10 lx melEDI before sleep | **02:16**; (01:30, 02:39); **02:05 ± 00:53**; N=13; d=67 | **01:50**; (01:21, 02:52); **01:59 ± 00:57**; N=12; d=50 | **01:42**; (00:53, 02:17); **01:41 ± 00:58**; N=18; d=84 |
| Time below 1 lx melEDI during sleep | **07:08**; (06:03, 08:18); **07:12 ± 01:46**; N=13; d=76 | **07:14**; (05:56, 08:06); **07:01 ± 01:50**; N=13; d=68 | **07:08**; (05:56, 08:16); **07:05 ± 01:50**; N=18; d=95 |
| Longest period above 250 lx melEDI | **00:56**; (00:28, 01:27); **01:12 ± 01:02**; N=13; d=78 | **00:48**; (00:23, 01:20); **01:09 ± 01:15**; N=13; d=78 | **01:00**; (00:21, 01:40); **01:20 ± 01:31**; N=18; d=107 |
| **Dynamics** |  |  |  |
| Interdaily stability | **0.311**; (0.272, 0.367); **0.315 ± 0.078**; N=13; d=78 | **0.258**; (0.239, 0.389); **0.288 ± 0.094**; N=13; d=78 | **0.264**; (0.219, 0.319); **0.274 ± 0.074**; N=18; d=107 |
| Intradaily variability | **0.997**; (0.818, 1.388); **1.088 ± 0.343**; N=13; d=78 | **1.193**; (0.96, 1.389); **1.238 ± 0.345**; N=13; d=78 | **1.182**; (0.928, 1.49); **1.177 ± 0.407**; N=18; d=107 |
| **Exposure history** |  |  |  |
| melEDI dose | **9.296**; (3.892, 21.895); **19.923 ± 27.784**; N=13; d=71 | **10.495**; (3.855, 17.395); **15.881 ± 20.395**; N=13; d=74 | **6.751**; (2.469, 14.435); **13.809 ± 23.711**; N=18; d=97 |
| **Level** |  |  |  |
| Mean melEDI | **7.569**; (4.55, 11.613); **9.171 ± 7.111**; N=13; d=78 | **6.659**; (4.18, 11.674); **8.274 ± 6.558**; N=13; d=78 | **7.704**; (3.081, 13.924); **12.017 ± 18.865**; N=18; d=107 |

Near-eye metric summaries: site group 1
| Metric<sup>1</sup> | Borås (SE) | Delft (NL) | Dortmund (DE) |
| --- | --- | --- | --- |
| Brightest 10 h mean | **252.689**; (104.514, 461.953); 399.302 ± 518.102; N=13; d=78 | **175.974**; (78.625, 299.106); 262.206 ± 329.201; N=13; d=78 | **156.184**; (67.783, 339.731); 422.54 ± 1,209.616; N=18; d=107 |
| Darkest 10 h mean | **0.08**; (0.016, 0.191); 0.183 ± 0.299; N=13; d=78 | **0.1**; (0.035, 0.23); 0.2 ± 0.332; N=13; d=78 | **0.14**; (0.036, 0.282); 0.223 ± 0.299; N=18; d=107 |
| Spectrum |  |  |  |
| Melanopic daylight efficacy ratio | **0.788**; (0.725, 0.872); 0.796 ± 0.162; N=13; d=74 | **0.743**; (0.699, 0.796); 0.743 ± 0.077; N=13; d=65 | **0.781**; (0.675, 0.853); 0.768 ± 0.118; N=18; d=97 |
| Timing |  |  |  |
| Midpoint of the brightest 10 hours | **13:16**; (12:28, 14:10); 13:20 ± 01:33; N=13; d=78 | **14:12**; (13:23, 15:18); 14:22 ± 01:51; N=13; d=78 | **13:52**; (12:54, 15:04); 14:00 ± 01:42; N=18; d=107 |
| Midpoint of the darkest 10 hours | **02:02**; (01:24, 02:48); 02:03 ± 01:07; N=13; d=78 | **02:55**; (02:01, 03:28); 02:50 ± 02:05; N=13; d=78 | **02:32**; (01:45, 03:22); 02:37 ± 01:19; N=18; d=107 |
| First light timing above 250 lx melEDI | **07:57**; (06:56, 08:33); 08:03 ± 01:32; N=13; d=74 | **09:04**; (08:18, 09:52); 09:17 ± 01:56; N=13; d=68 | **08:36**; (07:10, 10:13); 08:45 ± 02:21; N=18; d=93 |
| Last light timing above 250 lx melEDI | **18:41**; (17:38, 19:38); 18:39 ± 01:38; N=13; d=73 | **18:30**; (17:37, 19:59); 18:37 ± 02:08; N=13; d=62 | **19:34**; (18:04, 20:42); 19:16 ± 02:28; N=18; d=89 |
| Mean timing of exposure above 250 lx melEDI | **13:20**; (12:29, 14:24); 13:21 ± 01:25; N=13; d=72 | **13:52**; (13:13, 15:06); 14:02 ± 01:41; N=13; d=71 | **13:44**; (12:28, 14:55); 13:51 ± 01:56; N=18; d=93 |

Near-eye metric summaries: site group 2
| Metric<sup>1</sup> | Tübingen (DE) | Munich (DE) | Madrid (ES) |
| --- | --- | --- | --- |
| Duration |  |  |  |
| Time above 1,000 lx melEDI | **00:40**; (00:07, 01:25); 01:06 ± 01:22; N=26; d=150 | **00:39**; (00:10, 01:46); 01:09 ± 01:18; N=10; d=60 | **00:27**; (00:06, 00:53); 00:38 ± 00:47; N=23; d=129 |
| Time above 250 lx melEDI during wake | **02:01**; (00:47, 04:20); 02:52 ± 02:39; N=26; d=141 | **02:38**; (00:55, 04:19); 02:57 ± 02:19; N=10; d=54 | **02:43**; (00:54, 04:43); 03:02 ± 02:19; N=23; d=113 |
| Time below 10 lx melEDI before sleep | **01:52**; (01:04, 02:25); 01:52 ± 01:06; N=26; d=132 | **01:43**; (00:42, 02:32); 01:45 ± 01:10; N=10; d=52 | **01:43**; (00:58, 02:50); 01:50 ± 01:05; N=23; d=105 |
| Time below 1 lx melEDI during sleep | **06:54**; (05:52, 07:46); 06:47 ± 01:30; N=26; d=150 | **05:52**; (04:41, 07:03); 05:58 ± 01:58; N=10; d=60 | **08:06**; (07:10, 09:10); 08:02 ± 01:57; N=23; d=118 |

Near-eye metric summaries: site group 2
| Metric<sup>1</sup> | Tübingen (DE) | Munich (DE) | Madrid (ES) |
| --- | --- | --- | --- |
| Longest period above 250 lx melEDI | **00:35**; (00:13, 01:06);<br>00:50 ± 00:54; N=26;<br>d=150 | **00:38**; (00:18, 01:32);<br>01:01 ± 01:02; N=10;<br>d=60 | **00:38**; (00:16, 01:04);<br>00:45 ± 00:36; N=23;<br>d=129 |
| Dynamics |  |  |  |
| Interdaily stability | **0.329**; (0.263, 0.424);<br>0.344 ± 0.107; N=26;<br>d=150 | **0.244**; (0.218, 0.277);<br>0.252 ± 0.058; N=10;<br>d=60 | **0.352**; (0.321, 0.455);<br>0.368 ± 0.092; N=23;<br>d=129 |
| Intradaily variability | **1.154**; (0.801, 1.442);<br>1.125 ± 0.354; N=26;<br>d=150 | **1.48**; (0.869, 1.642);<br>1.332 ± 0.477; N=10;<br>d=60 | **1.251**; (0.999, 1.497);<br>1.251 ± 0.413; N=23;<br>d=129 |
| Exposure history |  |  |  |
| melEDI dose | **3.862**; (1.411, 10.217);<br>7.719 ± 9.847; N=26;<br>d=137 | **7.11**; (1.897, 15.789);<br>10.055 ± 11.237;<br>N=10; d=55 | **3.317**; (1.703, 6.29);<br>6.115 ± 8.79; N=23;<br>d=125 |
| Level |  |  |  |
| Mean melEDI | **5.232**; (3.224, 8.188);<br>7.221 ± 6.33; N=26;<br>d=150 | **9.013**; (4.011, 15.089);<br>12.102 ± 12.013;<br>N=10; d=60 | **3.505**; (1.469, 6.211);<br>4.166 ± 3.272; N=23;<br>d=129 |
| Brightest 10 h mean | **100.057**; (33.255,<br>209.904); 179.372 ±<br>241.53; N=26; d=150 | **130.78**; (54.044,<br>259.726); 201.947 ±<br>194.042; N=10; d=60 | **101.102**; (24.355,<br>218.457); 136.312 ±<br>134.07; N=23; d=129 |
| Darkest 10 h mean | **0.149**; (0.074, 0.309);<br>0.273 ± 0.465; N=26;<br>d=150 | **0.325**; (0.119, 0.71);<br>0.515 ± 0.653; N=10;<br>d=60 | **0.022**; (0, 0.08); 0.054<br>± 0.075; N=23; d=129 |
| Spectrum |  |  |  |
| Melanopic daylight efficacy ratio | **0.65**; (0.592, 0.728);<br>0.667 ± 0.098; N=26;<br>d=144 | **0.724**; (0.652, 0.773);<br>0.72 ± 0.093; N=10;<br>d=55 | **0.668**; (0.629, 0.721);<br>0.673 ± 0.07; N=22;<br>d=83 |
| Timing |  |  |  |
| Midpoint of the brightest 10 hours | **13:38**; (12:47, 14:44);<br>13:52 ± 01:42; N=26;<br>d=150 | **14:54**; (12:50, 16:03);<br>14:31 ± 02:14; N=10;<br>d=60 | **13:44**; (13:06, 15:20);<br>14:10 ± 01:51; N=23;<br>d=129 |
| Midpoint of the darkest 10 hours | **03:08**; (02:11, 04:10);<br>03:16 ± 01:34; N=26;<br>d=150 | **02:37**; (01:54, 04:17);<br>03:10 ± 02:00; N=10;<br>d=60 | **03:39**; (02:43, 04:34);<br>03:34 ± 01:49; N=23;<br>d=129 |
| First light timing above 250 lx<br>melEDI | **09:43**; (08:13, 10:53);<br>09:45 ± 02:21; N=25;<br>d=133 | **09:36**; (07:58, 10:53);<br>09:39 ± 03:12; N=10;<br>d=56 | **09:32**; (08:40, 11:21);<br>10:05 ± 02:18; N=23;<br>d=118 |
| Last light timing above 250 lx<br>melEDI | **17:37**; (15:35, 19:09);<br>17:18 ± 02:44; N=26;<br>d=121 | **19:50**; (18:37, 20:50);<br>19:33 ± 02:11; N=10;<br>d=53 | **17:48**; (16:46, 19:19);<br>18:01 ± 02:21; N=23;<br>d=106 |
| Mean timing of exposure above 250<br>lx melEDI | **13:17**; (12:17, 14:31);<br>13:24 ± 01:45; N=26;<br>d=138 | **14:04**; (12:58, 15:25);<br>14:09 ± 02:22; N=10;<br>d=55 | **13:43**; (12:55, 14:29);<br>13:47 ± 01:33; N=23;<br>d=117 |

| Metric<sup>1</sup> | Izmir (TR) | San José (CR) | Kumasi (GH) |
| --- | --- | --- | --- |
| Duration |  |  |  |
| Time above 1,000 lx melEDI | **00:30**; (00:06, 01:04);<br>00:44 ± 00:51; N=17;<br>d=101 | **00:19**; (00:09, 00:43);<br>00:32 ± 00:33; N=6;<br>d=32 | **00:43**; (00:17, 01:14);<br>00:58 ± 00:57; N=15;<br>d=81 |
| Time above 250 lx melEDI during wake | **01:30**; (00:41, 03:26);<br>02:09 ± 01:57; N=17;<br>d=94 | **01:39**; (00:32, 02:40);<br>01:48 ± 01:21; N=6;<br>d=30 | **01:14**; (00:25, 02:37);<br>01:45 ± 01:46; N=15;<br>d=78 |
| Time below 10 lx melEDI before sleep | **01:31**; (00:48, 02:15);<br>01:38 ± 01:00; N=17;<br>d=82 | **02:16**; (01:15, 02:39);<br>01:56 ± 01:00; N=6;<br>d=25 | **02:30**; (01:28, 02:52);<br>02:09 ± 00:57; N=14;<br>d=58 |
| Time below 1 lx melEDI during sleep | **06:45**; (05:08, 08:15);<br>06:40 ± 02:07; N=17;<br>d=99 | **06:56**; (06:06, 07:36);<br>06:39 ± 01:43; N=6;<br>d=31 | **07:31**; (06:31, 09:00);<br>07:35 ± 02:15; N=15;<br>d=81 |
| Longest period above 250 lx melEDI | **00:34**; (00:14, 00:55);<br>00:43 ± 00:41; N=17;<br>d=101 | **00:28**; (00:16, 00:43);<br>00:30 ± 00:20; N=6;<br>d=32 | **00:23**; (00:15, 00:49);<br>00:38 ± 00:45; N=15;<br>d=81 |
| Dynamics |  |  |  |
| Interdaily stability | **0.278**; (0.254, 0.394);<br>0.323 ± 0.104; N=17;<br>d=101 | **0.342**; (0.315, 0.473);<br>0.37 ± 0.118; N=6;<br>d=32 | **0.293**; (0.255, 0.354);<br>0.299 ± 0.062; N=15;<br>d=81 |
| Intradaily variability | **1.359**; (0.95, 1.546);<br>1.307 ± 0.415; N=17;<br>d=101 | **1.399**; (1.282, 1.483);<br>1.355 ± 0.256; N=6;<br>d=32 | **1.324**; (1.029, 1.595);<br>1.344 ± 0.413; N=15;<br>d=81 |
| Exposure history |  |  |  |
| melEDI dose | **4.35**; (1.791, 10.625);<br>7.49 ± 8.898; N=17;<br>d=96 | **2.195**; (1.339, 4.117);<br>3.281 ± 2.573; N=6;<br>d=30 | **6.045**; (2.49, 14.907);<br>10.016 ± 10.799;<br>N=15; d=76 |
| Level |  |  |  |
| Mean melEDI | **5.857**; (3.726, 7.986);<br>7.778 ± 6.946; N=17;<br>d=101 | **4.325**; (3.585, 6.77);<br>5.248 ± 2.85; N=6;<br>d=32 | **2.144**; (0.744, 4.042);<br>3.019 ± 2.896; N=15;<br>d=81 |
| Brightest 10 h mean | **83.24**; (53.17,<br>151.786); 120.172 ±<br>101.545; N=17; d=101 | **68.143**; (45.331,<br>132.458); 96.735 ±<br>72.311; N=6; d=32 | **46.66**; (9.242,<br>103.175); 82.489 ±<br>115.07; N=15; d=81 |
| Darkest 10 h mean | **0.152**; (0.052, 0.481);<br>0.475 ± 0.882; N=17;<br>d=101 | **0.133**; (0.055, 0.309);<br>0.411 ± 1.044; N=6;<br>d=32 | **0.01**; (0, 0.092); 0.059<br>± 0.104; N=15; d=81 |
| Spectrum |  |  |  |
| Melanopic daylight efficacy ratio | **0.747**; (0.634, 0.834);<br>0.735 ± 0.122; N=17;<br>d=91 | **0.719**; (0.661, 0.761);<br>0.706 ± 0.067; N=6;<br>d=30 | **0.755**; (0.729, 0.82);<br>0.758 ± 0.097; N=12;<br>d=48 |
| Timing |  |  |  |
| Midpoint of the brightest 10 hours | **14:10**; (13:32, 15:21);<br>14:25 ± 01:37; N=17;<br>d=101 | **12:34**; (11:49, 14:02);<br>12:55 ± 01:44; N=6;<br>d=32 | **12:49**; (12:06, 13:39);<br>13:03 ± 01:58; N=15;<br>d=81 |

Near-eye metric summaries: site group 3
| Metric<sup>1</sup> | Izmir (TR) | San José (CR) | Kumasi (GH) |
| --- | --- | --- | --- |
| Midpoint of the darkest 10 hours | **03:31**; (02:33, 04:14);<br>03:26 ± 01:25; N=17;<br>d=101 | **01:45**; (01:05, 03:01);<br>02:10 ± 01:27; N=6;<br>d=32 | **02:33**; (02:04, 03:17);<br>02:36 ± 01:36; N=15;<br>d=81 |
| First light timing above 250 lx<br>melEDI | **10:01**; (09:06, 11:52);<br>10:26 ± 02:28; N=17;<br>d=95 | **08:11**; (07:26, 09:57);<br>08:36 ± 01:30; N=6;<br>d=29 | **08:43**; (08:01, 10:10);<br>09:19 ± 01:52; N=15;<br>d=61 |
| Last light timing above 250 lx<br>melEDI | **18:45**; (17:02, 19:38);<br>18:23 ± 02:15; N=17;<br>d=89 | **15:45**; (13:20, 16:38);<br>15:22 ± 02:38; N=6;<br>d=28 | **15:54**; (14:34, 17:25);<br>15:46 ± 02:13; N=15;<br>d=66 |
| Mean timing of exposure above 250<br>lx melEDI | **13:58**; (12:54, 14:58);<br>14:04 ± 01:39; N=17;<br>d=94 | **11:30**; (10:28, 12:38);<br>11:26 ± 01:36; N=6;<br>d=31 | **12:17**; (11:04, 13:04);<br>12:11 ± 01:36; N=15;<br>d=71 |

**Median** (25th percentile, 75th percentile), *circular or arithmetic mean ± standard deviation*, N=participants; d=participant-days with a finite value for that metric.
<sup>1</sup> The brief meaning and relevance notes describe established physiological constructs; this descriptive table does not estimate individual health effects.

**Supplementary Table S3.**
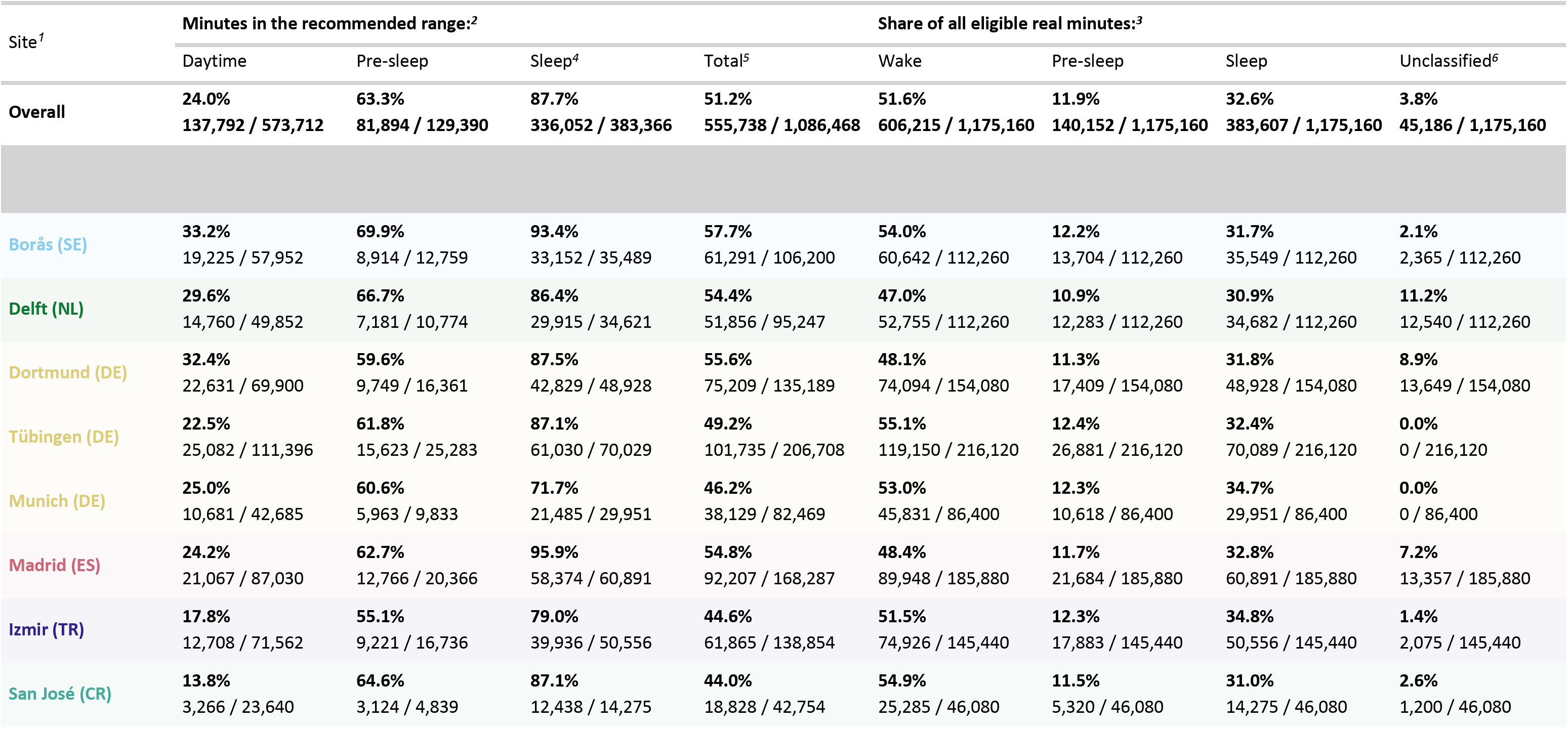
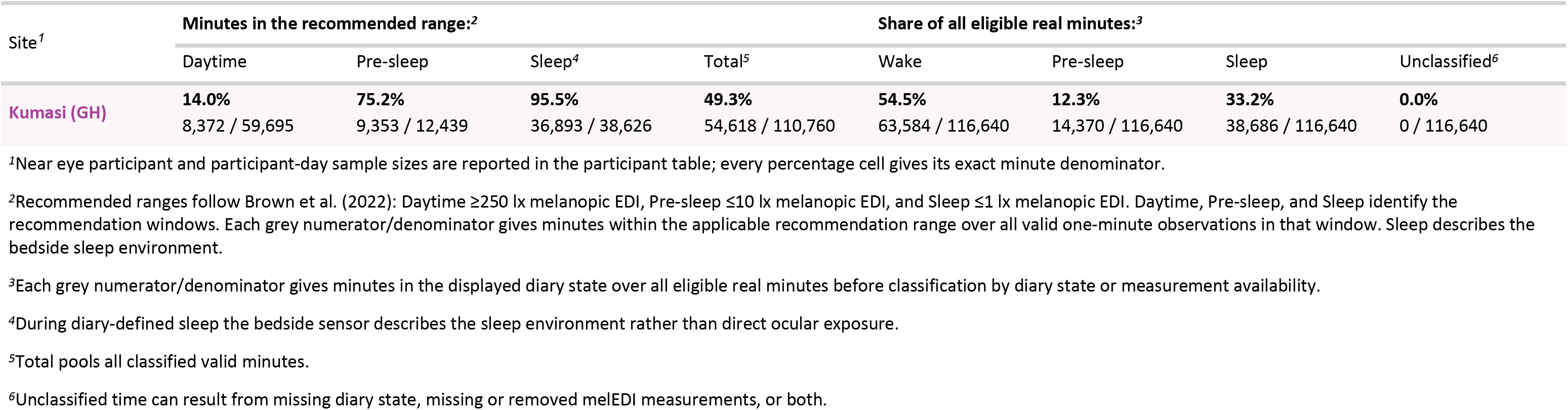
Fractions of valid near-eye minutes meeting each Brown et al. recommendation range, with explicit numerators and denominators. Daytime, Pre-sleep, and Sleep identify the recommendation windows. The Sleep summaries use near-eye illuminance recorded during sleep and describe the bedside sleep environment rather than verified ocular exposure. These pooled-minute summaries are descriptive and differ from the modelled recommendation-window-period estimates in main Table 2.

**Supplementary Table S4.** Exploratory within-participant and between-participant associations across Brown et al. recommendation windows. Results are percentage-point differences per 10 percentage points higher Daytime adherence. The four tests form one FDR family. The within-participant day-level claim is withheld because serial dependence remains unresolved. Between-participant results describe observed monitoring-period averages and do not establish rankings, stable traits, or causal effects.

| Window | Difference, percentage points (95% CI) | FDR-adjusted p |
| --- | --- | --- |
| Within participant |  |  |
| Sleep | -0.26 (-0.95 to 0.43) | 0.453 |
| Pre-sleep | -1.09 (-2.75 to 0.56) | 0.258 |
| Between participants |  |  |
| Sleep | -2.52 (-3.55 to -1.48) | <0.001 |
| Pre-sleep | -3.59 (-5.94 to -1.25) | 0.005 |

Contrasts are per 10 percentage points higher daytime adherence. Within-participant contrasts concern deviations from each participant's monitoring-period average; between-participant contrasts concern those averages. The four associations form one FDR family. Unresolved temporal dependence precludes a within-participant day-level claim.

**Supplementary Table S5.** Near-eye fitted-curve dispersion and Shapley allocation of full-model in-sample R². Fitted-curve variation is in squared log10(melanopic EDI + 0.1 lx) prediction units; dispersion and R²-share ratios are unitless. The allocation includes the shared local-clock curve in every component model as the baseline. Intervals are 95% hierarchical cluster-bootstrap percentile intervals from 2,000 replicates and are conditional on the fitted models. Dispersion and R² allocation describe different estimands and should not be added or interpreted causally.

|  | Fitted-curve result (95% CI) | Shapley allocation (95% CI) | Share of full-model R<sup>2</sup> (95% CI) |
| --- | --- | --- | --- |
| Shared local-clock curve | Not applicable | 0.611 (0.550 to 0.657) | 78.9 (73.6 to 82.6)% |
| Site pattern | 0.100 (0.041 to 0.139) | 0.016 (0.007 to 0.026) | 2.0 (0.9 to 3.5)% |
| Participant pattern | 0.180 (0.126 to 0.214) | 0.100 (0.079 to 0.125) | 12.9 (10.1 to 16.4)% |
| Participant-day shift | 0.019 (0.010 to 0.024) | 0.048 (0.035 to 0.064) | 6.2 (4.4 to 8.6)% |
| Participant pattern + day shift | 0.200 (0.140 to 0.230) | Not separately allocated | Not applicable |
| Participant / site | 1.797 (1.157 to 4.332) | 6.41 (3.85 to 15.14) | Not applicable |
| (Participant + day) / site | 1.991 (1.291 to 4.767) | 9.49 (5.77 to 21.81) | 90.5 (85.2 to 95.6)% |

Fitted-curve variation is in squared log<sub>10</sub>(melanopic EDI + 0.1 lx) prediction units; ratios are unitless. The shared local-clock curve remains the baseline in every component model. Intervals are conditional hierarchical cluster-bootstrap percentiles from 2,000 replicates. In the final row, the percentage is the participant-plus-day share of heterogeneity, excluding the shared local-clock contribution. Dispersion and R<sup>2</sup> allocation are distinct estimands.

**Supplementary Table S6.** Complementary chest fitted-curve dispersion and Shapley allocation of full-model in-sample R². The quantities, bootstrap intervals and interpretation follow Supplementary Table S5; chest and near-eye measurements remain distinct sensor-position estimands.

|  | Fitted-curve result (95% CI) | Shapley allocation (95% CI) | Share of full-model R<sup>2</sup> (95% CI) |
| --- | --- | --- | --- |
| Shared local-clock curve | Not applicable | 0.583 (0.523 to 0.629) | 79.1 (73.6 to 82.8)% |
| Site pattern | 0.101 (0.049 to 0.116) | 0.015 (0.008 to 0.024) | 2.1 (1.1 to 3.3)% |
| Participant pattern | 0.148 (0.100 to 0.199) | 0.087 (0.071 to 0.110) | 11.9 (9.5 to 15.3)% |
| Participant-day shift | 0.035 (0.020 to 0.039) | 0.051 (0.040 to 0.067) | 7.0 (5.3 to 9.4)% |
| Participant pattern + day shift | 0.183 (0.127 to 0.228) | Not separately allocated | Not applicable |
| Participant / site | 1.465 (0.973 to 3.179) | 5.69 (3.72 to 10.68) | Not applicable |
| (Participant + day) / site | 1.809 (1.219 to 3.736) | 9.02 (6.01 to 17.03) | 90.0 (85.7 to 94.5)% |

Fitted-curve variation is in squared log<sub>10</sub>(melanopic EDI + 0.1 lx) prediction units; ratios are unitless. The shared local-clock curve remains the baseline in every component model. Intervals are conditional hierarchical cluster-bootstrap percentiles from 2,000 replicates. In the final row, the percentage is the participant-plus-day share of heterogeneity, excluding the shared local-clock contribution. Dispersion and R<sup>2</sup> allocation are distinct estimands.

**Supplementary Table S7.**
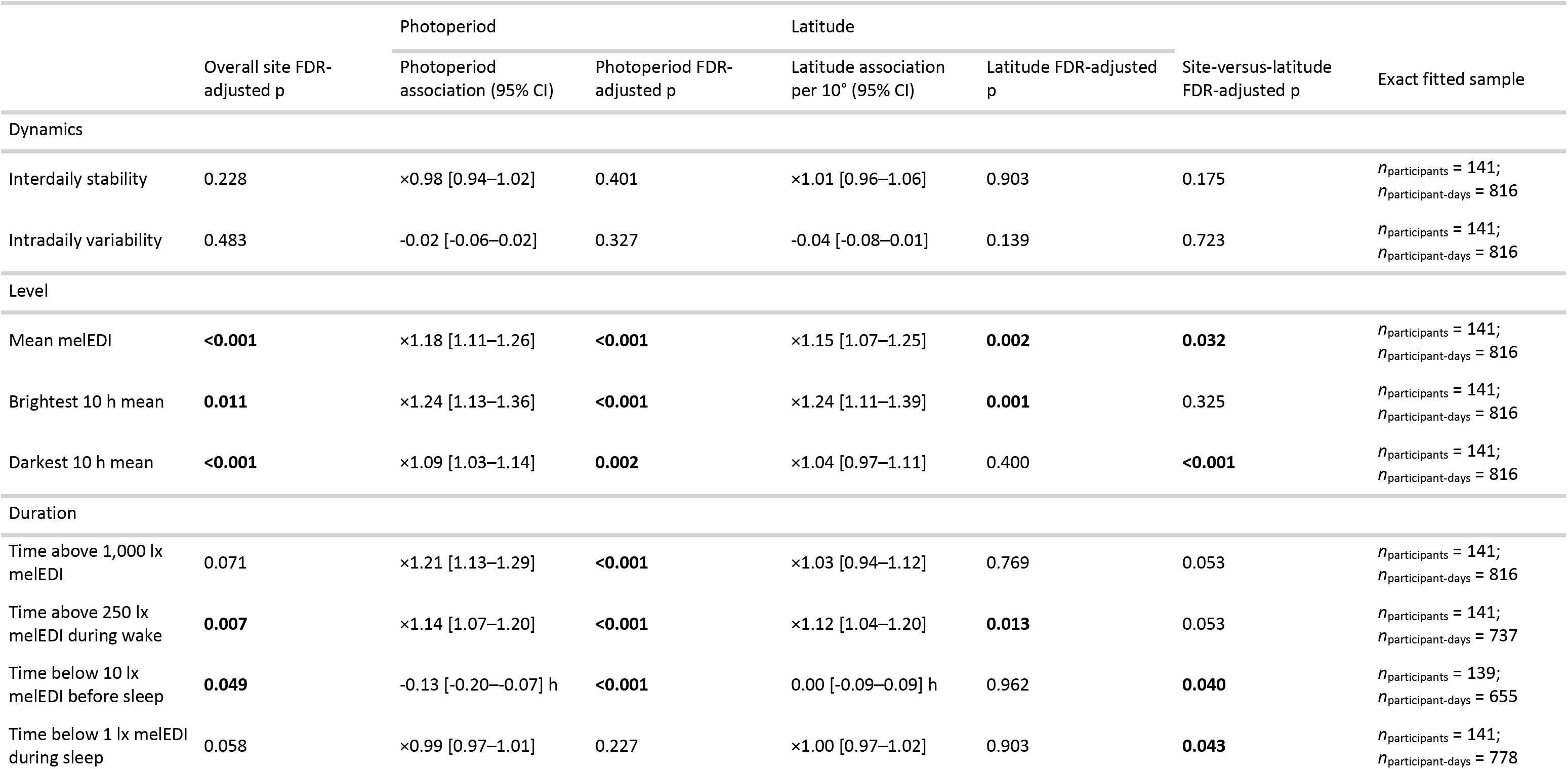
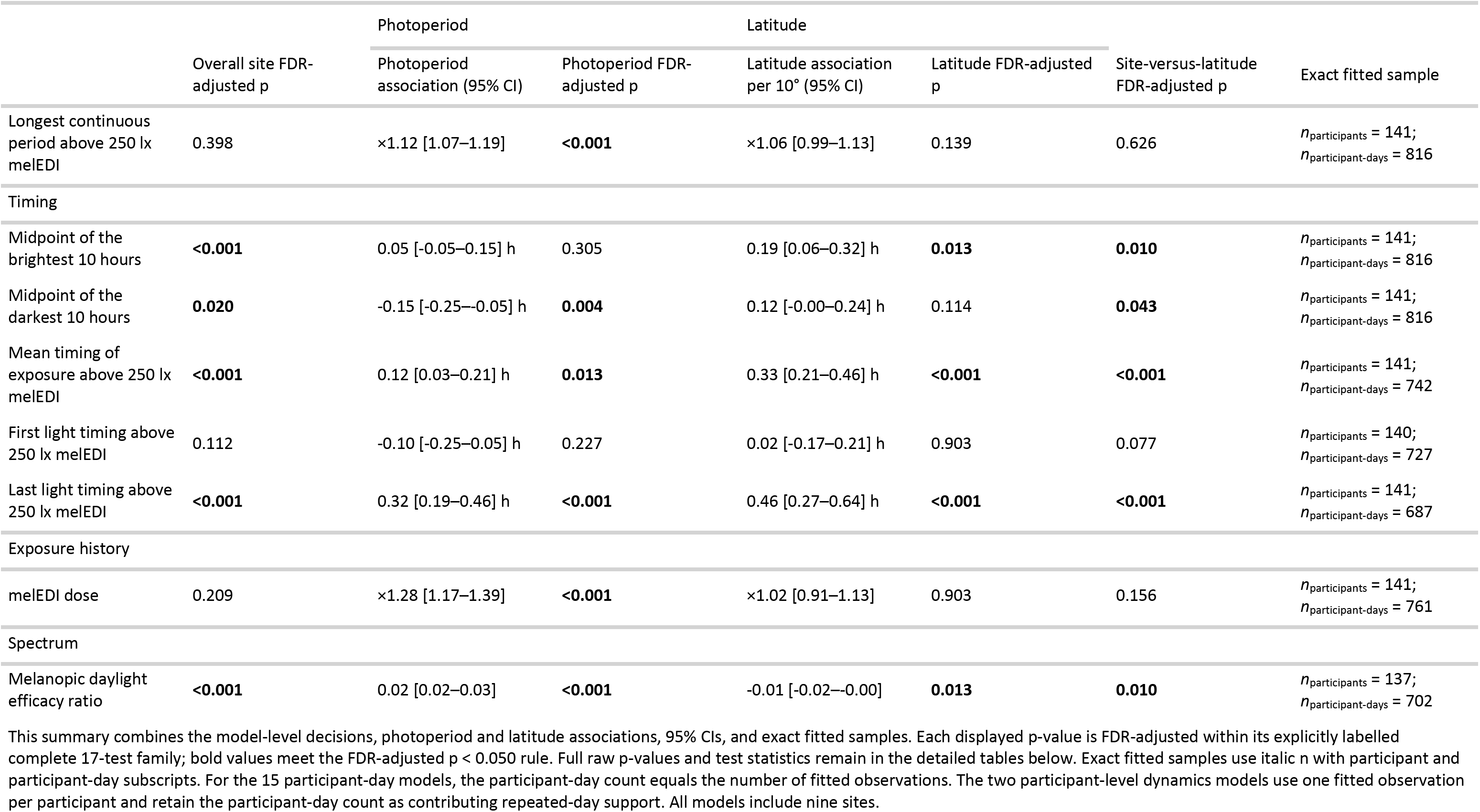
Overall-site, civil-photoperiod and absolute-latitude associations for the 17 near-eye personal light-exposure metrics. Each displayed *p* value is FDR-adjusted within its stated 17-test family. Site and latitude were evaluated in separate models because absolute latitude is fixed within and confounded with study site.

**Supplementary Table S8.** Derivative-based classifications for nine near-eye personal light-exposure metrics across the observed civil-photoperiod range. A qualifying transition required the preceding fitted-slope interval to be wholly above zero, the next interval to include zero and every later interval through the recorded maximum to remain zero-compatible. These classifications are descriptive and do not identify a physiological or environmental ceiling.

|  | Qualifying transition | Qualifying transition bracket (h) | Endpoint derivative | Interpretation |
| --- | --- | --- | --- | --- |
| Light level |  |  |  |  |
| Mean melEDI | Yes | 16.10–16.20 | 0.008 (95% CI -0.111 to 0.127) | Detected increase followed by a sustained zero-compatible tail |
| Brightest 10 h mean | Yes | 15.17–15.27 | 0.059 (95% CI -0.083 to 0.201) | Detected increase followed by a sustained zero-compatible tail |
| Darkest 10 h mean | Yes | 14.86–14.96 | -0.086 (95% CI -0.216 to 0.044) | Detected increase followed by a sustained zero-compatible tail |
| Duration and continuous period |  |  |  |  |
| Time above 1,000 lx melEDI | Yes | 14.86–14.96 | 0.098 (95% CI -0.176 to 0.371) | Detected increase followed by a sustained zero-compatible tail |
| Time above 250 lx melEDI during wake | Yes | 14.35–14.45 | 0.060 (95% CI -0.159 to 0.279) | Detected increase followed by a sustained zero-compatible tail |
| Time below 10 lx melEDI before sleep | No | Not applicable | -0.085 (95% CI -0.154 to -0.016) | No preceding detected increase |
| Time below 1 lx melEDI during sleep | No | Not applicable | 0.062 (95% CI -0.042 to 0.165) | No preceding detected increase |
| Longest continuous period above 250 lx melEDI | No | Not applicable | 0.041 (95% CI 0.025 to 0.058) | Increase remained detected at the recorded maximum |
| Exposure history |  |  |  |  |
| melEDI dose | Yes | 14.24–14.35 | 0.070 (95% CI -0.069 to 0.209) | Detected increase followed by a sustained zero-compatible tail |

A 'Yes' is the formal derivative-defined classification: the immediately preceding point has an interval wholly above zero, the next contains zero, and every later interval through the recorded maximum remains zero-compatible. Endpoint derivatives and 95% confidence intervals are on each model's linear-predictor scale, in model-scale units per hour of civil photoperiod.

**Supplementary Table S9.**
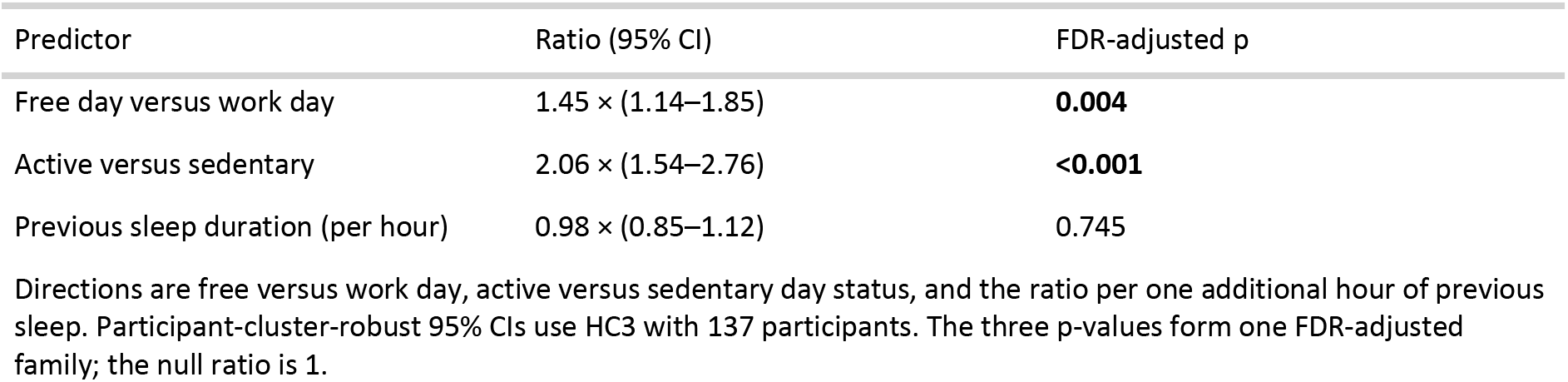
Common-association estimates from the participant-hour analysis of Free versus Work day, Active versus Sedentary daily activity status and each additional hour of previous-night sleep duration. Ratios and participant-cluster-robust 95% confidence intervals come from the fixed-site model; the three *p* values form one FDR-adjusted family. Predictor-by-site differences are shown in Supplementary Figure S13.

| Predictor | Ratio (95% CI) | FDR-adjusted p |
| --- | --- | --- |
| Free day versus work day | 1.45 × (1.14–1.85) | **0.004** |
| Active versus sedentary | 2.06 × (1.54–2.76) | **<0.001** |
| Previous sleep duration (per hour) | 0.98 × (0.85–1.12) | 0.745 |

Directions are free versus work day, active versus sedentary day status, and the ratio per one additional hour of previous sleep. Participant-cluster-robust 95% CIs use HC3 with 137 participants. The three p-values form one FDR-adjusted family; the null ratio is 1.

**Supplementary Table S10.** Summary of person-level results in the near-eye analyses. Each row identifies the analysis-specific FDR-correction set or decision structure applied to that result. Results that did not meet the applicable FDR criterion are inconclusive rather than evidence of no association. Four light-behaviour sleep-environment cells were unfit for inference. The two chronotype instruments were analysed separately. Biological sex and gender were recorded separately; gender was not analysed. The complete available-data curve difference was attenuated and not retained in the activity-complete near-eye analysis.

|  | Scale | Result | FDR family | Decision | Sample | Qualification |
| --- | --- | --- | --- | --- | --- | --- |
| Light-exposure behaviour and awareness | Metric associations per participant-level factor SD | None retained; 4 cells were unfit for inference | 68 factor-by-metric tests | None retained | Metric-specific samples in Supplementary Table S11 | Non-retention is inconclusive rather than evidence of no association. |
| Visual light sensitivity | Metric associations per VLSQ-8 SD | Melanopic EDI dose ratio: 0.846 (0.716 to 0.999) | 9 metric tests | None retained; dose adjusted p = 0.160 | 141 participants; 761 participant-days; 9 sites | Confidence intervals and FDR decisions are separate summaries. |
| Corrected midsleep on free days | Timing per one-hour later corrected midsleep | First light timing above 250 lx melEDI: 0.381 (0.146 to 0.616) h; Midpoint of the darkest 10 hours: 0.276 (0.118 to 0.435) h; Midpoint of the brightest 10 hours: 0.205 (0.045 to 0.366) h | 5 MCTQ timing outcomes | 3 associations retained | 139 to 140 participants; 722 to 810 participant-days; 9 sites | The two chronotype instruments are analysed separately; associations do not identify causal direction. |
| Morningness-eveningness preference | Timing per 10 points greater morning preference | First light timing above 250 lx melEDI: -0.444 (-0.699 to -0.189) h; Midpoint of the darkest 10 hours: -0.314 (-0.487 to -0.142) h; Midpoint of the brightest 10 hours: -0.278 (-0.451 to -0.105) h | 5 MEQ timing outcomes | 3 associations retained | 140 to 141 participants; 727 to 816 participant-days; 9 sites | The two chronotype instruments are analysed separately; associations do not identify causal direction. |
| Age | Metric associations per 10 years | Brightest 10 h mean: 1.31× (1.09–1.58); Time above 1,000 lx melEDI: 1.27× (1.11–1.45); melEDI dose: 1.31× (1.09–1.56) | 17 near-eye metric tests | 3 associations retained | 141 participants; 761 to 816 participant-days; 9 sites | Cross-sectional associations may reflect cohort, occupation, behaviour or other confounding. |
| Metric-level biological sex | Female minus Male | None retained | 17 near-eye metric tests | None retained | Metric-specific samples in Supplementary Table S14 | Biological sex and gender were recorded separately; gender was not analysed. |
| Biological-sex-specific daily curve | Complete curve and activity-complete sensitivity | Complete available-data curve: FDR-adjusted p = 0.028. Activity-complete adjusted curve: exploratory global curve not supported. | Global complete-curve tests and separate sensitivity decisions | Analysis-specific global decisions | 141 participants; 816 participant-days; 9 sites; 37,756 observations | Restriction and activity adjustment cannot be disentangled as mechanisms. Clock-specific intervals are pointwise. |

**Supplementary Table S11.**
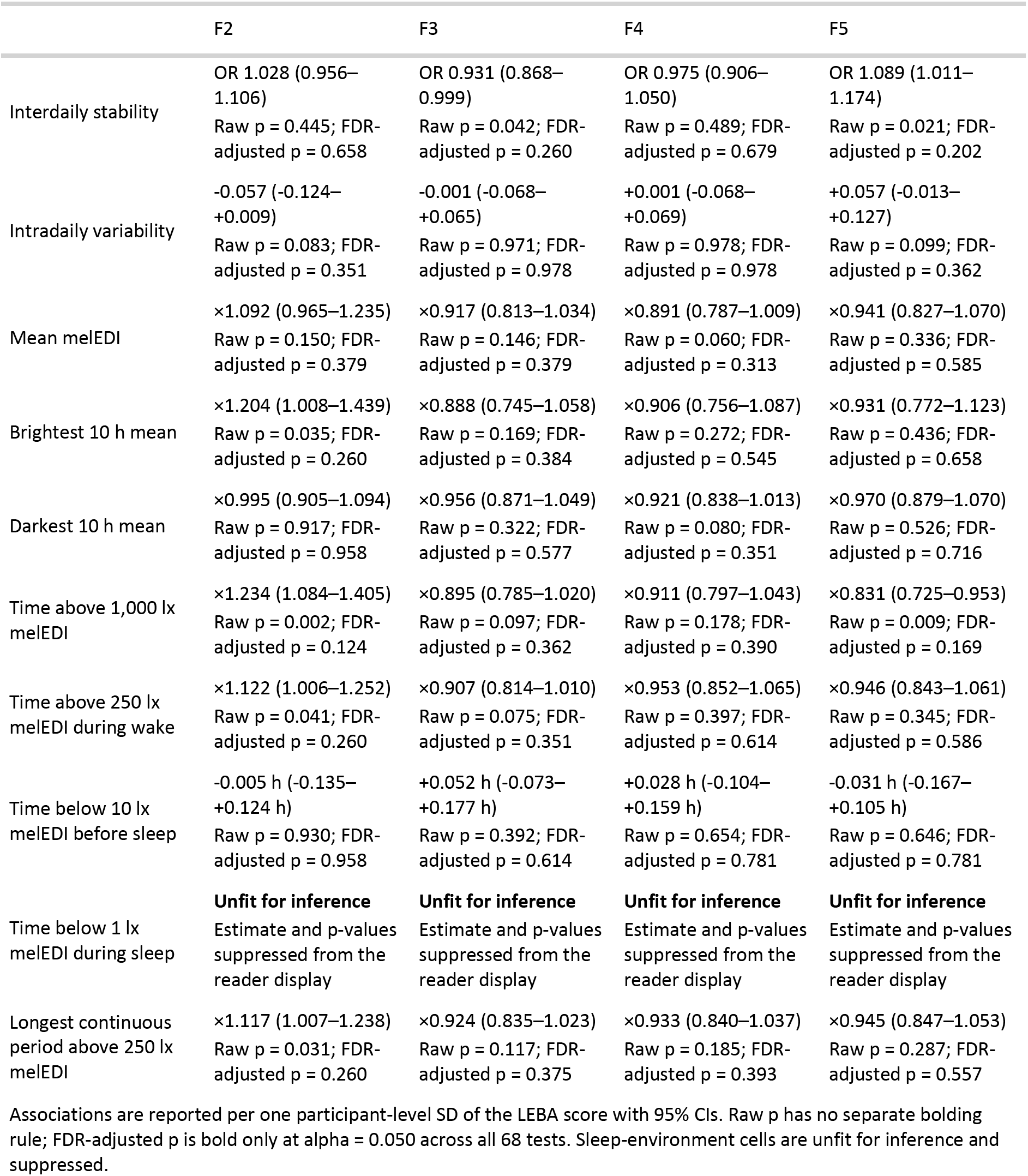
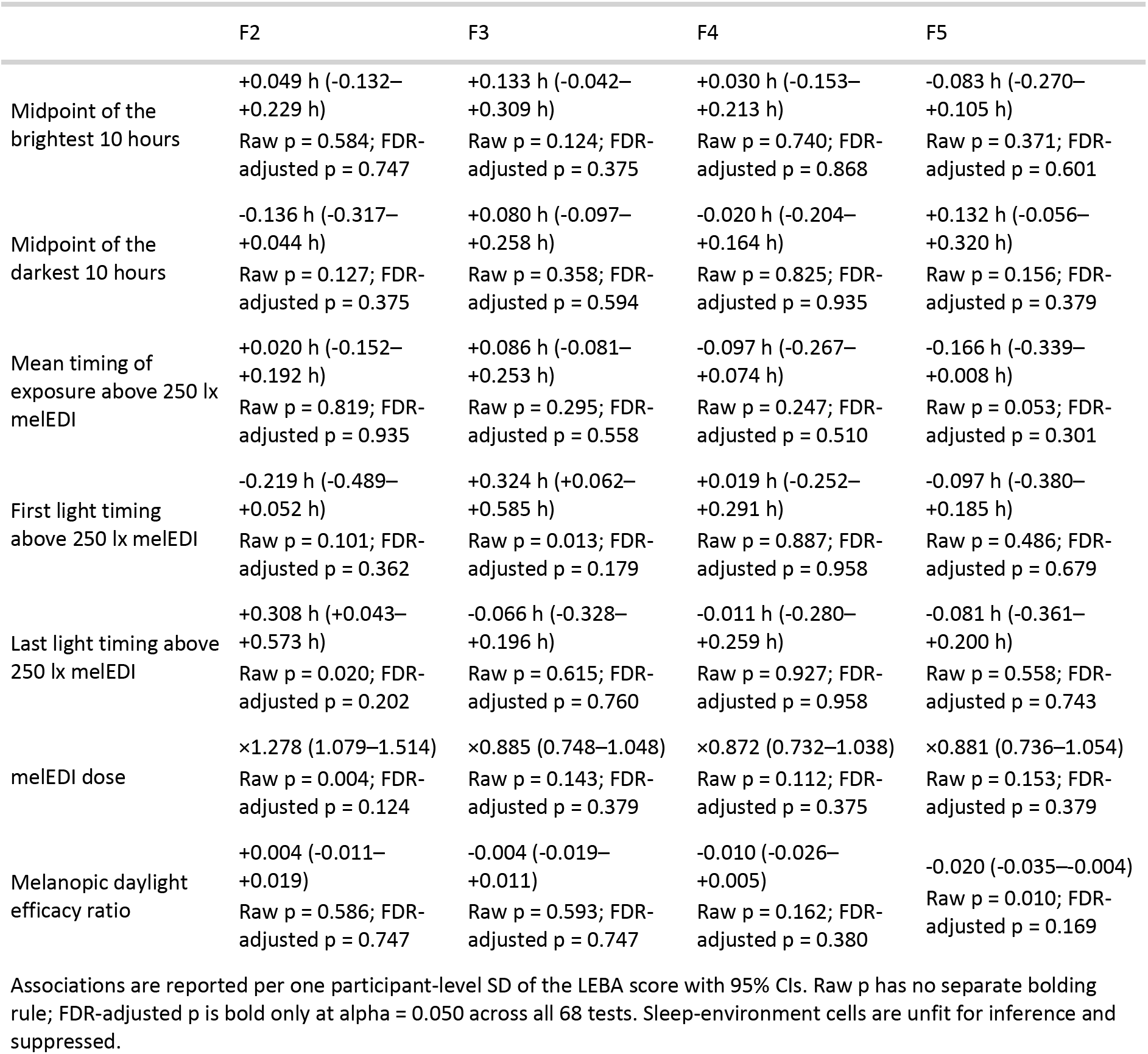
Light-exposure behaviour and awareness (part 1)

**Supplementary Table S12.** Visual light sensitivity.

| Metric | Scale | Association per VLSQ-8 SD (95% CI) | Raw p | FDR-adjusted p | Participants / participant-days / sites |
| --- | --- | --- | --- | --- | --- |
| Mean melEDI | Ratio (percentage change) | ×0.915 (0.811–1.031) | 0.147 | 0.221 | 141 / 816 / 9 |
| Brightest 10 h mean | Ratio (percentage change) | ×0.851 (0.716–1.013) | 0.071 | 0.160 | 141 / 816 / 9 |
| Darkest 10 h mean | Ratio (percentage change) | ×1.024 (0.934–1.123) | 0.612 | 0.689 | 141 / 816 / 9 |
| Time above 1,000 lx melEDI | Ratio (percentage change) | ×0.876 (0.765–1.002) | 0.056 | 0.160 | 141 / 816 / 9 |
| Time above 250 lx melEDI during wake | Ratio (percentage change) | ×0.900 (0.806–1.005) | 0.063 | 0.160 | 141 / 737 / 9 |
| Time below 10 lx melEDI before sleep | Difference in hours | +0.063 h (-0.062–+0.188 h) | 0.328 | 0.422 | 139 / 655 / 9 |
| Time below 1 lx melEDI during sleep | Ratio (percentage change) | ×0.998 (0.964–1.033) | 0.903 | 0.903 | 141 / 778 / 9 |
| Longest continuous period above 250 lx melEDI | Ratio (percentage change) | ×0.917 (0.828–1.014) | 0.093 | 0.167 | 141 / 816 / 9 |
| melEDI dose | Ratio (percentage change) | ×0.846 (0.716–0.999) | 0.051 | 0.160 | 141 / 761 / 9 |

Associations compare scores separated by one participant-level SD (5.540 VLSQ-8 points). The hour-scale row is an absolute difference; ratio rows can be read as percentage changes. Intervals are two-sided 95% Wald CIs. FDR adjustment is across all nine metrics; adjusted p-values would be bold at 0.050.

**Supplementary Table S13.** Chronotype and timing.

| Metric | Instrument | Signed association, h (95% CI) | Raw p | FDR-adjusted p | Fitted sample: participants / days / observations / hours / sites |
| --- | --- | --- | --- | --- | --- |
| Midpoint of the brightest 10 hours | MCTQ MSFsc | +0.205 (+0.045 to +0.366) | **0.010** | **0.017** | 140 / 810 / 810 / 18711.7 h / 9 |
| Midpoint of the brightest 10 hours | MEQ | -0.278 (-0.451 to -0.105) | **0.001** | **0.002** | 141 / 816 / 816 / 18851.0 h / 9 |
| Midpoint of the darkest 10 hours | MCTQ MSFsc | +0.276 (+0.118 to +0.435) | **<0.001** | **0.003** | 140 / 810 / 810 / 18711.7 h / 9 |
| Midpoint of the darkest 10 hours | MEQ | -0.314 (-0.487 to -0.142) | **<0.001** | **0.001** | 141 / 816 / 816 / 18851.0 h / 9 |
| First light timing above 250 lx melEDI | MCTQ MSFsc | +0.381 (+0.146 to +0.616) | **0.001** | **0.003** | 139 / 722 / 722 / 16716.3 h / 9 |
| First light timing above 250 lx melEDI | MEQ | -0.444 (-0.699 to -0.189) | **<0.001** | **0.001** | 140 / 727 / 727 / 16832.5 h / 9 |
| Last light timing above 250 lx melEDI | MCTQ MSFsc | -0.025 (-0.264 to +0.214) | 0.811 | 0.811 | 140 / 683 / 683 / 15900.8 h / 9 |
| Last light timing above 250 lx melEDI | MEQ | +0.029 (-0.236 to +0.294) | 0.814 | 0.814 | 141 / 687 / 687 / 15995.0 h / 9 |
| Midpoint of the longest continuous period above 250 lx melEDI | MCTQ MSFsc | +0.161 (-0.113 to +0.434) | 0.227 | 0.284 | 131 / 478 / 478 / 11325.5 h / 9 |
| Midpoint of the longest continuous period above 250 lx melEDI | MEQ | -0.228 (-0.521 to +0.064) | 0.108 | 0.135 | 132 / 482 / 482 / 11419.8 h / 9 |

MCTQ associations are per one-hour later MSFsc; MEQ associations are per 10 points greater morning preference. Positive associations indicate later timing and negative associations earlier. Intervals are 95% Wald CIs. Raw p-values are bold when raw $p < 0.050$ ; adjusted p-values are bold when FDR-adjusted $p < 0.050$ within the instrument-specific five-outcome family.

**Supplementary Table S14.** Age and biological sex.

| Placement and role | Predictor | Metric | Practical association (95% CI) | Raw p | FDR-adjusted p | Sample |
| --- | --- | --- | --- | --- | --- | --- |
| Near eye (primary) | Age, per 10 years | Brightest 10 h mean | 1.31× (1.09–1.58) | 0.003 | **0.018** | 141 participants; 816 participant-days/observations |
| Near eye (primary) | Age, per 10 years | Time above 1,000 lx melEDI | 1.27× (1.11–1.45) | <0.001 | **0.009** | 141 participants; 816 participant-days/observations |
| Near eye (primary) | Age, per 10 years | melEDI dose | 1.31× (1.09–1.56) | 0.003 | **0.018** | 141 participants; 761 participant-days/observations |
| Chest (complementary) | Age, per 10 years | Mean melEDI | 1.16× (1.04–1.30) | 0.006 | **0.017** | 154 participants; 902 participant-days/observations |
| Chest (complementary) | Age, per 10 years | Brightest 10 h mean | 1.35× (1.15–1.57) | <0.001 | **<0.001** | 154 participants; 902 participant-days/observations |
| Chest (complementary) | Age, per 10 years | Time above 1,000 lx melEDI | 1.31× (1.18–1.46) | <0.001 | **<0.001** | 154 participants; 902 participant-days/observations |
| Chest (complementary) | Age, per 10 years | Time above 250 lx melEDI during wake | 1.16× (1.06–1.26) | 0.001 | **0.004** | 154 participants; 818 participant-days/observations |
| Chest (complementary) | Age, per 10 years | Longest continuous period above 250 lx melEDI | 1.18× (1.09–1.27) | <0.001 | **<0.001** | 154 participants; 902 participant-days/observations |
| Chest (complementary) | Age, per 10 years | melEDI dose | 1.40× (1.21–1.61) | <0.001 | **<0.001** | 154 participants; 851 participant-days/observations |
| Chest (complementary) | Measured biological sex, Female minus Male | Mean melEDI | 0.74× (0.60–0.92) | 0.006 | **0.048** | 154 participants; 902 participant-days/observations |
| Chest (complementary) | Measured biological sex, Female minus Male | Darkest 10 h mean | 0.76× (0.65–0.90) | <0.001 | **0.016** | 154 participants; 902 participant-days/observations |

Associations are site-adjusted estimates per 10-year age increase or Female minus Male. Ratios are back-transformed to the practical scale. Each FDR-adjusted p-value belongs to its sensor-position- and predictor-specific complete 17-test family; bold values meet the FDR-adjusted $p < 0.050$ rule. Samples give the exact fitted participants and participant-day observations for these retained daily-response results.

**Supplementary Table S15.** Biological-sex-specific daily curves and global tests.

|  | Role | Test | F statistic | Numerator df | Denominator df | Raw p | FDR-adjusted p | Support |
| --- | --- | --- | --- | --- | --- | --- | --- | --- |
| Near eye | Primary | Complete Female-minus-Male 24-hour curve | 20.946 | 9.692 | 131 | **0.028** | **0.028** | Supported |
| Chest | Complementary | Complete Female-minus-Male 24-hour curve | 4.876 | 1.003 | 152 | **0.029** | **0.029** | Supported |

Primary near-eye fitted sample: 141 participants, 816 participant-days, 37,756 30-minute observations, and 9 sites. Complementary chest fitted sample: 154 participants, 902 participant-days, 41,842 30-minute observations, and 8 sites.
Each row is one joint complete-curve test. Raw p is bold at raw $p < 0.050$ ; FDR-adjusted p is bold independently at adjusted $p < 0.050$ . Each labelled global family contains one test, so the stored raw and adjusted values coincide.

