## Supplementary Methods: Preregistration deviations for "The health-relevant architecture of the everyday light exposome"

What changed from the registered plan, why it changed, and how to interpret the current analyses

### How to read this page

A preregistration records the analysis plan before the results are known. A deviation is a scientifically relevant difference between that plan and the analysis now used. This page states those differences directly. The result and preparation pages link to the relevant entries.

The first section describes scientific deviations. The second explains the eligibility sensitivities for the age-related and employment-related hypotheses and the population to which those results apply.

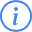
 Current interpretation

Use **Current analysis** and **What this means for interpretation** to understand the reported analysis. The **Preregistered or expected** field shows what differed from the registered or anticipated analysis.

#### Statistical terms used across entries

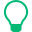
 Short glossary

- A **site-average estimate** is an average across sites that gives each site equal weight.
- A **predictor-by-site interaction** allows the association with a predictor to differ among study sites.
- A **false-discovery-rate (FDR) adjustment** accounts for a declared family of statistical tests; later entries use **FDR**.
- **Model checks** (model diagnostics) assess whether the fitted model is adequate for its intended interpretation.
- **Participant-cluster-robust uncertainty** allows repeated observations from the same participant to remain associated without treating them as independent.
- A **sensitivity analysis** repeats an analysis after one stated choice is changed to assess whether the interpretation is stable.

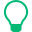
 Time curves and transformed results

- A **nonlinear generalized additive model (GAM) analysis** lets the estimated exposure pattern vary flexibly over time rather than forcing a straight line; later entries may use **GAM**.
- **AR(1)** is a model for autocorrelation in which observations closer together in a sequence are expected to be more similar; the similarity decreases with separation.
- A **derivative** is the estimated rate at which a fitted curve is changing at a given point.
- A **Shapley allocation** distributes fitted-model variation among model components by averaging their added contribution across possible entry orders.
- A **back-transformed** quantity has been converted from the model’s transformed scale to an interpretable response-scale quantity, such as a ratio or time difference.
- A **95% confidence interval (95% CI)** gives the range of parameter values compatible with the estimate under the stated model and uncertainty procedure.

#### Light-exposure measures named in entries

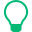
 Measure glossary

- **Melanopic equivalent daylight illuminance (melEDI)** describes light in terms of its stimulation of the melanopsin pathway and is reported in lux.
- **M10** and **L10** describe the brightest and darkest supported consecutive 10-hour windows, respectively.
- **Melanopic daylight efficacy ratio (MDER)** is the mean of viable one-minute ratios of melEDI to photopic illuminance in the current analysis.
- **Interdaily stability (IS)** describes regularity across days; **intradaily variability (IV)** describes fragmentation across adjacent hours within days.
- **MCTQ** and **MEQ** are distinct chronotype measures: corrected midsleep timing from the Munich Chronotype Questionnaire and morningness preference from the Morningness–Eveningness Questionnaire.
- **LEBA** is the Light Exposure Behaviour Assessment, and **VLSQ-8** is an eight-item self-report measure of visual light sensitivity.

### Current scientific deviations

These entries describe changes relevant to what was measured, analysed, or inferred. They are grouped by their role in the scientific analysis, not by the date on which the difference was documented.

#### Study population, samples, placement, and context

##### All hypotheses; placement

**Topic.** All hypotheses; placement

**Preregistered or expected.** Chest data primary; repeat with glasses for robustness

**Current analysis.** Near-eye glasses measurements are primary for the ocular-exposure estimand. Chest measurements are complementary and are compared on placement-matched samples when a direct placement comparison is shown.

**Why.** The placements measure related but non-interchangeable light fields, so simple pooling or unmatched identity comparisons are not valid.

**Affected hypotheses.** All hypotheses (H01–H11)

**What this means for interpretation.** Primary conclusions come from near-eye data; chest evidence is labelled complementary and does not replace the primary estimand.

##### Site sample size and placement availability

**Topic.** Site sample size and placement availability

**Preregistered or expected.** Target 15-30 participants per site using all eligible repository participants

**Current analysis.** Recruitment and placement availability remain uneven by site. Every reported hypothesis analysis now reports its exact participants, participant-days or participant-hours, observations, placements, and sites rather than presenting one universal sample size.

**Why.** Uneven support is an observed feature of the study design and cannot be repaired analytically; exact model-specific denominators and complementary-placement labels are the specified response.

**Affected hypotheses.** All hypotheses (H01–H11)

**What this means for interpretation.** Site and placement comparisons must be interpreted with their model-specific support, especially for Munich (DE), San José (CR), and glasses-versus-chest comparisons.

##### Metric construct; full-day and sleep

**Topic.** Metric construct; full-day and sleep

**Preregistered or expected.** The intended 24-hour environmental-light record combines worn near-eye waking measurements with bedside sleep-environment measurements during the diary-defined sleep window

**Current analysis.** The full-day hybrid record combines worn waking measurements with retained bedside sleep-environment measurements and is described explicitly as such.

**Why.** This preserves the practical 24-hour environmental record without mislabelling sleep values as ocular exposure.

**Affected hypotheses.** All hypotheses (H01–H11)

**What this means for interpretation.** Full-day metrics are hybrid personal/environmental summaries; sleep-specific interpretations remain environmental.

##### Placement pooling

**Topic.** Placement pooling

**Preregistered or expected.** No pooled placement model was preregistered; chest was primary and glasses a robustness repeat

**Current analysis.** Simple pooling of near-eye and chest measurements is rejected; placements retain separate estimands.

**Why.** The placements differ systematically and are not duplicate measures of one interchangeable quantity.

**Affected hypotheses.** All hypotheses (H01–H11)

**What this means for interpretation.** No pooled placement claim is permitted.

##### Chest complementary analysis

**Topic.** Chest complementary analysis

**Preregistered or expected.** Registered glasses analysis was a robustness repeat

**Current analysis.** Direct near-eye/chest displays use placement-matched participant, local-date, and clock keys, while chest remains complementary in all-available analyses.

**Why.** Matching makes the compared placements refer to the same observed periods and avoids conflating placement with sample composition.

**Affected hypotheses.** All hypotheses (H01–H11)

**What this means for interpretation.** Matched comparisons describe placement sensitivity, not equivalence, and unmatched all-available estimates are not plotted against each other as identities.

##### Sleep-period construct language

**Topic.** Sleep-period construct language

**Preregistered or expected.** Registered outcomes referred to chest-mounted measurements, including sleep metrics

**Current analysis.** Retained sleep-period logger values are described as bedside sleep-environment exposure.

**Why.** This language matches what the logger can measure when wear is not verified during sleep.

**Affected hypotheses.** All hypotheses (H01–H11)

**What this means for interpretation.** Sleep-period results must not be called ocular exposure.

#### Data preparation, timing, state, and support

##### Preprocessing; non-wear

**Topic.** Preprocessing; non-wear

**Preregistered or expected.** Remove non-wear periods identified by device logs

**Current analysis.** Logged non-wear outside reported sleep is set to missing, whereas logger values during the reported sleep interval are retained as measurements of the sleep environment and are not described as worn ocular exposure.

**Why.** The retained sleep values answer a different construct question from waking personal exposure; the placement decision and reader reports now distinguish those constructs.

**Affected hypotheses.** All hypotheses (H01–H11)

**What this means for interpretation.** Sleep-environment results describe light at the logger location during attempted sleep, not verified eye-level exposure while the device was worn.

##### Preprocessing; sleep windows

**Topic.** Preprocessing; sleep windows

**Preregistered or expected.** Registered rule refers to sleep but does not define attempted-sleep versus estimated-onset boundary

**Current analysis.** Attempted-sleep time is the operational boundary for sleep-environment and pre-sleep metrics rather than estimated physiological sleep onset.

**Why.** Attempted sleep aligns the analysed interval with the intention to sleep and is now defined consistently in preparation and reader-facing methods.

**Affected hypotheses.** All hypotheses (H01–H11)

**What this means for interpretation.** Sleep and pre-sleep denominators refer to attempted sleep; they should not be interpreted as intervals anchored to measured sleep onset.

##### Preprocessing saturation boundary

**Topic.** Preprocessing saturation boundary

**Preregistered or expected.** Set melanopic EDI values above 120000 lx to missing

**Current analysis.** ActLumus melEDI values at or above 100000 lx are treated as outside the instrument’s valid operating range and set to invalid rather than retained.

**Why.** The strict upper-bound rule follows the specified saturation decision and is applied before metric calculation.

**Affected hypotheses.** All hypotheses (H01–H11)

**What this means for interpretation.** Extremely high invalid readings do not contribute to exposure metrics or model samples.

##### Timestamp integrity; DST fall-back

**Topic.** Timestamp integrity; DST fall-back

**Preregistered or expected.** International site-local timestamps retain their absolute instants and remain uniquely keyed through daylight-saving transitions

**Current analysis.** Timestamp processing uses UTC to preserve real instants and a separate local wall-clock axis for cross-site clock-time summaries. Repeated fall-back minutes are averaged only for clock-aligned profiles and retained as separate real intervals for elapsed-time and sequential metrics.

**Why.** The dual-axis rule prevents daylight-saving transitions from either duplicating wall-clock profiles or erasing real elapsed intervals.

**Affected hypotheses.** All hypotheses (H01–H11)

**What this means for interpretation.** Clock-aligned and elapsed-time metrics can legitimately treat the repeated hour differently; the preparation pages explain which time axis each calculation uses.

##### State precedence; measurement context

**Topic.** State precedence; measurement context

**Preregistered or expected.** The sleep diary defines the relevant sleepprep-to-wake bedside window; wear logs are not sufficiently consistent to define sleep

**Current analysis.** Diary state is authoritative for sleep context. Wear=‘sleep’ records device-log context only; wear=‘off’ invalidates waking measurements outside diary sleep, while off or missing wear during diary sleep and site-leave records are retained according to the specified context rules.

**Why.** The rule separates measurement-context validity from the participant’s diary-defined state.

**Affected hypotheses.** All hypotheses (H01–H11)

**What this means for interpretation.** Waking non-wear is excluded, whereas retained sleep values remain environmental rather than verified worn exposure.

##### H02 and H11 participant-day inclusion

**Topic.** H02 and H11 participant-day inclusion

**Preregistered or expected.** The registration excludes days using declared coverage rules and does not exclude a day merely because its binned outcome is constant

**Current analysis.** H02 and H11 preserve supported constant profiles and exclude only otherwise eligible participant-days whose finite one-minute melEDI values are all exactly zero under the specified upstream plausibility rule.

**Why.** Individual zero observations are valid; the day-level exclusion targets only an implausible complete zero signal and is retained as a data sensitivity.

**Affected hypotheses.** H02 and H11

**What this means for interpretation.** The reported samples do not apply a general constant-profile deletion.

##### Daily 80% coverage denominator

**Topic.** Daily 80% coverage denominator

**Preregistered or expected.** The signed preregistration excludes participant-days below 80% valid wear time with sleep excluded from the daily denominator

**Current analysis.** Primary daily coverage uses finite melEDI minutes across a fixed 1440-wall-minute day for the hybrid waking-plus-bedside-sleep record. Hourly eligibility is applied separately and does not remove finite minutes from daily coverage. Sleep-excluded denominators are fixed sensitivities.

**Why.** The hybrid 24-hour construct is primary; alternative denominators show dependence on sleep handling.

**Affected hypotheses.** All hypotheses (H01–H11)

**What this means for interpretation.** Primary daily metrics intentionally treat unobserved eligible minutes as missing against the full wall-clock day and should be interpreted as hybrid full-day exposure summaries.

##### TUM exercise-diary participant key

**Topic.** TUM exercise-diary participant key

**Preregistered or expected.** Questionnaire and diary source records should be joined under their correct immutable site-participant identifiers

**Current analysis.** The corrected upstream TUM exercise diary assigns the seven affected May 2024 records to TUM_S001, not TUM_S101; no local identifier rewrite is applied.

**Why.** The local exercise-diary input contains the corrected participant key.

**Affected hypotheses.** H06

**What this means for interpretation.** H06 exercise analyses use the authoritative corrected participant identity without altering shared data locally.

##### Participant-day light-signal plausibility

**Topic.** Participant-day light-signal plausibility

**Preregistered or expected.** The preregistration uses coverage and signal-validity rules but does not specify exclusion of an otherwise eligible complete day-long exact-zero melEDI series

**Current analysis.** The main analysis excludes an otherwise eligible participant-day only when every finite one-minute melEDI value is exactly 0 lx. Individual zeros remain valid, the original rows remain available, and retaining these days is a sensitivity analysis.

**Why.** A complete exact-zero day is treated as a signal-plausibility failure rather than ordinary darkness.

**Affected hypotheses.** All hypotheses (H01–H11)

**What this means for interpretation.** Affected samples lose only the documented all-zero participant-days; zero-capable metrics and models otherwise retain zero observations.

#### Metric and outcome definitions

##### Metric definition; darkest level window

**Topic.** Metric definition; darkest level window

**Preregistered or expected.** Geometric mean melanopic EDI during the 5 darkest hours

**Current analysis.** The darkest-window level metric is the geometric mean melEDI during the ten darkest supported hours (L10 mean); no L5 level metric is used.

**Why.** The ten-hour window is the specified operational definition and is subject to the current complete-window support rule.

**Affected hypotheses.** H01, H05, H06, H07, H08, and H10

**What this means for interpretation.** Results labelled L10 mean refer to a ten-hour darkest supported window and must not be interpreted as L5.

##### Melanopic EDI dose

**Topic.** Melanopic EDI dose

**Preregistered or expected.** Daily dose should represent an adequately supported integration interval

**Current analysis.** Melanopic EDI dose is defined as observed actual-duration dose, with the specified time-sensitive correction applied only when profile-weighted coverage reaches 0.80. It is not treated as an unqualified complete-day sum when exposure is missing.

**Why.** The rule preserves the observed integration interval and avoids universal scaling over missing low-exposure night periods.

**Affected hypotheses.** H01, H05, H06, H07, H08, and H10

**What this means for interpretation.** Dose scale and model samples depend on the duration and temporal coverage of the valid measurements.

##### IS and IV temporal support

**Topic.** IS and IV temporal support

**Preregistered or expected.** IS and IV should quantify repeated-day regularity and true adjacent-hour fragmentation without bridging unobserved time

**Current analysis.** IS and IV use explicit one-hour grids and true temporal support: at least three eligible days, at least 20 repeated clock hours for IS, and at least 24 true adjacent pairs across three days for IV; repeated fall-back hours collapse only for IS.

**Why.** These rules prevent missing intervals from being treated as observed regularity or adjacency.

**Affected hypotheses.** H01, H05, H06, and H10

**What this means for interpretation.** Values and estimability depend on observed repeated-day support and true adjacent-hour pairs, without bridging gaps.

##### Window, continuous-period, and timing support

**Topic.** Window, continuous-period, and timing support

**Preregistered or expected.** M10/L10, continuous threshold periods, and threshold timings should represent supported contiguous clock intervals without compressing gaps

**Current analysis.** M10/L10 require 20 consecutive supported 30-minute bins with deterministic ties and midnight wrap for L10; continuous periods terminate at gaps; mean timing is circular and duration-weighted; first and last timing require boundary support.

**Why.** The rules preserve clock continuity and prevent incomplete windows or observed edges from being treated as complete events.

**Affected hypotheses.** H01, H05, H06, H07, H08, H09, and H10

**What this means for interpretation.** Window levels, durations, timings, and analytical samples depend on the temporal support required by each metric. The alternative preprocessing sensitivity assesses dependence on gap handling.

##### MDER metric definition and support

**Topic.** MDER metric definition and support

**Preregistered or expected.** MDER summarizes momentary MEDI/LIGHT ratios; the preregistration does not specify a ratio-of-integrals definition

**Current analysis.** MDER is the arithmetic mean of viable one-minute MEDI/LIGHT ratios. A minute is viable only when both channels are finite and strictly positive; a day requires at least 720 viable minutes on the fixed 1440-minute local wall-clock grid, otherwise MDER is missing for that day.

**Why.** Averaging momentary ratios across viable measurement pairs preserves the intended MDER construct; a ratio of integrals defines a different quantity.

**Affected hypotheses.** H01, H05, H06, and H10

**What this means for interpretation.** MDER represents the average spectral ratio during viable paired minutes, not a ratio of daily totals; metric-specific missingness and model samples follow the 50% viable-minute rule.

##### H01 L10 midpoint response conversion

**Topic.** H01 L10 midpoint response conversion

**Preregistered or expected.** A noon-based linearization represents L10 midpoint clock times strictly after noon on the preceding negative-hour scale

**Current analysis.** For H01 L10 midpoint models, clock times strictly after 16:00 are represented on the preceding negative-hour scale; a noon-based conversion is retained as a same-row, same-model sensitivity.

**Why.** The primary conversion avoids splitting the darkest-window midpoint across midnight in the fitted linear timing model.

**Affected hypotheses.** H01

**What this means for interpretation.** H01 L10 midpoint coefficients depend on the declared circular-to-linear representation and are interpreted with its sensitivity.

##### H01 time below 10 lx melEDI before sleep

**Topic.** H01 time below 10 lx melEDI before sleep

**Preregistered or expected.** The outcome was described as time below 10 lx melEDI in the three hours before sleep and was checked against a three-hour ceiling

**Current analysis.** Time below 10 lx melEDI before sleep is the cumulative duration across all diary-defined pre-sleep intervals on the local calendar date; no three-hour or six-hour cap is imposed, and values above six hours trigger a diagnostic warning only.

**Why.** The stated definition makes the temporal denominator explicit.

**Affected hypotheses.** H01

**What this means for interpretation.** The metric can exceed six hours and should not be interpreted as exposure within a fixed pre-sleep window.

#### Hypothesis models, estimands, and inference

##### H01 model

**Topic.** H01 model

**Preregistered or expected.** Metric ~ Site + Latitude + participant-within-site random intercept; photoperiod for day/night duration metrics

**Current analysis.** H01 now evaluates overall site, photoperiod, absolute latitude, and same-frame site-versus-linear-latitude adequacy as separate prespecified model-level questions with complete 17-metric FDR families; it does not select between separate site and latitude models by AIC.

**Why.** Site and latitude are structurally linked because each site has one latitude. The reported analysis separates the questions and uses the adequacy comparison to show when a linear latitude gradient does not capture site differences.

**Affected hypotheses.** H01

**What this means for interpretation.** Site effects and latitude gradients are not mutually interchangeable, and an adequacy result is not a residual site effect after latitude adjustment.

##### H02 smooth basis

**Topic.** H02 smooth basis

**Preregistered or expected.** Cyclic site-specific time smooth with bs=‘cc’

**Current analysis.** H02 uses a cyclic common time smooth with sum-to-zero site time deviations whose marginal basis is not forced cyclic; a fully cyclic site and participant formulation is retained as a model-form sensitivity.

**Why.** The reported basis comparison favoured the cyclic common smooth and found no residual benefit from replacing it with a non-cyclic global smooth.

**Affected hypotheses.** H02

**What this means for interpretation.** Midnight continuity is enforced for the global curve; site-deviation topology is an acknowledged model-form choice tested in sensitivity analysis.

##### H02 model structure

**Topic.** H02 model structure

**Preregistered or expected.** Registered formula does not state a separate overall time smooth

**Current analysis.** H02 uses a cyclic common time smooth, sum-to-zero site time deviations, participant-specific time curves, and a participant-day intercept with boundary-aware AR(1) sequences.

**Why.** The hierarchy was selected through the H02 model evaluation and is inherited by H11.

**Affected hypotheses.** H02

**What this means for interpretation.** Variation and uncertainty are partitioned across time, site, participant, and participant-day under this richer hierarchy rather than the simpler registered structure.

##### H02 outcome and epoch

**Topic.** H02 outcome and epoch

**Preregistered or expected.** Hourly geometric mean melanopic EDI

**Current analysis.** H02 models arithmetic-mean melEDI in supported 30-minute bins as log10(melEDI + 0.1 lx), while the registered hourly geometric-mean outcome remains documented as the preregistered specification.

**Why.** The 30-minute zero-aware outcome preserves the temporal curve and exact zeros needed by the reported model and diagnostic workflow.

**Affected hypotheses.** H02

**What this means for interpretation.** H02 estimates, sample counts, and autocorrelation refer to supported 30-minute arithmetic-mean observations, not registered hourly geometric means.

##### H03-H04 site structure

**Topic.** H03-H04 site structure

**Preregistered or expected.** Predictor random slope by site

**Current analysis.** H03 and H04 use fixed study-site adjustment for the population-average category association and fit separate category-by-site interaction models to evaluate variation among sites.

**Why.** The available number and support of sites did not justify the registered random-slope formulation. Separating the average association from the site interaction makes the two questions explicit.

**Affected hypotheses.** H03 and H04

**What this means for interpretation.** Primary effects describe the study-site-average association; site interactions are separate tests and do not redefine the primary omnibus test.

##### H03-H04 error distribution

**Topic.** H03-H04 error distribution

**Preregistered or expected.** Registered as LMM without a non-Gaussian family specified

**Current analysis.** H03 and H04 use fixed-power quasi-Tweedie log-mean models with participant-cluster robust covariance rather than Gaussian linear mixed models.

**Why.** The response is non-negative, right-skewed, and includes exact zeros. The reported working mean models are supported by targeted diagnostics but are not treated as calibrated zero-generating distributions.

**Affected hypotheses.** H03 and H04

**What this means for interpretation.** Effects are multiplicative mean comparisons on the response scale; residual dependence and zero-mass mismatch remain stated limitations.

##### H06 outcome

**Topic.** H06 outcome

**Preregistered or expected.** Daily light-exposure metrics

**Current analysis.** H06 analyses supported hourly geometric-mean melEDI rather than the preregistered daily metric set.

**Why.** The hourly formulation supports a parsimonious, directly interpretable association with temporal context.

**Affected hypotheses.** H06

**What this means for interpretation.** The main H06 result concerns hourly melEDI and repeated supported hours; it must not be presented as the preregistered daily-metric estimand.

##### H07 predictors and outcome selection

**Topic.** H07 predictors and outcome selection

**Preregistered or expected.** Absolute-latitude-by-photoperiod tensor applied to all level, duration, and exposure-history metrics

**Current analysis.** H07 retains all nine eligible metrics. The registered latitude-by-photoperiod surface is reported as non-identifiable in this design, and a separately labelled site-adjusted nonlinear photoperiod analysis provides descriptive evidence, including a derivative-defined plateau pattern.

**Why.** Absolute latitude is determined by site and the observed joint support cannot identify the registered surface. The adapted analysis avoids significance-based outcome selection and avoids a mechanistic ceiling claim.

**Affected hypotheses.** H07

**What this means for interpretation.** The nonlinear patterns are descriptive, sensitive to model and site choices, and are not causal latitude effects or evidence of a physiological ceiling.

##### H01 predictor scope

**Topic.** H01 predictor scope

**Preregistered or expected.** Photoperiod adjustment was explicitly registered only for metrics with a day/night-duration component

**Current analysis.** H01 includes photoperiod in all 17 primary metric models and retains the narrower literal preregistration scope as a named sensitivity.

**Why.** A common primary adjustment set makes the analyses comparable; a sensitivity follows the literal registered predictor scope.

**Affected hypotheses.** H01

**What this means for interpretation.** Primary H01 estimates are photoperiod-adjusted even for metrics where the preregistration named it only selectively.

##### H01 response models

**Topic.** H01 response models

**Preregistered or expected.** H01 registered linear mixed models for the metric outcomes

**Current analysis.** H01 uses a specified metric-specific set of Gaussian, transformed-Gaussian, and Tweedie models with fixed diagnostic criteria and prespecified alternatives across data scenarios.

**Why.** The 17 metrics differ in scale, bounds, exact-zero mass, and tail behaviour, so one universal Gaussian mixed model is not adequate.

**Affected hypotheses.** H01

**What this means for interpretation.** Effect scales and diagnostic limitations differ by metric and are reported explicitly.

##### H02 photoperiod representation

**Topic.** H02 photoperiod representation

**Preregistered or expected.** H02 registered time site participant pattern and an AR(1) correction; it did not register a categorical day/night-state temporal deviation

**Current analysis.** The H02 primary model does not include a categorical day/night-state smooth as a representation of photoperiod.

**Why.** A categorical day/night state does not represent daily photoperiod duration.

**Affected hypotheses.** H02

**What this means for interpretation.** H02 temporal results are not adjusted for a day/night-state smooth labelled as photoperiod.

##### H02 hierarchy

**Topic.** H02 hierarchy

**Preregistered or expected.** H02 registered participant-time patterns within sites

**Current analysis.** H02 uses participant-specific time curves, a participant-day random intercept, and boundary-aware AR(1) sequence starts.

**Why.** The repeated 30-minute structure requires participant, day, and within-sequence dependence beyond the registered simple participant effect.

**Affected hypotheses.** H02

**What this means for interpretation.** Uncertainty and fitted variation reflect this hierarchy; H11 inherits it.

##### H03 light-source categories

**Topic.** H03 light-source categories

**Preregistered or expected.** H03 registered all prespecified hourly primary light-source categories

**Current analysis.** H03 retains seven prespecified light-source categories under one placement-independent support rule: indoor electric light, outdoor electric light, indoor daylight, outdoor daylight, emissive display light, darkness during sleep, and external light during sleep.

**Why.** The same category definitions apply to both placements, and unsupported category-site cells are reported explicitly.

**Affected hypotheses.** H03

**What this means for interpretation.** The H03 association pertains to the seven retained categories; sparse site-category cells are not silently removed to change the construct.

##### H03 primary test estimand

**Topic.** H03 primary test estimand

**Preregistered or expected.** H03 registered a light-source association with predictor variation by site

**Current analysis.** H03 uses the additive site-plus-light-source model for the primary six-restriction category omnibus and a separate category-by-site interaction model for site-specific variation and descriptive estimates.

**Why.** Separating the two models isolates the study-site-average category association from the interaction question.

**Affected hypotheses.** H03

**What this means for interpretation.** The primary omnibus is not a joint association-plus-interaction test; site-varying summaries are secondary to that primary test.

##### H03 multiplicity and contrasts

**Topic.** H03 multiplicity and contrasts

**Preregistered or expected.** H03 requires FDR control within a declared hypothesis family

**Current analysis.** H03 applies FDR correction to explicitly assembled complete contrast families; category and site-interaction questions are labelled separately and no hidden by-group adjustment is described as a single family.

**Why.** Vector-wide adjustment keeps each contrast within its declared complete comparison family.

**Affected hypotheses.** H03

**What this means for interpretation.** Highlighted contrasts depend on the declared complete family rather than software defaults.

##### H04 activity categories

**Topic.** H04 activity categories

**Preregistered or expected.** H04 registered all prespecified hourly activity categories

**Current analysis.** H04 converts selected activity flags to a long activity variable, collapses outdoor work, outdoor free time, and open-air travel into Outdoors before pivoting, removes duplicate labels within hour, gives each co-selected category weight 1/k, and retains Other only when it is the sole selected category for descriptive display.

**Why.** This construction preserves multi-label hours without allowing them to contribute more total weight than single-label hours, while combining sparse conceptually related outdoor activities.

**Affected hypotheses.** H04

**What this means for interpretation.** H04 estimates compare the defined activity categories; Other is not part of the primary five-category omnibus and multi-label hours are fractionally weighted.

##### H04 primary test estimand

**Topic.** H04 primary test estimand

**Preregistered or expected.** H04 registered an activity association with predictor variation by site

**Current analysis.** H04 uses the additive site-plus-activity model for the primary activity omnibus and a separate activity-by-site interaction model for site-specific variation and descriptive estimates.

**Why.** The separation isolates the study-site-average activity association from variation among sites.

**Affected hypotheses.** H04

**What this means for interpretation.** The primary result is not a joint main-plus-interaction test.

##### H04 multiplicity and contrasts

**Topic.** H04 multiplicity and contrasts

**Preregistered or expected.** H04 requires FDR control within a declared hypothesis family

**Current analysis.** H04 applies FDR correction to explicitly assembled complete activity and site-interaction contrast families rather than relying on hidden by-groups or default Dunnett adjustments.

**Why.** Explicit family definitions keep every displayed adjusted p-value tied to the complete set of comparisons.

**Affected hypotheses.** H04

**What this means for interpretation.** Highlighted activity/site comparisons reflect the declared full families.

##### H05 registered model

**Topic.** H05 registered model

**Preregistered or expected.** H05 registered correlation matrices plus a site-adjusted Metric ~ LEBA + (1|Site) model

**Current analysis.** H05 uses site-adjusted association models for each personal light-exposure metric and LEBA factor rather than participant-level pooled bivariate correlations.

**Why.** Site adjustment preserves the registered scientific question while the reported fixed-site implementation supplies a stable population-average estimate and a separate site-interaction assessment.

**Affected hypotheses.** H05

**What this means for interpretation.** H05 effects are site-adjusted model associations, not pooled correlation coefficients.

##### H05 metric-factor set and multiplicity

**Topic.** H05 metric-factor set and multiplicity

**Preregistered or expected.** The registration refers to selected metrics and LEBA factors with FDR within H05 but does not enumerate the complete family

**Current analysis.** H05 analyses four LEBA factors across 17 metrics and treats each declared 68-test vector as a complete FDR family, including non-significant members.

**Why.** Retaining the full factor-by-metric grid controls multiplicity across the complete set of associations.

**Affected hypotheses.** H05

**What this means for interpretation.** Individual raw associations are interpreted only in the context of their complete 68-test family; the reported analysis retains no primary association after adjustment.

##### H06 predictor scope

**Topic.** H06 predictor scope

**Preregistered or expected.** H06 registered weekday/weekend and free/work day, daily exercise, and sleep onset, wake time, and duration as predictors of daily exposure metrics

**Current analysis.** H06 uses three primary predictors: work versus free day, Active versus Sedentary day, and previous-night sleep duration. Weekday/weekend is a sensitivity; other registered sleep and exercise predictors are exploratory rather than members of the primary family.

**Why.** This narrower predictor set was used for parsimony and interpretability in the hourly adaptation.

**Affected hypotheses.** H06

**What this means for interpretation.** The primary H06 claim is limited to these three predictors and cannot be generalized to all preregistered sleep or exercise variables.

##### H06 model and site structure

**Topic.** H06 model and site structure

**Preregistered or expected.** H06 registered metric-specific daily models with measure random slopes by site and participant nested in site

**Current analysis.** H06 fits a zero-inclusive quasi-Poisson log-mean model with fixed site, participant-cluster HC3 covariance, and finite-cluster inference. The primary additive model estimates study-site-average associations; separate predictor-by-site models evaluate interactions. Local clock is reserved for a distinct exploratory nonlinear analysis.

**Why.** The robust population-average route addresses repeated supported hours without claiming a Gaussian individual-level error model; the separate nonlinear analysis describes time of day.

**Affected hypotheses.** H06

**What this means for interpretation.** Primary ratios are marginal mean comparisons for supported hourly melEDI. Residual zero mass, clustering, site dependence, and temporal-model limitations remain explicit.

##### H06 multiplicity and contrasts

**Topic.** H06 multiplicity and contrasts

**Preregistered or expected.** H06 requires FDR control within a declared hypothesis family

**Current analysis.** H06 uses separately declared complete FDR families for the three primary average associations, three site interactions, three practical contrasts, and corresponding gap-timing-unaware sensitivity families; exploratory site screens are labelled separately.

**Why.** Separate complete families distinguish average effects from interactions.

**Affected hypotheses.** H06

**What this means for interpretation.** Adjusted decisions must be read within their declared three-member family and exploratory site screens cannot override a non-retained interaction.

##### H07 outcome selection and multiplicity

**Topic.** H07 outcome selection and multiplicity

**Preregistered or expected.** H07 registered all nine eligible level, duration, and exposure-history outcomes with FDR within H07

**Current analysis.** H07 retains the complete nine-metric outcome set and does not select metrics using H01 significance or H07 AIC. Non-identifiable or diagnostically limited planned members remain visible rather than being dropped from the family.

**Why.** Keeping the complete family prevents outcome selection based on significance in H01 or H07.

**Affected hypotheses.** H07

**What this means for interpretation.** The nonlinear descriptive evidence must be interpreted across all nine planned metrics, including non-identifiable and limited branches.

##### H07 response model

**Topic.** H07 response model

**Preregistered or expected.** The registered H07 GAMM applies to metrics with different units and support properties

**Current analysis.** H07 uses metric-specific response specifications and diagnostics rather than one universal zero-aware log-Gaussian model for level, duration, and dose outcomes.

**Why.** The metrics differ in units, bounds, exact-zero mass, and tail behaviour; the reported working families are chosen and assessed per metric.

**Affected hypotheses.** H07

**What this means for interpretation.** Model adequacy and the meaning of changes differ by metric, and the sleep-environment Tweedie branch remains descriptive because its zero-mass diagnostic fails.

##### H08 primary test estimand

**Topic.** H08 primary test estimand

**Preregistered or expected.** H08 registered Metric ~ VLSQ8 * Site with the VLSQ-8 association and site heterogeneity distinguishable

**Current analysis.** H08 estimates the study-site-average visual-light-sensitivity association separately from the visual-light-sensitivity-by-site interaction for each metric.

**Why.** Separate nested comparisons preserve the registered distinction between an average association and variation among sites.

**Affected hypotheses.** H08

**What this means for interpretation.** A non-significant average association is not a joint statement that both the average and every site difference are absent.

##### H08 multiplicity

**Topic.** H08 multiplicity

**Preregistered or expected.** H08 requires FDR within a declared family across nine outcomes

**Current analysis.** H08 uses separate complete nine-metric FDR families for average associations and site interactions; scalar adjustments are not used.

**Why.** Separate complete families distinguish the average-association question from variation among sites.

**Affected hypotheses.** H08

**What this means for interpretation.** No H08 association or site interaction is retained after its declared family adjustment.

##### H09 chronotype predictors

**Topic.** H09 chronotype predictors

**Preregistered or expected.** H09 registered both MCTQ and MEQ chronotype predictors

**Current analysis.** H09 includes both MCTQ midsleep on free days corrected for sleep debt and MEQ morningness score as distinct chronotype constructs, with separate models and families.

**Why.** The available aggregate fields represent distinct instruments and are analysed separately.

**Affected hypotheses.** H09

**What this means for interpretation.** MCTQ effects are per one hour later timing; MEQ effects are per ten points greater morning preference. They are not interchangeable measures.

##### H09 site adjustment

**Topic.** H09 site adjustment

**Preregistered or expected.** H09 registered Metric ~ chronotype * Site with participant nested in site

**Current analysis.** H09 reports site-adjusted main chronotype models and evaluates chronotype-by-site interactions separately; estimates and intervals come from the declared fitted model for each instrument and timing outcome.

**Why.** Using a fixed formula and sample for each reported estimate avoids significance-driven model selection.

**Affected hypotheses.** H09

**What this means for interpretation.** Chronotype slopes are study-site-adjusted associations; site interactions are a separate question.

##### H09 multiplicity and model selection

**Topic.** H09 multiplicity and model selection

**Preregistered or expected.** H09 requires within-hypothesis FDR for five timing outcomes crossed with both registered chronotype instruments and site interactions

**Current analysis.** H09 uses four separate complete five-outcome FDR families: MCTQ average associations, MEQ average associations, MCTQ-by-site interactions, and MEQ-by-site interactions. Model selection is not driven by adjusted significance.

**Why.** The family partition reflects the two instruments and the distinction between average and interaction tests.

**Affected hypotheses.** H09

**What this means for interpretation.** Each reported adjusted p-value belongs to one named five-member family; non-estimable interaction branches remain visible.

##### H10 sex-by-site interaction

**Topic.** H10 sex-by-site interaction

**Preregistered or expected.** H10 registered age and biological-sex predictors each interacting with site

**Current analysis.** H10 now fits, tests, and reports both age-by-site and biological-sex-by-site interactions for all planned metrics, with non-estimable branches explicitly identified rather than omitted.

**Why.** This retains the full registered interaction scope while keeping sensor placements separate.

**Affected hypotheses.** H10

**What this means for interpretation.** H10 can distinguish average age/sex associations from variation of those associations among sites; sparse interactions remain qualified by diagnostics.

##### H10 multiplicity

**Topic.** H10 multiplicity

**Preregistered or expected.** H10 requires within-hypothesis FDR for age and sex associations and their site interactions

**Current analysis.** H10 uses four separate complete 17-metric FDR families for age average associations, biological-sex average associations, age-by-site interactions, and biological-sex-by-site interactions.

**Why.** The partition was fixed in the reported analysis and all planned slots, including non-estimable ones, are retained.

**Affected hypotheses.** H10

**What this means for interpretation.** The current conclusion of 11 primary average associations and two chest age-by-site interactions is evaluated within these complete families.

##### H11 outcome and epoch

**Topic.** H11 outcome and epoch

**Preregistered or expected.** H11 registered hourly geometric-mean melanopic EDI

**Current analysis.** H11 deliberately inherits H02’s zero-aware transformed arithmetic-mean melEDI in 30-minute bins rather than the registered hourly geometric-mean outcome.

**Why.** Using the H02 temporal frame provides consistent epochs, temporal support, gap handling, autocorrelation, and placement definitions for the sex-specific curve analysis.

**Affected hypotheses.** H11

**What this means for interpretation.** H11 estimates sex-specific 30-minute daily curves on the inherited transformed scale; its sample size and dependence structure differ from the preregistered hourly analysis.

##### H11 sex estimand

**Topic.** H11 sex estimand

**Preregistered or expected.** The registered H11 full model contains both a parametric sex effect and a sex-specific cyclic time smooth

**Current analysis.** The reported H11 full model contains a parametric biological-sex level term and a cyclic sex-specific time-deviation smooth. The primary test evaluates the complete level-plus-shape Female-minus-Male curve.

**Why.** This construction restores the registered distinction between a time-constant level difference and a time-varying shape difference while making their joint curve the primary estimand.

**Affected hypotheses.** H11

**What this means for interpretation.** The global result is about the whole sex-specific curve; secondary level and shape tests are attribution analyses and do not replace it.

##### H11 temporal and site smooths

**Topic.** H11 temporal and site smooths

**Preregistered or expected.** H11 registered cyclic sex smooths, a site factor smooth, and a site random effect

**Current analysis.** H11 uses the H02 temporal architecture: a cyclic global time smooth, a cyclic biological-sex deviation, sum-to-zero site time deviations, participant-specific time curves, and a participant-day intercept. The separate registered site random intercept is replaced by this inherited site and participant hierarchy.

**Why.** The inherited architecture is evaluated in H02 and checked for H11 with boundary-aware autocorrelation and robust complete-curve inference.

**Affected hypotheses.** H11

**What this means for interpretation.** Site and participant temporal structure is richer than the preregistered formula; interpretation is conditional on the reported inherited architecture and diagnostics.

##### H11 participant hierarchy

**Topic.** H11 participant hierarchy

**Preregistered or expected.** H11 registered a participant random-effect smooth

**Current analysis.** H11 models participant-specific time curves and participant-day intercepts rather than only a participant random intercept.

**Why.** The reported hierarchy reflects repeated 30-minute observations within days and repeated days within participants and is inherited from H02.

**Affected hypotheses.** H11

**What this means for interpretation.** Uncertainty and weighting account for participant and day structure; variance components refer to the stated hierarchy.

##### H11 contextual activity analysis

**Topic.** H11 contextual activity analysis

**Preregistered or expected.** No activity-adjusted causal or mediation estimand was registered for H11

**Current analysis.** H11 includes an explicitly exploratory same-sample activity-context sensitivity. It first refits the reported model without activity on the activity-complete sample and then adds activity on exactly the same observations.

**Why.** The paired same-sample design separates sample restriction from covariate adjustment as far as the observational data allow.

**Affected hypotheses.** H11

**What this means for interpretation.** The analysis cannot establish mediation or a behavioural mechanism; near-eye support was already lost through sample restriction and the chest attenuation is non-causal context only.

##### H11 curve inference and multiplicity

**Topic.** H11 curve inference and multiplicity

**Preregistered or expected.** H11 registered one temporal sex hypothesis with FDR if multiple tests were used

**Current analysis.** H11 uses one participant-cluster-robust global complete-curve test per placement as the primary decision. Level and shape form a secondary two-test FDR family. Displayed clock-time intervals are pointwise, not simultaneous, and do not define a familywise-significant period.

**Why.** The analysis uses AR-whitened participant-cluster CR1 curve inference with smoothing-bias covariance and a finite-cluster reference distribution.

**Affected hypotheses.** H11

**What this means for interpretation.** The supported global curves may be described, but local clock ranges are descriptive and the secondary decomposition does not cleanly attribute the effect to level or shape.

### Eligibility and generalisation

The eligibility checks below assess the age-related and employment-related hypotheses in restricted samples. They describe the robustness of these analyses while retaining the distinction between the observed study population and the registered target population.

#### Entries

##### Inclusion criteria

**Topic.** Inclusion criteria

**Preregistered or expected.** Protocol age 18–65 years and employment at least 80%.

**Current analysis.** The study population included nine people outside the registered age or employment criteria: one older than 65 years, two recorded as not employed and seven recorded as marginally employed, with one person meeting both the age and employment exclusion criteria. The age and biological-sex analysis (H10) jointly excluded these age and employment categories. Six affected participants contributed to the near-eye models and nine to the chest models; metric-specific samples fell from 137–141 to 131–135 near-eye participants and from 152–154 to 143–145 chest participants. All 11 FDR-supported main findings retained their direction and support, and all 68 main-effect FDR decisions were unchanged. This was a joint age-and-employment restriction, not an age-only refit.

For the day-type, exercise and sleep analysis (H06), a separate near-eye employment sensitivity excluded six people recorded as not employed or marginally employed. The sample fell from 137 to 131 participants, from 715 to 684 participant-days and from 16,596 to 15,871 participant-hours. The additive and predictor-by-site models preserved the directions and FDR conclusions for the main associations and the site-heterogeneity conclusions. For example, the ratio for active versus sedentary days changed from 2.06 to 1.88 and remained FDR-supported. Students and trainees were retained in both H10 and H06; age above 65 years was an explicit exclusion only in H10.

**Why.** H10 directly examines age associations, and H06 examines work-free versus work-day patterns alongside exercise and prior sleep. These targeted restrictions assess whether the recorded age and employment exceptions account for those findings.

**Affected hypotheses.** H10 (age and biological sex) and H06 (day type, exercise and sleep) have the targeted eligibility checks described here. The retained-cohort eligibility deviation also applies to other analyses using these participants.

**What this means for interpretation.** The restricted samples support the same substantive conclusions for H10 main effects and H06 main associations and site heterogeneity, although individual estimates change. These checks do not establish equivalence between samples, isolate an age-only exclusion effect, or verify every protocol eligibility criterion. Estimates describe the observed populations under their stated sampling rules; the targeted checks do not establish eligibility-restricted results for the other hypotheses.
